# Cyclic lipopeptide natural products as taxa-specific antibacterial inhibitors of the lipid II flippase

**DOI:** 10.1101/2025.10.02.680127

**Authors:** Elizabeth B Barrois, Jonas H Costa-Martini, Lina Thoma, Justin M Van Riper, Anne Wochele, Calla Olson, Heike Brötz-Oesterhelt, Chad W Johnston

## Abstract

Antimicrobial resistance (AMR) is an existential threat to modern healthcare; one fueled by selection pressure provided by the use of broad-spectrum antibiotics in medicine and agriculture. As these antibiotics rely on a small set of chemical scaffolds and affect an even smaller number of biological targets, emergent AMR genes can spread through microbiomes to simultaneously inactivate multiple classes and generations of drugs. Long-overlooked for their perceived clinical limitations, antibacterial natural products with taxa-specific activities now present an underexplored source of design principles for precision antibiotics that can selectively eliminate individual microbes and limit community-wide incentives for AMR. Here, we present our re-investigation of one such taxa-specific antibacterial natural product, imacidin, a forgotten inhibitor of cell wall biosynthesis. We show that imacidin is the first natural product inhibitor of the peptidoglycan lipid II flippase MurJ, representing a larger, nascent class of taxa-specific cyclic lipopeptides that offer new leads for precision antibiotics.

## Introduction

Antibiotics are essential medicines that are used to treat and prevent bacterial infections, indirectly enabling invasive surgeries, immunosuppressive therapies, and other interventions for chronic and infectious disease. Antimicrobial resistance (AMR) thus presents an existential threat to healthcare^1^, underscoring the urgent need to preserve existing antibiotics and to develop new classes of antibacterial therapeutics. In particular, it is crucial that we identify antibiotics with novel chemical scaffolds and biological targets, both to bypass existing mechanisms of AMR and limit future resistance^2,3^. The peptidoglycan cell wall remains one of the most attractive sites for the development of bactericidal agents^2^, as this conserved and essential structure is supported by an array of druggable enzymes that can be readily accessed by extracellular molecules.

Historically, leads for antibiotics have come from microbial natural products: small molecules with chemical structures and biological activities honed through evolution^4^. During the Golden Age of Antibiotic Discovery, whole cell bioactivity screens were used to look for molecules with broad activity spectra, informing our modern antibacterial arsenal^5^. During a time of slow, inaccurate diagnostics, these indiscriminate antimicrobial drugs were essential and revolutionary. While they remain clinically convenient, broad-spectrum antibiotics are now appreciated to harm beneficial microbiomes, eroding colonization resistance and incentivizing the spread of AMR that threatens the long-term viability of these drugs^3,6^. As diagnostics continue to improve, they support the development of increasingly selective ‘precision’ antibiotics^7,8^, that can selectively eliminate pathogens while sparing microbiomes and limiting community-wide incentives for AMR^9,10^.

Like the broad-spectrum molecules that came before them, leads for precision antibiotics can be found from bioactivity screens of microbial natural products^4,11^. While previous efforts biased discovery towards non-selective antimicrobials, they also uncovered rare natural products with taxa-specific antibacterial activity^7^. Historically neglected for their perceived clinical limitations, re-investigations of taxa-specific antibiotics like griselimycin (1971^12^) and nargenicin (1980^13^) revealed that these molecules bypass AMR and act on novel targets^14,15^. These studies highlight the potential of overlooked taxa-specific natural products as a source of chemical and biological leads for targeted antimicrobial therapeutics that can help turn the tide against AMR.

To uncover new foundations for precision antibiotics, we chose to investigate imacidin: a forgotten cyclic lipopeptide with taxa-specific antibacterial activity (**Fig. 1a**). Imacidin was discovered in 1979^16^ and structurally characterized in 1982^17^ as part of a prescient screening effort by Prof. Hans Zähner^18^, who was looking for narrow spectrum antibiotics that could bypass emergent AMR and spare healthy flora^19^. Isolated from *Streptomyces* sp. Tü 1379, imacidin exclusively inhibited the growth of related Actinomycetes^16^. Early mechanistic studies found that imacidin disrupts biosynthesis of the peptidoglycan sacculus, driving an accumulation of the peptidoglycan precursor Park nucleotide (UDP-N-acetylmuramic acid-pentapeptide), matching a vancomycin control^16^. As imacidin and vancomycin have profoundly different structures and activity spectra, this result suggested to us that imacidin likely affected a novel target in cell wall biosynthesis. Here, we report our re-investigation of imacidin, detailing the molecular mechanism of action (MoA) of a previously unappreciated class of taxa-specific antibacterial inhibitors of peptidoglycan biosynthesis and presenting new insights to guide the development of precision antibiotics.

**Fig. 1.**
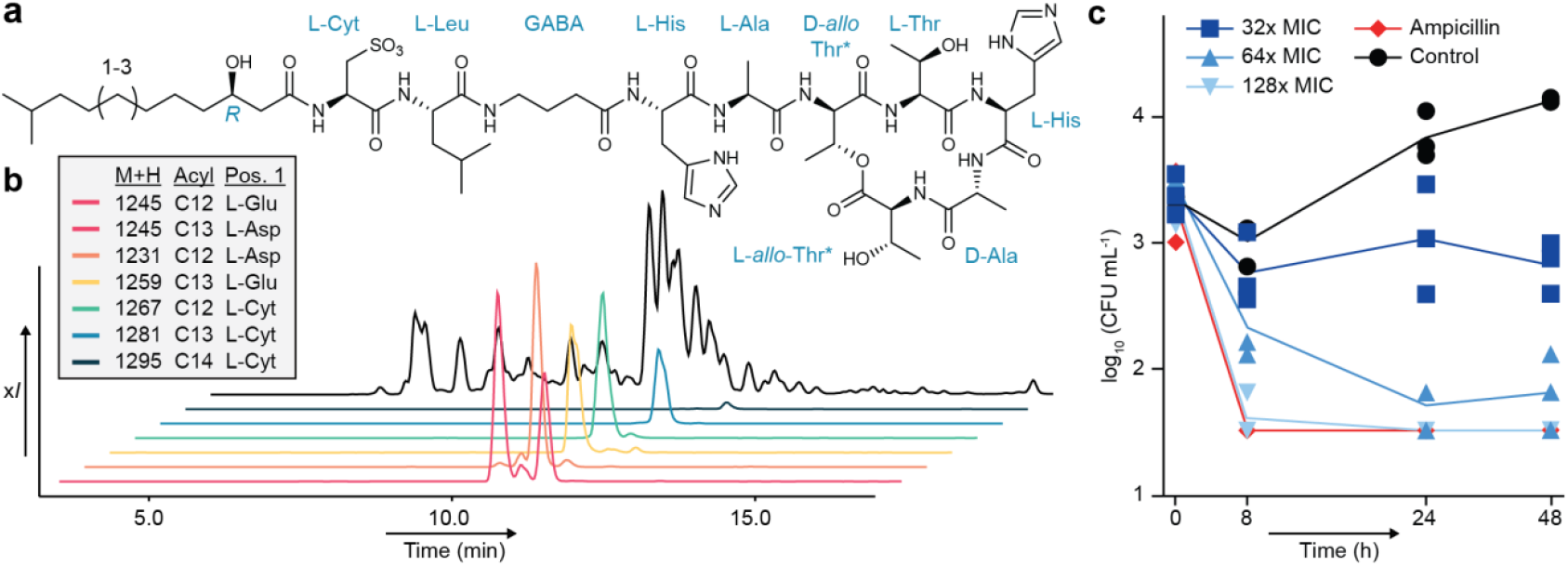
Chemical and biological confirmation of imacidin’s structure and activity. (a) The chemical structure of imacidin with “*” denoting corrections from the originally reported structure. (b) LCMS showing the separation of isolated imacidin congeners. (c) Growth curves of *S. coelicolor* M1154 were performed with increasing concentrations of imacidin and an ampicillin control (32x MIC; 1.6 mg mL^-1^). Monitoring of colony forming units (CFUs) in the presence of these antibiotics show that imacidin is bactericidal.

## Results

### Re-investigation of imacidin

To begin our re-investigation of imacidin, bacteria from the original 1979 report^16^ were revived from the strain collection at the University of Tübingen. *Streptomyces* Tü 1379 was cultured and extracted in accordance with the reported methods, recovering a series of imacidin congeners at yields between 2 and 0.002 mg L^-1^. High resolution liquid chromatography-coupled mass spectrometry (HR-LCMS) was used to isolate these congeners for structure elucidation by 2D NMR (Supplementary Information). Variations were limited to acyl tail length and substitutions in the first amino acid, with aspartate and glutamate observed in addition to the previously described cysteate (Cyt; **Fig. 1b** and Supplementary Fig. 1-3). Abundant congeners were tested in microbroth dilution assays to determine minimum inhibitory concentrations (MICs) against the model organism *Streptomyces coelicolor* M1154^20^. Longer acyl tails marginally improved activity and variants with cysteate were superior to those with glutamate and aspartate (Supplementary Table 4). Inhibition of this vancomycin-resistant strain (Supplementary Fig. 4) reinforced our belief that these molecules possessed different targets. Beyond *S. coelicolor*, we found that imacidin possessed potent, taxa-specific activity against *Streptomyces*, achieving MICs as low as 4 ng mL^-1^ (3 nM) against some species, while others were completely resistant (**Table 1**). While the original report had noted activity against *Nocardia*^16^, we did not observe this effect against a panel of isolates. Culture and 16S rRNA gene sequencing of the original test strain (denoted as *N. brasiliensis* Tü 69) indicated that this sensitive bacterium is a *Streptomyces*, further solidifying imacidin’s taxa-specificity. Next, we tested the reported bacteriostatic activity of imacidin^16^, which contradicts its proposed mode of action in disrupting cell wall biosynthesis. Monitoring colony-forming units (CFUs) from *S. coelicolor* M1154 cultures treated with imacidin or an ampicillin control confirmed that imacidin is in fact bactericidal (**Fig. 1c**). Tests against a human cell line and sheep red blood cells did not reveal cytotoxic or hemolytic activity (>64 µg mL^-1^; Supplementary Fig. 5, 6), indicating imacidin does not act on lipid membranes and likely affects a distinct target involved in peptidoglycan biosynthesis.

**Table 1.** MIC of imacidin against various bacteria.

| MIC as determined by microbroth dilution |  |
| --- | --- |
| Organism (MIC) | Imacidin ($\mu\text{g mL}^{-1}$) |
| *Escherichia coli* | >64.000 |
| *Staphylococcus aureus* ATCC 19095 | >64.000 |
| *Curtobacterium* KPL 2560 | >64.000 |
| *Dermacoccus* KPL 2534 | >64.000 |
| *Dermacoccus* KPL 2528 | >64.000 |
| *Corynebacterium striatum* KPL 1959 | >64.000 |
| *Nocardia asteroides* DSM 43132 | >64.000 |
| *Nocardia farcinica* DSM 43665 | >64.000 |
| *Nocardia brasiliensis* DSM 43758 | >64.000 |
| *Nocardia nova* DSM 44481 | >64.000 |
| *Microbispora* ATCC 55410 | 64.000 |
| *Actinomadura madurae* DSM 43067 | 10.000 |
| *Streptomyces edwardsiae* DSM 41636 | 2.500 |
| *Streptomyces europaeiscabiei* AK02-03A | 1.000 |
| *Streptomyces europaeiscabiei* AK05-03A | 0.625 |
| *Streptomyces europaeiscabiei* WA08-01A | 0.625 |
| *Streptomyces ruber* DSM 40304 | 0.100 |
| *Streptomyces rochei* ATCC 10739 | 0.080 |
| *Streptomyces bottropensis* DSM 40262 | 0.0625 |
| *Streptomyces* sp. Tü 57 | 0.040 |
| *Streptomyces europaeiscabiei* ND06-05F | 0.030 |
| *Streptomyces coelicolor* M1154 | 0.008 |
| *Streptomyces* sp. Tü 22 | 0.004 |
| MIC as determined by agar disk test |  |
| Organism (MIC) | Imacidin ($\mu\text{g mL}^{-1}$) |
| *Streptomyces* sp. Tü 6071 | >64 |
| *Streptomyces* sp. Tü 69 | 0.60 |
| *Streptomyces coelicolor* M1154 | 0.30 |
| *Streptomyces* sp. Tü 6028 | 0.30 |
| MurJ Mutants – MIC determined by microbroth dilution |  |
| Organism and MurJ mutant type | Imacidin ($\mu\text{g mL}^{-1}$) |
| *Streptomyces coelicolor* M1154 G310D | >10.0 |
| *Streptomyces coelicolor* M1154 P320H | <5.00 |
| *Streptomyces coelicolor* M1154 P491A | <5.00 |
| *Streptomyces coelicolor* M1154 Y324H | 0.10 |
| *Streptomyces coelicolor* M1154 S486Y | 0.10 |
| *Streptomyces coelicolor* M1154 P491L | 0.10 |
| *Streptomyces coelicolor* M1154 P320L | NG |
| *Streptomyces coelicolor* M1154 P491L | NG |
| NG, no growth from the isolated strain |  |
| Genetically complemented *S. coelicolor* M1154 - MIC determined by microbroth dilution |  |
| Expression vector and MurJ homolog | Imacidin ($\mu\text{g mL}^{-1}$) |
| pIJ10257 Orf6 | >10.0 |
| pIJ10257 ImcL | >10.0 |
| pIJ10257 MurJ<sup>G310D</sup> | >10.0 |
| pIJ10257 MurJ<sup>WT</sup> | 1.00 |
| pIJ10257 empty vector | 0.008 |

### Biosynthesis of imacidin

*Streptomyces* Tü 1379 is morphologically similar to the model *Streptomyces, S. coelicolor* A3(2), which produces two colored metabolites, actinorhodin and prodigiosin (Supplementary Fig. 7). As *S. coelicolor* is well-studied and profoundly imacidin-sensitive, we sequenced the genome of *Streptomyces* Tü 1379 to identify the imacidin biosynthetic gene cluster (BGC) and this bacterium’s means of self-protection. Genome sequencing with Oxford Nanopore and Illumina technologies yielded an 8.5 Mb contig with 99.99% completeness. Genome comparison between *Streptomyces* Tü 1379 and *S. coelicolor* A3(2)^21^ revealed 97.1% pairwise identity among aligned positions and 98.0% identical nucleotide sites overall (Supplementary Fig. 8). The most substantial difference was a 60 kb transposon-flanked BGC in *Streptomyces* Tü 1379, which included genes for nonribosomal peptide synthetases (NRPSs) and cysteate biosynthesis^22–24^, with NRPS adenylation domain substrate predictions^25^ largely matching imacidin (**Fig. 2a**). Two unusual features were observed in this pathway. First, the final NRPS module in ImcC lacks an A domain, with the L-*allo*-Thr substrate apparently provided *in trans* by ImcD. This arrangement was previously characterized in the BGC for WS-9326^26,27^, where a homologous stand-alone enzyme (Cal22) delivers L-*allo*-Thr *in trans*, assisted by a type II thioesterase (Cal20). A phylogenetic tree of similar NRPS enzymes collected from the MiBIG^28^ database shows that ImcD groups with other L-*allo*-Thr activating modules (Supplementary Fig. 9). A homolog of the Cal20 type II thioesterase can also be found in the imacidin BGC (ImcI; 50% identity). A second unexpected finding was the placement of the NRPS epimerization (E) domains and substrate specificities of the condensation (C) domains, which reliably indicate where D-amino acids will be incorporated into nonribosomal peptides. These suggested the originally proposed structure of imacidin had mis-assigned the positions of the D-*allo*-Thr and L-*allo*-Thr residues in the macrolactone^17^.

**Fig. 2.**
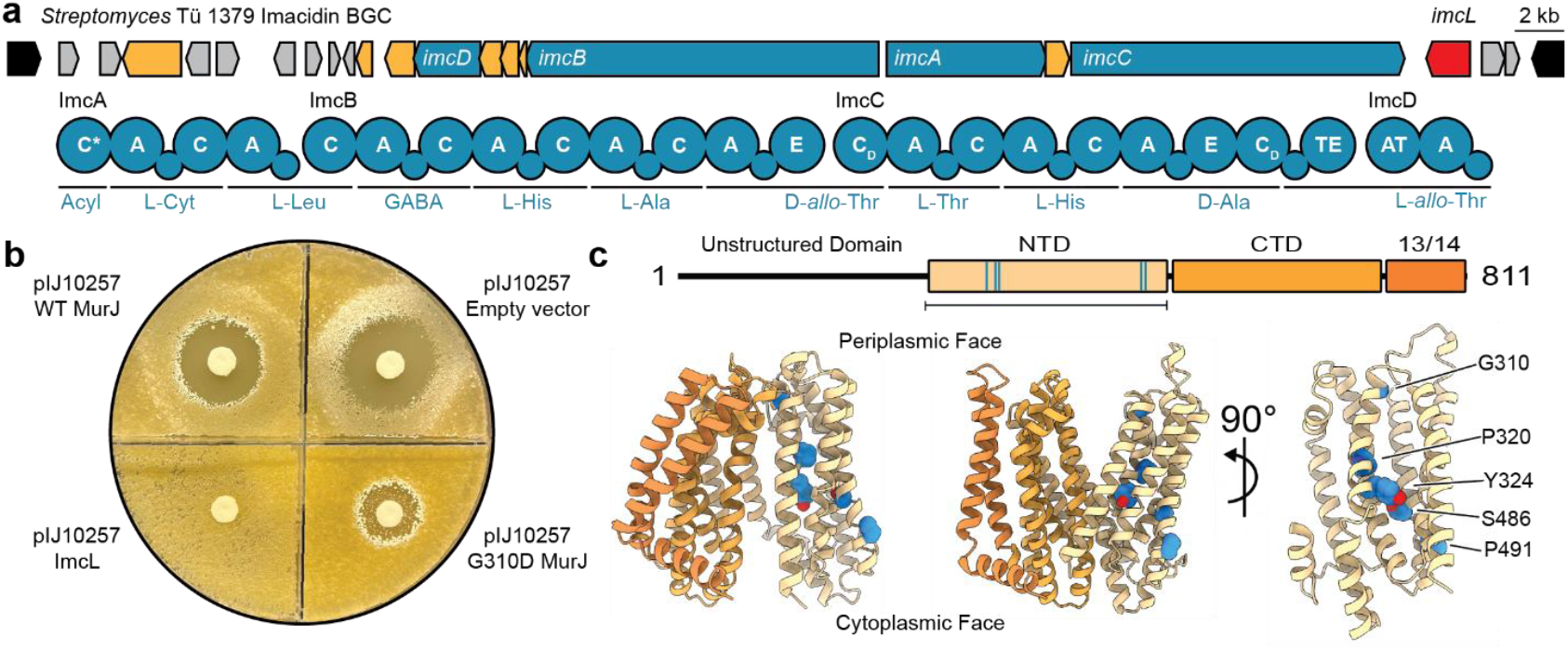
Genetics confirm MurJ as the target of imacidin. (a) On top is a layout of the imacidin BGC from *Streptomyces* Tü 1379. Below are the substrate predictions for the NRPS adenylation domains within the imacidin BGC. (b) Disk diffusion assay of imacidin against genetically complemented *S. coelicolor* M1154 expressing different MurJ homologs in the pIJ10257 vector under the *imcL* promoter. The pIJ10257 vector expresses either MurJ^WT^ (top left), an empty vector with no MurJ (top right), ImcL (bottom left), or MurJ^G310D^ (bottom right). (c) AlphaFold predicted structure of *S. coelicolor* M1154 MurJ excluding the unstructured domain. Mutations (shown as blue spheres) in imacidin-resistant mutants map to the N-terminal domain (NTD) of MurJ.

To resolve the structure of imacidin, we developed a total solid phase peptide synthesis (SPPS) scheme. Ester formation was performed in advance, using Steglich conditions to couple Boc-L-*allo*-Thr(tBu)-OH to Fmoc-D-*allo*-Thr-OH that had been loaded onto 2-chlorotrityl (2-CTC) resin. This ester linked dipeptide was eluted using hexafluoroisopropanol, preserving protecting groups and enabling direct use in downstream Fmoc-based SPPS. Synthesis of imacidin was initiated at the penultimate D-Ala residue, with rounds of deprotection and amino acid coupling building a linear peptide capped with 3-(*R*)-hydroxy-myristic acid. Trifluoroacetic acid (TFA) was used to deprotect and elute the linear peptide for purification and macrolactam formation. Purified synthetic imacidin was found to have an identical fragmentation pattern and biological activity to the natural product (Supplementary Fig. 10, 11), confirming the re-assigned structure (**Fig. 1a**).

### Genetic investigation of imacidin’s mechanism of action

The boundaries of the imacidin BGC are defined by transposons that mark its point of insertion into the genome. Along with genes for imacidin biosynthesis, the BGC encoded an additional copy of the lipid II flippase MurJ^29^ (ImcL) as an apparent resistance gene (**Fig. 2a**). As vancomycin works by binding to lipid II following its dissociation from MurJ, inhibition of this flippase by imacidin would explain phenotypic similarities^16^ following treatment with these two distinct molecules. To test whether this MurJ homolog provides imacidin resistance, we cloned the *imcL* open reading frame and promoter into the pIJ10257^30^ expression vector and transferred this into *S. coelicolor* M1154 (Supplementary Fig. 12). In contrast to an empty vector control, expression of *imcL* provided complete resistance to imacidin (**Fig. 2b**). Expression of a wildtype, codon-scrambled *S. coelicolor murJ* (SCO3894) also provided considerable resistance, which would be expected if this increased the abundance of imacidin’s target (**Fig. 2b**). To further test our hypothesis that MurJ is the target of imacidin, we used UV-irradiation to mutagenize M1154 and select for imacidin resistance, recovering seven colonies for whole genome sequencing. Each isolate carried distinct mutations in *murJ*, with some occurring in codons for the same residues. Consistent mutations were not observed in other genes (Supplementary Table 7, 8). Spontaneous resistance to imacidin was infrequent (1×10^-9^) and typically resulted in a slight decrease in sensitivity accompanied by a dramatic decrease in growth rate (Supplementary Fig. 13, 14). The sole exception to this trend was MurJ^G310D^, which demonstrated extensive imacidin resistance and grew similarly to wildtype M1154. Expression of a codon-scrambled *murJ*^G310D^ provided nearly complete imacidin resistance (**Fig. 2b**). Notably, ImcL bears a similar residue (Asn) at this position, as does the MurJ of *S. cinnamoneous*, which we previously observed to be imacidin-resistant. To assess whether this position was predictive of imacidin sensitivity, we constructed a phylogenetic tree of *Streptomyces* MurJ sequences and tested strains from the Tübingen strain library found in different clades. As expected, strains with asparagine or aspartate at this position were resistant to imacidin, while those with the more common glycine were sensitive (**Table 1**, Supplementary Fig. 15). To assess why specific sequences may be correlated with imacidin resistance, we mapped observed mutations to a model of *S. coelicolor* MurJ generated by AlphaFold^31^ (**Fig. 2c**). MurJ is comprised of 14 transmembrane (TM) helices, with TM1-6 forming the ‘N-lobe’, TM7-12 forming the ‘C-lobe’, and TM13-14 forming a hydrophobic slot that is unique to MurJ and not seen in other MOP (<u>M</u>ultidrug/<u>O</u>ligosaccharidyl-lipid/<u>P</u>olysaccharide) flippases^29^. In *Streptomyces*, a poorly conserved, unstructured intracellular domain can be found at the N-terminus. With the omission of this unstructured feature, our AlphaFold model closely resembles the ‘inward-open’ structures of MurJ from other bacteria. Resistance mutations were localized to the interior of the N-lobe, lining the central cavity used to transit lipid II. The glycine at position 310 is adjacent to a key aspartate residue that stabilizes the inward-open state in *E. coli* MurJ^32^, suggesting imacidin binds MurJ in an outward-open position and prevents the conformational change to the inward-open state.

### Phenotypic and biophysical analysis

Disturbance of peptidoglycan biosynthesis caused by inhibition of the lipid II flippase MurJ should eventually result in cell lysis. To check whether the observed bactericidal activity of imacidin (**Fig. 1c**) is due to cell lysis, we analyzed the phenotype of imacidin-treated (1-4x MIC) *S. coelicolor* M1154 by microscopy and observed swollen and lysed hyphae (**Fig. 3a**). Furthermore, at 1x MIC (8 ng mL^-1^) some hyphae showed multiple short branches suggesting disruption of polar tip extension by imacidin, with tip growth then resumed from several sites at the lateral walls (**Fig. 3a**). Time-lapse microscopy showed that these branches formed at short distance to each other from intact regions of the mycelium (Supplementary Information. Movie S1-3). These mycelial parts often lysed afterwards, suggesting that active growth is promoting cell lysis and further confirming that imacidin inhibits peptidoglycan biosynthesis. To assess the cellular localization of MurJ during vegetative growth, we expressed a fusion of the wildtype *S. coelicolor* MurJ protein with N-terminal <u>mCh</u>erry under control of the inducible P_*tipA*_ promoter^33^. Increasing concentrations of the inducer thiostrepton led to rising mCh-MurJ levels in the cells (Supplementary Fig. 16) and reduced imacidin sensitivity in disk diffusion assays (**Fig. 3b**, Supplementary Fig. 17). This correlation of mCh-MurJ expression level and imacidin sensitivity indicates that the fluorescently-labeled MurJ is functional and further supports that MurJ is the target of imacidin. We observed that mCh-MurJ specifically localized to extending tips, new branching points and presumably newly forming cross-walls which coincides with sites of peptidoglycan synthesis during vegetative growth (Supplementary Fig. 18, 19).

**Fig. 3.**
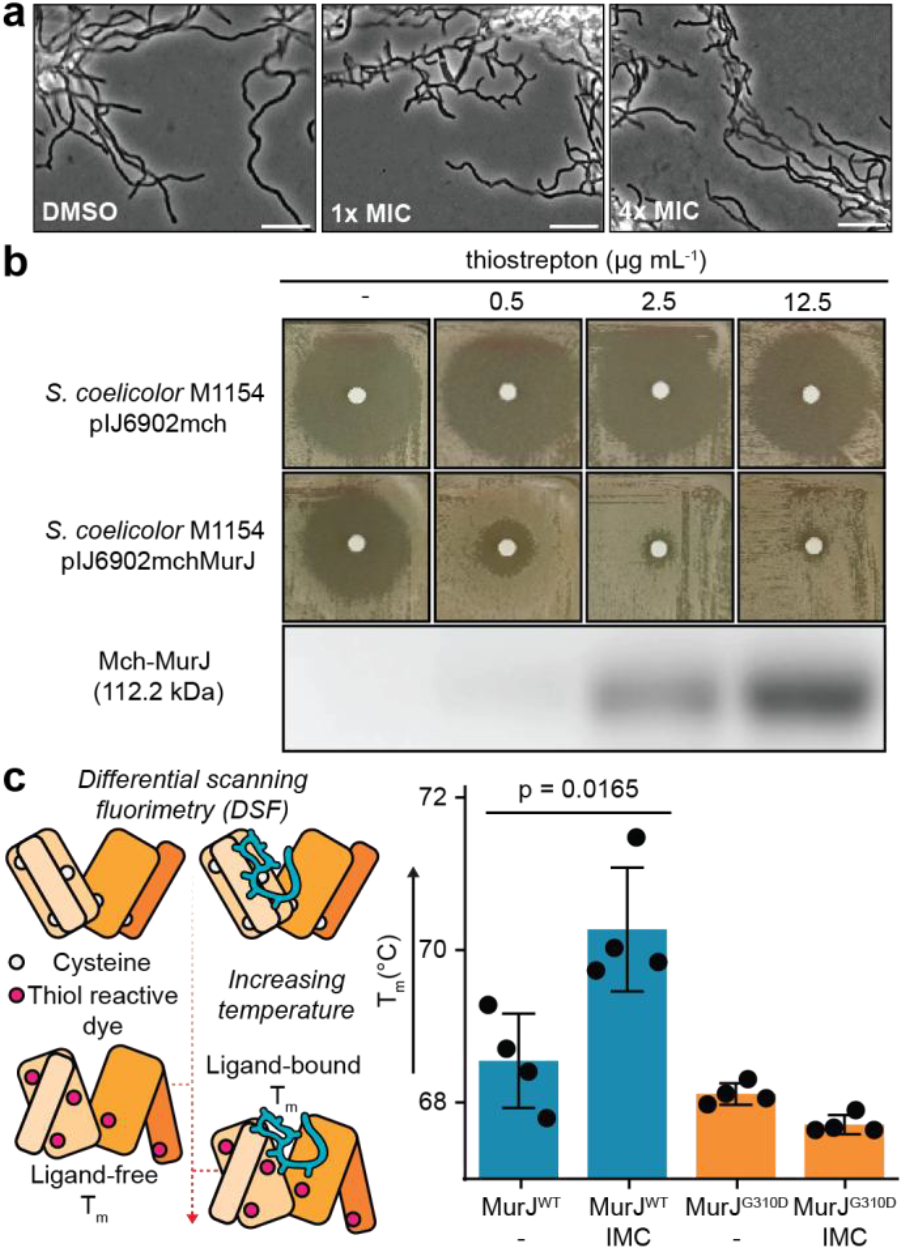
Imacidin’s antibacterial activity is dependent on MurJ expression level and genotype. (a) Imacidin-treated (1-4x MIC) *S. coelicolor* M1154 as observed by microscopy. The DMSO control shows no effect on *Streptomyces* hyphal growth. Upon the addition of imacidin (1-4x MIC), swollen and lysed hyphal branches are observed indicating bactericidal activity. (b) Increased expression of an N-terminal mCherry labeled wildtype *S. coelicolor* MurJ (mCh-MurJ) by the inducer thiostrepton shows dose-dependent imacidin resistance in disk diffusion assays. Increased expression of mCh-MurJ was confirmed by Western blot. (c) Schematic of DSF with thiol sensitive dye. When MurJ is folded, free thiols on cysteine residues are hidden in the interior of the protein. As temperature increases, MurJ unfolds revealing cysteine residues that allow the thiol reactive dye to bind and fluoresce. Ligands that directly interact with MurJ can stabilize the interaction and increase the T_m_. The graph shows the average T_m_ from 4 replicates across MurJ^WT^ and MurJ^G310D^ with and without the addition of imacidin. The addition of imacidin with MurJ^WT^ causes a significant increase in the T_m_ suggesting that imacidin directly interacts and stabilizes MurJ. The addition of imacidin with MurJ^G310D^ has no significant effect on the T_m_ indicating that the G310D substitution is likely preventing imacidin binding. Scale bar: 10 µm.

As imacidin sensitivity is dependent on both the expression level and the primary sequence of MurJ, we hypothesized that imacidin’s antibacterial activity results from a direct interaction with this protein. To test this, we endeavored to express and purify sensitive and resistant forms of *Streptomyces* MurJ for use in biophysical assays. Ancestral sequence reconstruction^34^ of imacidin sensitive MurJ proteins was used to identify candidates with optimal stability for purification from *E. coli*. Five candidates were cloned and tested for expression in *E. coli*, with stability and yield assessed by Western blot (Supplementary Fig. 20). The MurJ sequence from the exceptionally imacidin-sensitive *Streptomyces* Tü 22 performed best, with purified soluble protein obtained at yields of 0.25 mg L^-1^ (Supplementary Fig. 21). We had initially intended to probe interactions between purified MurJ and fluorescent derivatives of imacidin but we were unable to identify modifications that were functional and preserved activity (Supplementary note). Ultimately, we chose to assess potential interactions between MurJ and imacidin by differential scanning fluorimetry (DSF), alleviating the need to modify the protein or natural product. This technique measures the thermal stability of a protein by quantifying its melting temperature (T_m_), with ligand binding typically indicated by a shift to a higher T_m_ due to stabilization. As MurJ is a membrane protein, unfolding was detected using a thiol sensitive dye (BODIPY FL L-cystine^35^) to react with newly exposed cysteine residues^36^ (**Fig. 3c**). Using this experimental design, we observed a significant increase in the T_m_ of MurJ^WT^ – but not MurJ^G310D^ – in the presence of imacidin (**Fig. 3c**). These results show that imacidin directly interacts with MurJ, suggesting antibacterial activity likely stems from inhibition.

### Genome mining for additional MurJ-directed antibiotics

Given its prominent role in peptidoglycan biosynthesis and accessibility to extracellular molecules, MurJ presents an attractive target for antibacterial drug development^2,29^. To assess whether imacidin represented a larger family of natural products with this target, we performed a bioinformatic search for *Streptomyces* MurJ proteins that may act as self-protection genes within BGCs. A close homolog of ImcL (78% identity, excluding the disordered domain; Supplementary Fig. 15) was encoded in the BGC for the cyclic lipopeptide taeanamide A^37^ (**Fig. 4a, b**). Identified in 2022, taeanamide A was effectively inactive against a panel human cell lines and bacterial pathogens, with only weak activity against *Mycobacterium tuberculosis* (MIC_50_ of 32 µg mL^-1^)^37^. Given the *imcL* homolog in its BGC, we suspected that taeanamide was a second *Streptomyces*-selective, MurJ-directed natural product.

**Fig. 4.**
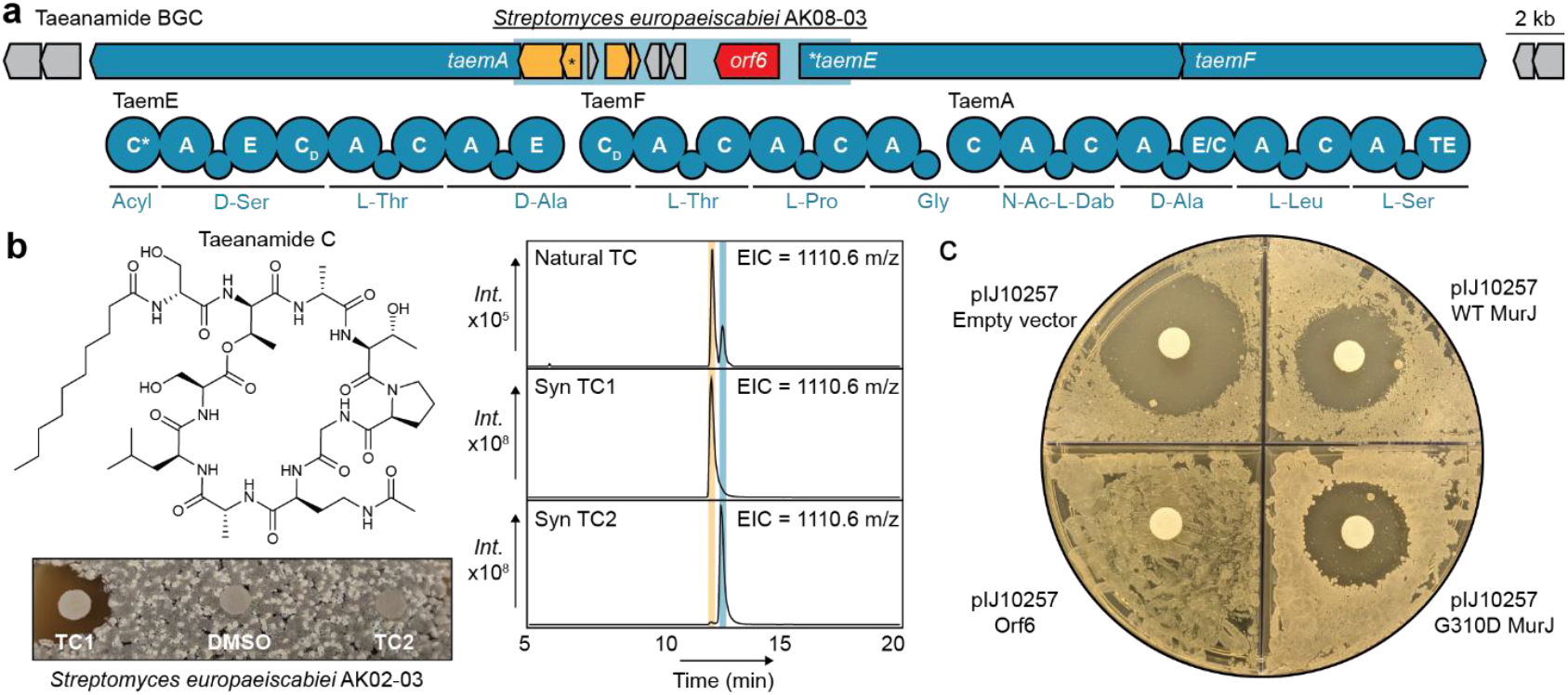
Genome mining reveals another MurJ-directed antibiotic taeanamide. (a) The partial taeanamide BGC from *Streptomyces europaeiscabiei* AK08-03 (light blue) is overlaid onto the described BGC from *S. europaeiscabiei* AMDM43, with established substrates of the NRPS modules indicated. Genes that appear to have been incorrectly sequenced in the original BGC are starred. (b) The proposed structure of taeanamide C is provided along with a disk diffusion assay of synthetic taeanamide C (TC1) and its inactive isomer (TC2) against *S. europaeiscabiei* AK02-03. LCMS retention times for natural and synthetic taeanamide C (TC) compounds are also provided (*right*). Natural TC and Syn TC compounds have identical retention times and fragmentation patterns (Supplementary Fig. 22). (c) Disk diffusion assay of imacidin against genetically complemented *S. coelicolor* M1154 expressing different MurJ homologs in the pIJ10257 vector under the *imcL* promoter. The pIJ10257 vector expresses either an empty vector with no MurJ (top left), MurJ^WT^ (top right), MurJ^G310D^ (bottom right), or Orf6 (bottom left).

Taeanamide was reported from a South Korean isolate of the plant pathogen *S. europaeiscabiei*, which complicates international shipment. Bioinformatic searches showed the partially sequenced USDA isolate *S. europaeiscabiei* AK08-03^38^ likely carried an identical BGC (**Fig. 4a**). In particular, one diagnostic contig featured the MurJ homolog *orf6*, as well as genes that were missed before, including an acylating condensation domain and an L-2,4-diamino-butyrate Nγ-acetyltransferase. Using the original media and culture conditions^37^, we grew, extracted, and fractionated cultures of AK08-03 for HR-LCMS analysis. We did not observe the reported masses for taeanamide A or the linear methyl-ester taeanamide B, but rather two isomers of a new, related molecule we term taeanamide C. Based on high resolution MS/MS, this molecule is identical to A but features a saturated C_12_ fatty acid *in lieu* of 10-methylundec-2-enoic acid (**Fig. 4b**). As taeanamide C was present in exceedingly low yield, we developed a total SPPS scheme to confirm its structure and provide material for activity testing.

Synthesis was initiated at the penultimate residue, sequentially coupling Fmoc-protected amino acids before capping the nascent peptide with lauric acid. Esterification was achieved using Steglich conditions to attach a Boc-protected serine to the appropriate threonine alcohol on the completed resin-bound lipopeptide, providing an amine for macrolactam closure following elution and deprotection with TFA. Synthetic taeanamide C was obtained as two isomers whose retention times, masses, and fragmentation patterns were identical to those seen in the natural products (**Fig. 4b**, Supplementary Fig. 22).

To assess whether taeanamide could inhibit the growth of *Streptomyces*, we tested it against a panel of strains. While the major taeanamide C isomer was inactive against *S. coelicolor*, it inhibited the growth of *Streptomyces* related to *S. europaeiscabiei* (**Fig. 4b**; **Table 2**; MIC: 1 to 5 µg mL^-1^), further suggesting *orf6* is used for self-protection. Taeanamide-sensitive strains were also sensitive to imacidin (**Table 1**; MIC: 0.025 to 1.25 µg mL^-1^), suggesting nested activity spectra between these two natural products. To test if imacidin and taeanamide shared a common target, we returned to the genetically amenable *S. coelicolor* M1154, which was transformed with pIJ10257 expressing Orf6 or previously tested MurJ sequences (**Fig. 2b**). Like ImcL before, Orf6 expression provided total resistance to imacidin (**Fig. 4c**), indicating taeanamide likely also acts as an inhibitor of *Streptomyces* MurJ and suggesting these molecules may represent a larger family of MurJ-directed natural products.

**Table 2.**
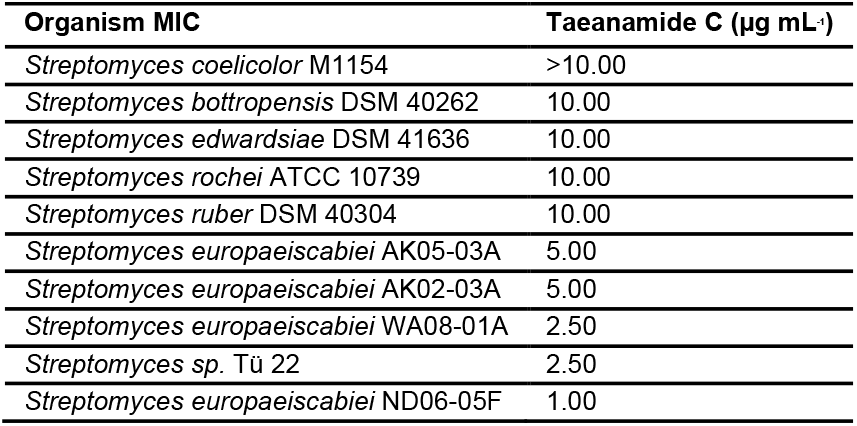
MIC of taeanamide C against *Streptomyces*.

## Discussion

Natural products with taxa-specific antibacterial activities are an overlooked resource for antibiotic development, offering evolved chemical scaffolds and target interactions that can bypass clinical resistance to selectively eliminate individual families of microbes. To provide further insights into the mechanisms of taxa-specific antibiotics, we re-investigated the forgotten, *Streptomyces*-selective molecule imacidin. Our results indicate that imacidin is the first natural product inhibitor of the lipid II flippase MurJ^29^. While we lack a means of directly quantifying the inhibition of MurJ in *Streptomyces*, several lines of evidence support this conclusion. Both the imacidin and taeanamide BGCs contain homologs of *murJ* that provide antibiotic resistance (**Fig. 2a, b, Fig. 4a, c**). Spontaneous imacidin resistance was associated with mutations in *murJ* (**Fig. 2c**). Expression of MurJ in sensitive *Streptomyces* provided dose-dependent resistance to imacidin (**Fig. 3b**). Imacidin-treated cells displayed phenotypes consistent with disrupted peptidoglycan biosynthesis (**Fig. 3a**; Supplementary Movie 1-3). Finally, we found that wildtype – but not mutant – MurJ was thermally stabilized by imacidin, indicating direct binding (**Fig. 3c**).

While inhibitors of MurJ have been identified^29^, such molecules remain rare and relatively poorly characterized. Direct evidence of MurJ inhibition has only been reported for the *E. coli* phage lysis protein LysM, first by an *in vivo* flippase assay in *E. coli*^39^, and recently by solving the structure of the complex^40^ by single particle cryogenic electron microscopy. Putative synthetic small molecule inhibitors of MurJ were recovered from chemical screens^41,42^ that predated its description as the lipid II flippase^43^, with this target proposed on the basis of resistance mapping, genetic complementation, and phenotypic analysis. Later, the biologically-inspired, linear lipopeptide humimycin^44,45^ was also proposed as a MurJ inhibitor. In addition to non-specific hemolytic activity, humimycin treatment of *Staphylococcus* led to resistance mutations in *murJ* and sensitized cells to β-lactam antibiotics^44^, as seen with the earlier synthetic compounds^41,42^. Against the most sensitive strains, these synthetic inhibitors displayed antibacterial MICs between 0.12 and 4 µM^41^.

Imacidin has the most potent antibacterial activity among MurJ inhibitors identified thus far (Table 1), with MICs as low as 3 nM, reflecting the refined nature of this interaction. Further, as imacidin is a natural product, bioinformatic searches using the BGC-associated self-protection gene can now be used to identify other MurJ-inhibitors, irrespective of their structural similarity. The example of taeanamide indicates the existence of larger family of lipopeptide natural products that act as taxa-specific inhibitors of MurJ. This specificity contrasts with the many known broad-spectrum compounds that disrupt cell wall biosynthesis and may explain why MurJ inhibitory natural products were not found in earlier bioactivity-guided screening campaigns. As MurJ inhibition is bactericidal and also sensitizes bacteria to clinically essential β-lactams, these potent, non-toxic^37^ (Supplementary Fig. S5), and synthetically accessible cyclic lipopeptides present a fascinating new scaffold for antibiotic development.

Despite its specificity, mutations that provided spontaneous resistance to imacidin were rare (1×10^-9^) and typically resulted in a modest increase in MIC at a considerable cost to growth rate. The sole exception we identified was the G310D substitution, which grows similarly to wildtype, provides considerable resistance, and mirrors the sequence of resistant MurJ-homologs encoded in cyclic lipopeptide BGCs. Given its position adjacent to the extracellular gate^32^, we speculate that this change stabilizes the inward-open conformation to limit antibiotic access to the central cavity. Across tested *Streptomyces*, the identity of this position was predictive of inhibitor sensitivity, cleanly dividing MurJ proteins into sensitive or resistant clades (Supplementary Fig. 15). Interestingly, the imacidin and taeanamide BGCs were acquired by ‘sensitive’ strains through horizontal gene transfer, presenting a means of antibiotic production and self-protection that allowed them to antagonize close relatives. While antagonism between related bacteria is a common phenomenon^7,46,47^, the selectivity of these inhibitors is still surprising. Both lipid II and MurJ’s central cavity are conserved between bacteria^32^, and prior synthetic inhibitors were generally active against Gram-positive bacteria and efflux-deficient Gram-negatives. Moving forward, we will investigate the interactions that support taxa-specific MurJ inhibition to inform the design of new antibiotics with customized activity spectra.

## Acknowledgements

The authors wish to thank E. Draskoczy (University of Tübingen) for her contributions to the microscopy work; Prof. K. MacKenzie (Baylor College of Medicine; BCM) for assistance with NMR data collection and analysis; Prof. M. Zhou (BCM) for his kind guidance, mentorships, and assistance with membrane protein purification; Prof. G. Wright (McMaster University) and the scientists of the John Innes Centre (Norwich, UK) for providing *S. coelicolor* M1154, pIJ10257, and *E. coli* ET12567 pUZ8002; Dr. C. Clarke (USDA) for providing strains of *S. europaeiscabiei;* Prof. K. Lemon (BCM) for providing *Corynebacterium, Curtobacterium*, and *Dermacoccus* strains; and the Alkek Center for Metagenomics and Microbiome Research (BCM) for Illumina sequencing and data processing. This work was funded by the National Institutes of Health (NIH; NIGMS; 1R21GM154190) and by the Cancer Research and Prevention Institute of Texas (CPRIT; RR210066). C.W.J. is a CPRIT Scholar in Cancer Research. H.B.O and L.T. were funded by the Cluster of Excellence EXC2124 project ID 390838134.

## Notes

### Competing Interest Statement

The authors have declared no competing interest.

### Summary of Updates

The revised manuscript and supplementary information corrects several minor typos in the main text, supplementary tables, and figure legends. Author order has also been slightly changed.

## References

1. Hobson, C., Chan, A. N. & Wright, G. D. The Antibiotic Resistome: A Guide for the Discovery of Natural Products as Antimicrobial Agents. Chem. Rev. 121, 3464–3494 (2021).

2. Theuretzbacher, U., Blasco, B., Duffey, M. & Piddock, L. J. V. Unrealized targets in the discovery of antibiotics for Gram-negative bacterial infections. Nat Rev Drug Discov 22, 957– 975 (2023).

3. Cook, M. A. & Wright, G. D. The past, present, and future of antibiotics. Sci. Transl. Med. 14, eabo7793 (2022).

4. Lewis, K. et al. Sophisticated natural products as antibiotics. Nature 632, 39–49 (2024).

5. Lewis, K. The Science of Antibiotic Discovery. Cell 181, 29–45 (2020).

6. Fishbein, S. R. S., Mahmud, B. & Dantas, G. Antibiotic perturbations to the gut microbiome. Nat Rev Microbiol 21, 772–788 (2023).

7. Johnston, C. W. & Badran, A. H. Natural and engineered precision antibiotics in the context of resistance. Current Opinion in Chemical Biology 69, 102160 (2022).

8. Maxson, T. & Mitchell, D. A. Targeted treatment for bacterial infections: prospects for pathogen-specific antibiotics coupled with rapid diagnostics. Tetrahedron 72, 3609–3624 (2016).

9. Muñoz, K. A. et al. A Gram-negative-selective antibiotic that spares the gut microbiome. Nature 630, 429–436 (2024).

10. Zampaloni, C. et al. A novel antibiotic class targeting the lipopolysaccharide transporter. Nature 625, 566–571 (2024).

11. Wright, G. D. Opportunities for natural products in 21 ^st^ century antibiotic discovery. Nat. Prod. Rep. 34, 694–701 (2017).

12. Terlain, B. & Thomas, J. P. [Structure of griselimycin, polypeptide antibiotic extracted from streptomyces cultures. II. Structure of griselimycin]. Bull Soc Chim Fr 6, 2357–2362 (1971).

13. Celmer, W. D. et al. Structure of natural antibiotic CP-47,444. J. Am. Chem. Soc. 102, 4203– 4209 (1980).

14. Kling, A. et al. Targeting DnaN for tuberculosis therapy using novel griselimycins. Science 348, 1106–1112 (2015).

15. Painter, R. E. et al. Elucidation of DnaE as the Antibacterial Target of the Natural Product, Nargenicin. Chemistry & Biology 22, 1362–1373 (2015).

16. Brecht-Fischer, A., Zhner, H. & Laatsch, H. Stoffwechselprodukte von Mikroorganismen 183. Mitteilung. Imacidin, ein neues Acylpeptidantibioticum aus Streptomyces olivaceus. Arch. Microbiol. 122, 219–229 (1979).

17. Laatsch, H. Stoffwechselprodukte von Mikroorganismen, 204 ^1^) Die Struktur von Imacidin C. Liebigs Ann. Chem. 1982, 28–40 (1982).

18. Bioactive Microbial Products : Search and Discovery /. (Published for the Society for General Microbiology by Academic Press, London;, 1982).

19. Zähner, H. & Maas, W. K. The Future of Antibiotics. in Biology of Antibiotics 114–122 (Springer US, New York, NY, 1972). doi:10.1007/978-1-4613-9373-3_7.

20. Gomez‐Escribano, J. P. & Bibb, M. J. Engineering Streptomyces coelicolor for heterologous expression of secondary metabolite gene clusters. Microbial Biotechnology 4, 207–215 (2011).

21. Bentley, S. D. et al. Complete genome sequence of the model actinomycete Streptomyces coelicolor A 3(2). (2002).

22. Bekiesch, P. et al. Viennamycins: Lipopeptides Produced by a Streptomyces sp. J. Nat. Prod. 83, 2381–2389 (2020).

23. Liu, W.-T. et al. MS/MS-based networking and peptidogenomics guided genome mining revealed the stenothricin gene cluster in Streptomyces roseosporus. J Antibiot 67, 99–104 (2014).

24. Takeda, K. et al. N -Phenylacetylation and Nonribosomal Peptide Synthetases with Substrate Promiscuity for Biosynthesis of Heptapeptide Variants, JBIR-78 and JBIR-95. ACS Chem. Biol. 12, 1813–1819 (2017).

25. Blin, K. et al. antiSMASH 8.0: extended gene cluster detection capabilities and analyses of chemistry, enzymology, and regulation. Nucleic Acids Research 53, W32–W38 (2025).

26. Johnston, C. W. et al. An automated Genomes-to-Natural Products platform (GNP) for the discovery of modular natural products. Nat Commun 6, 8421 (2015).

27. Kim, M. et al. Unprecedented Noncanonical Features of the Nonlinear Nonribosomal Peptide Synthetase Assembly Line for WS9326A Biosynthesis. Angew Chem Int Ed 60, 19766–19773 (2021).

28. Zdouc, M. M. et al. MIBiG 4.0: advancing biosynthetic gene cluster curation through global collaboration. Nucleic Acids Research 53, D678–D690 (2025).

29. Kuk, A. C. Y., Hao, A. & Lee, S.-Y. Structure and Mechanism of the Lipid Flippase MurJ. Annu Rev Biochem 91, 705–729 (2022).

30. Hong, H.-J., Hutchings, M. I., Hill, L. M. & Buttner, M. J. The Role of the Novel Fem Protein VanK in Vancomycin Resistance in Streptomyces coelicolor. Journal of Biological Chemistry 280, 13055–13061 (2005).

31. Abramson, J. et al. Accurate structure prediction of biomolecular interactions with AlphaFold 3. Nature 630, 493–500 (2024).

32. Zheng, S. et al. Structure and mutagenic analysis of the lipid II flippase MurJ from Escherichia coli. Proc. Natl. Acad. Sci. U.S.A. 115, 6709–6714 (2018).

33. Takano, E., White, J., Thompson, C. J. & Bibb, M. J. Construction of thiostrepton-inducible, high-copy-number expression vectors for use in Streptomyces spp. Gene 166, 133–137 (1995).

34. Spence, M. A., Kaczmarski, J. A., Saunders, J. W. & Jackson, C. J. Ancestral sequence reconstruction for protein engineers. Current Opinion in Structural Biology 69, 131–141 (2021).

35. Hofmann, L., Gulati, S., Sears, A., Stewart, P. L. & Palczewski, K. An effective thiol-reactive probe for differential scanning fluorimetry with a standard real-time polymerase chain reaction device. Analytical Biochemistry 499, 63–65 (2016).

36. Helbling, R. E., Aeschimann, W., Simona, F., Stocker, A. & Cascella, M. Engineering Tocopherol Selectivity in α-TTP: A Combined In Vitro/In Silico Study. PLoS ONE 7, e49195 (2012).

37. Cui, J. et al. Taeanamides A and B, Nonribosomal Lipo-Decapeptides Isolated from an Intertidal-Mudflat-Derived Streptomyces sp. Marine Drugs 20, 400 (2022).

38. Weisberg, A. J., Pearce, E., Kramer, C. G., Chang, J. H. & Clarke, C. R. Diverse mobile genetic elements shaped the evolution of Streptomyces virulence. Microbial Genomics 9, (2023).

39. Chamakura, K. R. et al. A viral protein antibiotic inhibits lipid II flippase activity. Nat Microbiol 2, 1480–1484 (2017).

40. Kohga, H. et al. Phage lysis protein Lys^M^ acts as a wedge to block MurJ conformational changes. Preprint at 10.1101/2025.04.26.650401 (2025).

41. Huber, J. et al. Chemical Genetic Identification of Peptidoglycan Inhibitors Potentiating Carbapenem Activity against Methicillin-Resistant Staphylococcus aureus. Chemistry & Biology 16, 837–848 (2009).

42. Mott, J. E. et al. Resistance mapping and mode of action of a novel class of antibacterial anthranilic acids: evidence for disruption of cell wall biosynthesis. Journal of Antimicrobial Chemotherapy 62, 720–729 (2008).

43. Sham, L.-T. et al. MurJ is the flippase of lipid-linked precursors for peptidoglycan biogenesis. Science 345, 220–222 (2014).

44. Chu, J. et al. Discovery of MRSA active antibiotics using primary sequence from the human microbiome. Nat Chem Biol 12, 1004–1006 (2016).

45. Rahman, M. R. T., Guay, L.-D., Fliss, I. & Biron, E. Structure–Activity Study of the Antimicrobial Lipopeptide Humimycin A and Screening Against Multidrug-Resistant Staphylococcus aureus. Antibiotics 14, 385 (2025).

46. Heilbronner, S., Krismer, B., Brötz-Oesterhelt, H. & Peschel, A. The microbiome-shaping roles of bacteriocins. Nat Rev Microbiol 19, 726–739 (2021).

47. Peterson, S. B., Bertolli, S. K. & Mougous, J. D. The Central Role of Interbacterial Antagonism in Bacterial Life. Current Biology 30, R1203–R1214 (2020).

