## Supplementary Information for "Cyclic lipopeptide natural products as taxa-specific antibacterial inhibitors of the lipid II flippase"

**Table of Contents**

| | |
|---|---|
| Materials and methods | 2 – 10 |
| Supplemental tables | 11 – 17 |
| Supplemental figures | 18 – 32 |
| Supplemental movies | 33 |
| NMR spectra and tables | 34 – 52 |
| Supplemental notes | 53 – 54 |
| Supplemental references | 55 |

### Materials and methods

#### Bacterial strains and growth conditions

Strains are listed in Table S1. *Escherichia coli*, *Curtobacterium*, *Corynebacterium*, *Dermacoccus,* and *Staphylococcus* strains were cultured in Luria Bertani (LB) broth (BD DIFCO™, Miller) or LB agar at 30-37°C. When antibiotic selection was necessary, media were supplemented with apramycin (100 μg $mL^{-1}$), kanamycin (25-50 μg $mL^{-1}$), ampicillin (100 μg $mL^{-1}$), or hygromycin b (100 μg $mL^{-1}$). *Streptomyces* strains were cultivated at 30 °C in LB broth or LB agar, GYM medium (glucose 4 g $L^{-1}$, yeast extract 4 g $L^{-1}$, malt extract 10 g $L^{-1}$, pH 7.2), GYM agar (glucose 4 g $L^{-1}$, yeast extract 4g $L^{-1}$, malt extract 10 g $L^{-1}$, $CaCO_3$ 2 g $L^{-1}$, 20 g agar, pH 7.2) or NMMP media ($(NH_4)_2SO_4$ 3 g $L^{-1}$, Difco casaminoacids 5 g $L^{-1}$, $MgSO_4 \cdot 7H_2O$ 0.6 g $L^{-1}$, $NaH_2PO_4 \cdot 1H_2O$ 2.07 g $L^{-1}$, $K_2HPO_4$ 2.6 g $L^{-1}$, PEG 6000 [60% PEG 8000 & 40% PEG 3350] 50 g $L^{-1}$, mannitol 10 g $L^{-1}$, 1 mL minor elements solution pH 6.8 [$ZnSO_4 \cdot 7H_2O$ 1 mg $mL^{-1}$, $FeSO_4 \cdot 7H_2O$ 1 mg $mL^{-1}$, $MnCl_2 \cdot 4H_2O$ 1 mg $mL^{-1}$, anhydrous $CaCl_2$ 1 mg $mL^{-1}$]). When necessary, media was supplemented with apramycin (50 μg $mL^{-1}$) or (nalidixic acid 25 μg $mL^{-1}$). In order to obtain *Streptomyces* spores, mycelia or spores were streaked on MS agar (soya flour 20 g $L^{-1}$, mannitol 20 g $L^{-1}$, 16 g agar $L^{-1}$) and incubated for 7-8 days at 30 °C. Plasmid-harboring *Streptomyces* strains were streaked on MS agar supplemented with apramycin (50 μg $mL^{-1}$) or hygromycin b (100 μg $mL^{-1}$) to ensure plasmid maintenance during sporulation.

#### Minimum inhibitory concentration (MIC) assays

Microbroth dilution was used to determine MICs. The spores from various *Streptomyces* were collected and stocked[1]. Each *Streptomyces* spore stock was serially diluted, plated onto GYM agar, and incubated for 2 days at 30 °C to determine colony forming units (CFUs). For the MICs, spores were diluted to $1.6 \times 10^6$ CFU $mL^{-1}$ and inoculated into 96 deep well plates (Greiner) containing imacidin or taeanamide solutions serially diluted twofold in either GYM media or NMMP media. MIC plates were cultured at 30°C for 48 hours with 85% humidity and shaking at 900 rpm. Non-*Streptomyces* bacteria were grown to an $OD_{600}$ of 0.1 and inoculated into 96 well plates (Greiner) containing Mueller Hinton broth (Criterion™). MIC plates were cultured overnight at 37°C with 85% humidity and shaking at 900 rpm. The imacidin and taeanamide MICs were determined as the minimum concentration where no growth could be detected by the eye. All MIC assays were performed in triplicate.

#### Cell culture

hTERT-HME1 (Infinity; female) cells were cultured in MEGM (Lonza, CC-3150) and incubated at 37°C and 5% $CO_2$. The cell line was obtained from ATCC and tested yearly for mycoplasma contamination.

#### Time-dependent killing curves

35 μL of a $10^8$ spores $mL^{-1}$ solution of *S. coelicolor* M1154 were diluted in 4 mL of NMMP media containing small glass beads and incubated overnight at 30°C shaking at 220 rpm[2]. *S. coelicolor* was treated with 32x, 64x and 128x MIC imacidin at time point 0 hours. Ampicillin (32x MIC) was used as positive control. At each collection time point, 100 μL aliquots were collected and centrifuged at 15,000 rpm for 10 min. The cell pellet was washed with 100 μL of PBS and resuspended in 100 μL of PBS. Tenfold serially dilutions were plated over GYM agar plates that were then cultured for 24 hours at 30°C. Colonies of *S. coelicolor* were counted to calculate CFU $mL^{-1}$. Time-dependent killing curve experiments were performed in triplicate.

#### Cytotoxicity assay

hTERT-HME1 cells were seeded at a density of 5,000 onto a 96 well black clear bottom plate (Corning). After seeding, cells were treated with a dilution series (0 μM to 40 μM) of puromycin

(Sigma-Aldrich) or imacidin. Methanol was used as a vehicle control. Cells were incubated for 48 hours at 37°C and 5% $CO_2$. Cytotoxicity was determined by the raw cell count from brightfield imaging using the Celigo imaging cell cytometer (Brooks).

**Red blood cell (RBC) hemolysis assay**
Imacidin's hemolytic activity was determined using an adapted published protocol[3]. Imacidin was serially diluted (64 µg $mL^{-1}$ to 1 µg $mL^{-1}$) and incubated for 1 hour at 37°C with 0.25% sheep red blood cell in phosphate buffered saline. RBC alone and 0.5% DMSO were used as negative controls. 1% Triton X was used as a positive control. After incubation, the RBC were centrifuged at 1,000*g* for 5 min. The supernatant was transferred to a flat-bottom 96 well polystyrene plate (Greiner) and absorbance was measured at 540 nm.

***De novo* assembly of the *Streptomyces* Tü 1379 producer strain**
The Monarch Genomic DNA Purification Kit (#T3010S) was used to extract genomic DNA (gDNA) from *Streptomyces* Tü 1379. The gDNA was sequenced by long reads using the Oxford Nanopore MinION Mk1B system and by short reads using Illumina Next-Generation Sequencing. For quality control, long reads were checked by chopper v0.9[4] using a minimum quality of 10 and a minimum length of 100. Short read quality control was assessed using fastp v0.23.4[5,6] in its default settings. Reads were assembled using the de novo long read assembler Trycycler v0.5.5[7] followed by short read polishing with Polypolish v0.6.[8] Final contig quality was assessed using CheckM2 v1.0.2.[9]

**Frequency of resistance**
*S. coelicolor* M1154 spores were plated at a concentration of $1x10^9$ CFU $mL^{-1}$ onto GYM agar plates containing 5 µg $mL^{-1}$ of imacidin. Plates were incubated at 30°C for 48 hours. Resistant colonies were transferred to liquid cultures containing 1 µg $mL^{-1}$ imacidin to ensure colonies were resistant. Plating of spores for resistance was done in triplicate.

**Identification of single-nucleotide mutations in imacidin-resistant *S. coelicolor* M1154**
The Monarch Genomic DNA Purification Kit (#T3010S) was used to extract genomic DNA (gDNA) from wildtype (WT) and imacidin-resistant *S. coelicolor* M1154 colonies. Imacidin-resistant colonies were obtained by plating *S. coelicolor* M1154 spores onto plates at $1x10^9$ CFU $mL^{-1}$ and exposing to UV light for 30 seconds. Illumina Next-Generation Sequencing was used to obtain genomic sequence data. Quality control was assessed using FastQC v0.11.9. The sequencing reads were paired and trimmed using BBDuk on Geneious Prime software. To identify single-nucleotide mutations, we used the computational pipeline BRESEQ v0.38.1.[10]

**Structural modeling of MurJ**
The structural predictions of MurJ from (species) were generated using AlphaFold. The inward open state was predicted from the amino acid sequence (274-811) using AlphaFold 3[11]. The outward open state was generated by AlphaFold 2 from the same amino acid and run on ColabFold[12]. The prediction parameters on ColabFold were adjusted to utilize the structure of the outward-facing conformation of Lipid II flippase MurJ from *Thermosipho africanus* TCF52B (PDB: 6NC9) as a template. Additionally, the MSA_mode was set to single_sequence and resulted in a confident prediction (pLDDT=76.1 pTM=0.791).

Sequence used:
GLLKSSAVMAAGTMVSRLTGFVRSALIVSALGVGLLGDTFQVAYQLPTMIYILTVGGGLNSVFV
PQLVRAMKDDEDGGEAFANRLLTLVMVALGALTVITVFAAPLLIRLLSNPVASDPAANEVGITFV
RYFLPSIFFMGLHVVMGQVLNARGRFGAMMWTPVLNNIVIIVTLGLFIWVYGTAETSGMKVTSIP
PEGERLLGIGVLLGLIVQSLAMIPYLRETGFRLRLRFDWRGHGLGKAITLAKWTVLFVLANQAGA

MIVIQLSTAAGKASPVDGTGFAAYANAQLIWGLPQAIITVSLMAALLPRISRSASEEDGGAVRDDI
SQGLRTTAVAIVPVSFGFVALGIPMCTLMFGSSGTSEATNMGYMLMAFGLGLIPYSVQYVVLRA
FYAYEDTRTPFYNTVIVAVVNAAASGLCYLLLPSRWAVVGMAASYGLAYVIGVGIAWRRLRKRL
GGDLDGARVLRTYARLCIASVPAALIGGAACYAISRSLGQGVVGSLAALLAGGVLLFGVFFVAA
RRMRIEEVNSLVGMVRGRLGR

**Genetic complementation of *Streptomyces coelicolor* M1154**

Generation and transformation of *Streptomyces* protoplasts and the genetic complementation of plasmid DNA into *S. coelicolor* M1154 was completed using previously established protocols[1]. All strains, primers, and plasmids used for genetic complementation are listed in Tables 1-3. USER® Cloning (New England Biolabs) was used for all plasmid construction. Early attempts to obtain *Streptomyces* transformants with pIJ10257 were unsuccessful due to possible toxicity from the overexpression of MurJ, so we exchanged the constitutive *ermE** promoter for the native MurJ promoter using primers CJ_386 and CJ_387. To construct pIJ10257-MurJ$^{WT}$, wildtype MurJ was amplified by PCR from a codon-optimized TWIST gene using primers CJ_584 and CJ_585 and cloned into pIJ10257 using primers CJ_611 and CJ_612. To construct pIJ10257-MurJ$^{imcL}$ , MurJ from the imacidin BGC was amplified by PCR from *Streptomyces* Tü 1379 gDNA using primers CJ_366 and CJ_388 and cloned into pIJ10257 using primers CJ_367 and CJ_389. To construct pIJ10257-MurJ$^{G310D}$, primers CJ_613 and CJ_614 were phosphorylated and used in a blunt ligation to introduce the G310D mutation. To construct pIJ10257-MurJ$^{orf6}$ primers CJ_1035 and CJ_1036 were used to amplify the pIJ10257 backbone by PCR. Primers CJ_1033 and CJ_1034 were used to amplify MurJ from *S. europaeiscabiei* AK08-03 gDNA and cloned into pIJ10257. Correct cloning was confirmed by Sanger sequencing. All constructs were introduced into *S. coelicolor* M1154 using protoplasts. Transformed protoplasts were plated onto R2YE (sucrose 103 g $L^{-1}$, $K_2SO_4$ 0.25 g $L^{-1}$, $MgCl_2.6H_2O$ 10.12 g $L^{-1}$, glucose 10 g $L^{-1}$, Difco casaminoacids 0.1 g $L^{-1}$; 100 mL of this solution was combined with 2.2 g of agar and autoclaved, following the autoclave, these solutions were added in order: $KH_2PO_4$ (0.5%) 1 mL, $CaCl_2.2H_2O$ (5M) 0.4 mL, L-proline (20%) 1.5 mL, NaOH (1N) 0.7 mL) media and grown overnight. The next day plates were overlayed with 1 mL of hygromycin B (100 µg) to select for transformed colonies. After 3 days transformants were streaked onto MS agar plates and harvested after 7 days of sporulation.

**Disk diffusion assay**

To probe for susceptibility of strains from the Tübingen strain library to imacidin, disk diffusion assays were performed by plating a spore suspension containing $5x10^6$ CFUs of the respective *Streptomyces* strain on a squared format petri dish with GYM agar. Imacidin was diluted in DMSO and 10 µl of the respective dilutions were spotted on filter disks. DMSO and vancomycin (30 µg, Oxoid™) were used as controls. Filter disks were placed on the agar plates and incubated for 45-51 hours at 30°C, before zones of growth inhibition were recorded. When thiostrepton was used to induce expression of mCherry (mCh) or mCh-MurJ from the *tipA* promoter in plasmid pIJ6902, thiostrepton was added to the GYM agar.

**Detection of mCherry/mCherry-MurJ by Western blot**

A sheet of autoclaved cellophane was placed on a GYM agar plate (with or without thiostrepton) and inoculated with 100 µl of a spore suspension containing approx. $5x10^6$ CFU $mL^{-1}$. After 24 hours of growth at 30°C, the mycelium was scraped off the cellophane. 30 µl PBS buffer containing complete EDTA protease inhibitor (Roche) was added per milligram cell mass. Cells were disrupted using glass beads (0.5 mm glass beads, Roth) in a homogenizer (Precellys). Then reducing sample buffer (Bolt™, LDS sample buffer + reducing agent) (Invitrogen) was added. Samples were heated for 10 min at 98°C and run on a Bolt™ 12%, Bis Tris Plus WedgeWell™ Gel (Invitrogen) in Bolt™ MOPS SDS running buffer (Invitrogen). Proteins were semi-dry transferred to a PVDF membrane. For the Western blot, a mouse monoclonal antibody detecting mCherry (Abcam, ab125096, 1:2000) was used as a primary antibody and a Goat Anti-Mouse IgG Fc (HRP) (Abcam, ab97265, 1:10,000) was used as a secondary antibody. Finally, the HPR activity was detected using the ECL™ Prime Western Blotting Detection reagent (Amersham™) and charge-coupled devices (CCDs) imaging (Chemidoc™ MP Biorad). As a loading control, a second PAGE was run with same samples under the same conditions and stained using a single-step Coomassie Blue protein gel dye (Blauer Jonas, German Research Products).

**Isolation of total DNA of actinomycetes strains**
For isolation of total DNA, 10 mL of LB broth were inoculated with 10 µl of a dense spore suspension of the respective actinomycete. After 1-2 days of incubation on a rotary shaker at 30°C, 1 mL of culture was taken, and total DNA was extracted with the innuPREP Bacteria DNA Kit (Innuscreen GmbH) according to the manufacture's instruction.

**16SrRNA-gene sequencing**
For 16SrRNA gene sequencing, a PCR using the proofreading Q5 polymerase (NEB) with primers 27Fbac and 1492Runi and total DNA from *Nocardia brasiliensis* Tü 69 as the template was performed. PCR products were blunt end cloned into pJET1.2 (Thermo Scientific™) according to the manufacture's instruction. 16S rRNA gene sequence containing plasmids were sent for sequencing. Retrieved sequences were used for a BLASTN[13] against the rRNA/ITS databases.

**Construction of plasmids and strains for microscopy**
Plasmids used and constructed for microscopy are listed in Table 3. For the construction of plasmid pRM43mchMurJ, *sco3894 (murJ)* was amplified by PCR using primers GAsco3894fw and Sco3894HindIIIrev and total DNA of *S. coelicolor* M1154 as a template. The amplified *murJ* was then fused to the 3' end of *mCherry* in pRM43mch by restriction with BsrGI/HindIII and subsequent ligation leading to plasmid pRM43mchMurJ, which carries a *mCherry-murJ* gene fusion under transcriptional control of the constitutive *ermE** promoter. Correct cloning was confirmed by Sanger sequencing. For inducible expression from plasmid pIJ6902, the *mCherry-murJ* gene fusion was cleaved out using NdeI/EcoRI digestion and ligated to NdeI/EcoRI digested pIJ6902 resulting in plasmid pIJ6902-mChMurJ. As a control, the *mCherry* gene from pRM43mCh was cloned using the restriction sites NdeI/EcoRI into plasmid pIJ6902, resulting in plasmid pIJ6902-mCh. All constructs were introduced into *S. coelicolor* M1154 via intergeneric conjugation using *E. coli* ET12567/pUZ8002 carrying the respective plasmid as a donor. Ex-conjugants were streaked for sporulation on MS agar plates containing apramycin and harvested after 7 days of sporulation.

**Microscopy and image analysis**
To test the effect of imacidin treatment on the morphology of *Streptomyces coelicolor*, approx. $10^6$ spores were inoculated in 100 µl of GYM medium in a flat-bottom 96-well plate (Sarstedt) and incubated for 18-22 hours in a custom-made humidity chamber at 30 °C on a rotary shaker, then imacidin was added. DMSO was used as a control. After 4 hours, 10 µl of the treated culture were taken and mounted on a 1% agarose pad and covered with a coverslip. When cells were stained with BODIPY™ FL Vancomycin (VanFL), cells were induced with thiostrepton at 0.1 µg $mL^{-1}$ for the expression of mCh-MurJ or mCh from the beginning of the cultivation. Cells were stained for 5 min using 0.1 µg $mL^{-1}$ BODIPY™ FL Vancomycin. Imaging was performed using an Olympus BX60 microscope equipped with an Olympus UPlanFI 100 x oil objective (NA 1.3) and an Olympus BX-FLA reflected light fluorescence attachment or with a Nikon Eclipse Ti microscope with a perfect focus system and a CFI Plan-Apo DM 100x/1.45 oil Ph3 objective (Nikon) using an ORCA-Flash 4.0 LT camera (Hamamatsu). For visualizing mCherry by fluorescence microscopy, the TxRed-4040C filterset (excitation 562/40 BrightLine HC, beamsplitter HC 593, emission: 624/40 BrightLine HC; Semrock) was used. For visualization of BODIPY™ FL Vancomycin, the filterset

GFP-3035D (excitation: 472/30 BrightLine HC, beamsplitter HC 495, emission: 520/35 BrightLine HC; Semrock) was used. Image acquisition was carried out using NIS Elements Advanced Research (Nikon).

Time lapse microscopy was started by using pre-grown mycelium or, alternatively, with spores. To prepare the mycelium, cover slips were inserted into a GYM agar-containing petri dish at a 45-degree angle. Five microliter spore suspension (~ $10^6$ CFU $mL^{-1}$) of the respective *S. coelicolor* strain were pipetted on the edge between the agar and the cover slip. After 16-22 hours of incubation at 30°C, the cover slip was removed from the agar plate, cleaned on one side, and mounted onto a thin agarose pad. When spores were used as a starting material, 1 µl of a spore suspension ( ~ $10^6$ CFU $mL^{-1}$) was pipetted directly on the agarose pad. The thin agarose pad was prepared beforehand and mounted in a custom-made metal carrier as follows: a cover slip consisting of an air permeable polymer (IBIDI) was glued over a hole in the metal carrier using a gene frame (Thermo Scientific™). Later, when the imaging setup is assembled completely the hole in the metal carrier ensures oxygen supply for the growing streptomycetes. A second gene frame was glued on top of the air-permeable polymer. The 1.5 % agarose (w/v) was boiled in either GYM medium or 25% GYM medium, poured into the second gene frame and was immediately covered with a clean glass slide. After the agarose solidified, the glass slide was carefully removed, and the cover slip with mycelia was glued on top of the agarose pad using the gene frame. Alternatively, 1 µl of the spore suspension was spotted onto the agar pad and was covered with a cover slip after drying. Time lapse imaging was performed using a Nikon Eclipse Ti microscope equipped as described above and a cage incubator (Okolab) at 30°C. Images were taken every 5-10 min. Image analysis was performed using ImageJ Fiji[14].

**Western blot of candidate MurJ homologs**

Cell lysates of MurJ homologs were run for 3 hours at 80 V on a NuPAGE Bis-Tris gel (Invitrogen). The iBlot™ 2 Gel Transfer Device (ThermoFisher) was run for 7 min at 20 V to transfer the proteins to a membrane. The Monoclonal ANTI-FLAG® M2 antibody produced in mouse (MilliporeSigma) (1:2000) and ANTI-HIS TAG antibody produced in rabbit (MilliporeSigma) were used as primary antibodies (1:1000). Irdye® 680RD Donkey Anti-Mouse IGG Secondary Antibody 680 (LICORbio) and Irdye® 800CW Goat Anti-Rabbit IGG Secondary Antibody (LICORbio) were used as secondary antibodies to visualize the protein.

**Construction of *Streptomyces* Tü 22 $MurJ^{WT}$ and $MurJ^{G310D}$ expression vectors**

We modified pNYCOMPs to express either the wildtype MurJ protein $MurJ^{WT}$-$(GSSS)_3$-PPX-$(GSSS)_3$-sfGFP-FLAG-$His_{10}$ (pNYCOMPs-$MurJ^{WT}$) or the G310D mutant $MurJ^{G310D}$-$(GSSS)_3$-PPX-$(GSSS)_3$-sfGFP-FLAG-$His_{10}$ (pNYCOMPs-$MurJ^{G310D}$). All strains, primers, and plasmids used for purification can be found in Tables 1-3. USER® Cloning was to construct both pNYCOMPs-$MurJ^{WT}$ and pNYCOMPs-$MurJ^{G310D}$. To construct pNYCOMPs-$MurJ^{WT}$, the pNYCOMPs plasmid was amplified by PCR using primers CJ_746 and CJ_750. MurJ was amplified by PCR from a codon-optimized Tü 22 TWIST gene and cloned into pNYCOMPs using primers CJ_751 and CJ_871. A TEV protease cleavage site with extended GSSS linkers was cloned into pNYCOMPs using primers CJ_821 and CJ_872. We cloned sfGFP into pNYCOMPs using primers CJ_749 and CJ_818 to track protein expression levels. To construct pNYCOMPs-$MurJ^{G310D}$, primers CJ_946 and CJ_947 were phosphorylated and used in a blunt ligation to introduce the G310D mutation.

**Purification of *Streptomyces* Tü 22 $MurJ^{WT}$ and $MurJ^{G310D}$**

Growth and purification protocols were adapted from previously established methods[15–19]. BL21-AI *E. coli* was transformed with either pNYCOMPs-$MurJ^{WT}$ or pNYCOMPs-$MurJ^{G310D}$ and plated onto LB agar plates containing kanamycin (25 µg $mL^{-1}$). A single transformed colony was used to start a 50 mL overnight culture. 8 mL of starter culture was used to inoculate 4 L of terrific broth (IBI Scientific) containing kanamycin (25 µg $mL^{-1}$). Cultures were grown at 37°C and once they reached an $OD_{600}$ of ~0.7 (2.5-3 hours), protein expression was induced with 0.5 mM isopropyl-β-d-thiogalactopyranoside (IPTG) and 0.2% L-arabinose. Induced cultures were grown for 18 hours at 15°C then harvested by centrifugation at 4000*g* for 15 min at 4°C.

Cell pellets were resuspended in 25 mL of lysis buffer (500 mM NaCl, 20 mM HEPES pH 7.5, 10% glycerol). Resuspended pellets had 1 mM phenylmethyl-sulfonyl fluoride (PMSF), 2 mM β-mercaptoethanol (BME), and one complete EDTA-free protease inhibitor cocktail (Roche) added. Cell lysates were sonicated and then solubilized in 30 mM n-dodecyl-β-D-maltoside (DDM) rotating for 1.5 hours at 4°C. The lysates were clarified by centrifugation at 58,500*g* for 40 min at 4°C. During centrifugation 2 mL of TALON Cobalt resin was equilibrated with 20 bed volumes (BV) equilibration buffer (500 mM NaCl, 20 mM HEPES pH 7.5, 10% glycerol, 2 mM BME, 1 mM DDM) for subsequent immobilized metal affinity chromatography (IMAC). The lysate was combined with resin in a 50 mL falcon tube and rotated for 45 min at 4°C to allow the his-tagged protein to bind. To prevent off-target binding, 5 mM of imidazole was added to the lysate-resin mixture. The lysate supernatant was removed after centrifugation at 700*g* for 5 min. Subsequent washing of the resin was performed by batch washing. Briefly, the resin was mixed with 10-20 BV of wash buffer (500 mM NaCl, 20 mM HEPES pH 7.5, 10% glycerol, 2 mM BME, 1 mM DDM) and then centrifuged at 1000*g* for 2 min. Supernatant from the wash was removed from the tube. The wash step was repeated 3-4 times until no more protein was seen in the supernatant (identified by using the Bio-Rad Protein Assay Dye Reagent Concentrate). After washing, the resin was resuspended in 1 BV of wash buffer and transferred to a gravity column. The resin was incubated in 2 BV of elution buffer (500 mM NaCl, 20 mM HEPES pH 7.5, 10% glycerol, 1 mM DDM, 2 mM BME, 250 mM imidazole) for 10 min and eluted off. This elution step was repeated 2 more times. The elution fractions containing MurJ (confirmed by a green color from sfGFP & Bio-Rad Protein Assay Dye Reagent Concentrate) were concentrated using a 30-kilodalton molecular weight centrifugal concentrator (MilliporeSigma).

MurJ was further purified by size exclusion chromatography (SEC) on an SRT-10 SEC 300 column in a reduced salt buffer (150 mM NaCl, 20 mM HEPES pH 7.5, 1 mM DDM, 2 mM BME). Fractions containing MurJ (confirmed by SDS-PAGE) were combined and incubated overnight at 4°C with 2 x 10 mg $mL^{-1}$ his-tagged TEV protease. After cleavage, fractions were incubated with 1.5 mL of TALON Cobalt resin for 10 min to remove cleaved sfGFP protein and his-tagged TEV protease from solution. MurJ samples were concentrated to ~20 µM using a 30-kilodalton molecular weight centrifugal concentrator, then flash frozen using liquid nitrogen.

**Differential scanning fluorimetry (DSF)**
All DSF experiments were carried out in a 384 well PCR plate (Corning), sealed with an Applied biosystems MicroAMP Optical Adhesive Film (ThermoFisher), and kept on ice. The BODIPY FL L-cystine (BFC) probe[20,21] (Invitrogen) was diluted to a final concentration of 4 µM. $MurJ^{WT}$ and $MurJ^{G310D}$ were buffer-exchanged into BME-free buffer (150 mM NaCl, 20 mM HEPES pH 7.5, 1 mM DDM) using Pierce™ Protein Concentrators PES, 30K molecular weight cutoff (MWCO) to prevent the probe from reacting with the free thiol groups on BME. The proteins were concentrated to 4 µM and imacidin was diluted to 4.4 µM for final DSF measurements.

DSF measurements were performed with a QuantStudio™ 7 Flex Real-Time PCR System. Melting curve experiments were set to cool down to 4°C as quickly as possible to start. Plates were incubated at 4°C for 15 min, then began ramp up at 1°C per minute until it reached 95°C. Multicomponent data was exported to Microsoft Excel, and thermal shift data was analyzed in Protein Thermal Shift™ software by Thermo Fisher Scientific. The melting temperatures were determined by the Boltzmann derivative. The means of the experimental melting temperatures were compared using an unpaired t test with Welch's correction.

**Synthesis of imacidin and taeanamide**

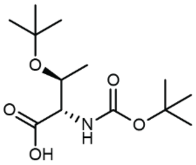

**Boc-L-*allo*-Thr(tBu)-OH (1):** 1 g of L-allo-Thr(tBu)-OH (5.7 mmol; Chem-Impex) was added to a round bottom flask with DIPEA (2.18 g; 2.94 mL; 16.8 mmol) in 30 mL DCM. Boc anhydride (2.74 ; 2.88 mL; 12.6 mmol) was added and the mixture was sealed with a septum and stirred at room temperature under $N_2$ for 18 hours. The resulting solution was dried, then partitioned between ethyl acetate and water. The aqueous fraction was dried by rotary evaporation, yielding a sticky clear oil.

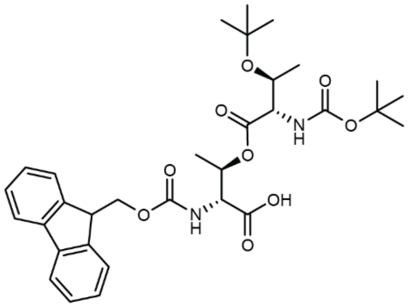

**Fmoc-D-*allo*-Thr(Boc-L-*allo*-Thr(tBu))-OH (2):** 110 mg (1 eq) of Fmoc-D-allo-threonine was loaded onto 300 mg of 2-CTC resin, swollen in DCM, by addition of 9 eq. (250 µL) of DIPEA. Loading proceeded over 2 hours, shaking at room temperature. After washing the loaded resin with DCM, the Boc-protected threonine (**1**) was coupled by a Steglich esterification. Fmoc- D-allo-threonine on resin (1 eq.; 0.135 mmol) was mixed with 3.5 eq. DIC (64 µL) and 5 eq. **1** (0.675 mmol) resuspended in DCM. The vessel was filled with anhydrous DCM and the reaction was initiated by addition of 0.4 eq. DMAP (4 mg), before being sealed and shaken overnight at room temperature. Once completed, the resin was washed with DCM, drained, and eluted using 2x 5 mL 20% HFIP in DCM. The eluate was dried and then partitioned between ethyl acetate and water. The organic fraction was dried under rotary evaporation, yielding pure **2**.

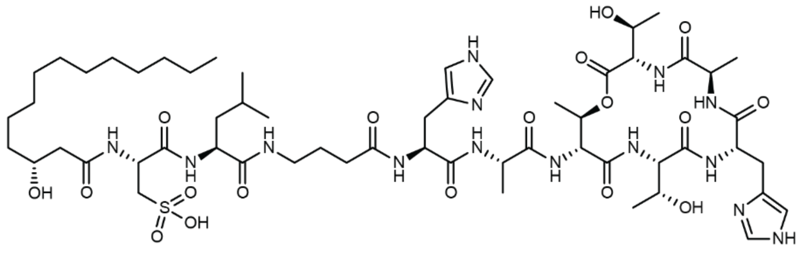

**Imacidin (3):** 300 mg of swollen 2-CTC resin (calculated loading capacity 0.45 mmol/g) was loaded with 3 eq. Fmoc-D-alanine (140 mg) and 9 eq. DIPEA (250 µL) for 2 hours, shaking at room temperature. Loaded resin was washed with DCM and DMF, deprotected with 20%

piperidine in DMF (2x 5 mL), and washed against with DMF, DCM, and DMF. Fmoc protected amino acids (3 eq.) were loaded with 3 eq. HATU (255 mg) and DIPEA (95 µL). After capping the final amine by addition of the acyl tail, the linear peptide was released from resin by treatment with 2 mL of a TFA mixture (95% TFA, 2.5% water, 2.5% TIPS) for 30 minutes (2x). The eluate was dried under nitrogen. Linear imacidin was the major product, and it was purified by reverse phase HPLC. Cyclization was achieved by resuspending the linear imacidin in DMF with 8 eq. PyAOP and 30 eq. DIPEA, mixing overnight at room temperature. Cyclized imacidin was purified by HPLC.

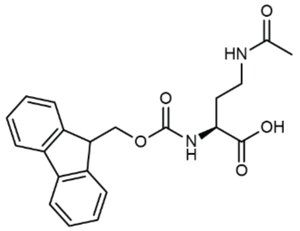


**Fmoc-4-*N*-acetyl-L-2,4-diaminobutyric acid (4):** 1 g of 4-*N*-acetyl-L-2,4-diaminobutyric acid (6.24 mmol; Chem-Impex) was added to a round bottom flask with 12 mL 10% $Na_2CO_3$ and 5 mL dioxane. Fmoc N-hydroxysuccinimide ester (2.21 g; 6.55 mmol) in 9 mL dioxane was added dropwise to the amino acid before this mixture was sealed with a septum and stirred at room temperature under $N_2$ for 18 hours. The resulting solution was dried, then partitioned between diethyl ether and water. The aqueous fraction was acidified to pH 3.5, extracted with ethyl acetate, and dried over $MgSO_4$. Fmoc-4-*N*-acetyl-L-2,4-diaminobutyric acid was then purified by silica gel column chromatography. The cleaned reaction mixture was dried onto silica, placed on a silica gel column and washed with 70% hexanes and 30% ethyl acetate, then ethyl acetate with 0.1% formic acid, before eluting in 90% ethyl acetate and 10% methanol with 0.1% formic acid. The eluent was dried to yield 700 mg of pure Fmoc-4-*N*-acetyl-L-2,4-diaminobutyric acid (**4**).

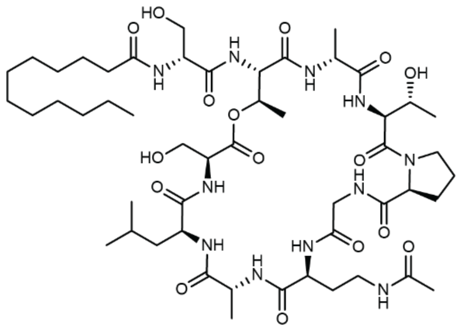

**Taeanamide (5):** 600 mg of swollen 2-CTC resin (calculated loading capacity 0.45 mmol/g) was loaded with 3 eq. Fmoc-L-leucine (300 mg) and 9 eq. DIPEA (510 µL) for 2 hours, shaking at room temperature. Loaded resin was washed with DCM and DMF, deprotected with 20% piperidine in DMF (2x 5 mL), and washed against with DMF, DCM, and DMF. Fmoc protected amino acids (3 eq.) or lauric acid were loaded with 3 eq. HATU and 3 eq. DIPEA. To incorporate the ester-linked serine residue, we had initially attempted to follow our method used for imacidin. Steglich esterification conditions were used to attach L-Boc-Ser(tbu)-OH to the alcohol of a resin bound threonine, which could then be eluted by HFIP and purified by HPLC. While this monomer could be added to the growing peptide chain during SPPS, we were unable to find conditions to attach the final components. Attachment of L-Boc-Ser(tbu)-OH to a growing peptide chain capped with Fmoc-L-Thr-OH could be achieved by Steglich conditions exclusively in DCM, but again, this peptide could not be extended. We ultimately settled on attaching L-Boc-Ser(tbu)-OH to a completed resin-bound peptide capped with lauric acid. 15 eq. L-Boc-Ser(tbu)-OH and 15 eq. DIC in 10 mL anhydrous DCM was mixed briefly and added to our resin-bound peptide, swollen in DCM. To initiate the reaction, 0.5 eq. DMAP in DCM was added. This reaction proceeded shaking at room temperature for 12 hours before being drained and repeated, achieving nearly complete conversion to the esterified product. Following a final wash with DMF and DCM, the resin was eluted with 5 mL of a TFA mixture (95% TFA, 2.5% water, 2.5% TIPS) for 30 minutes (2x). The eluate was dried under nitrogen and resuspended in methanol with a drop of ammonia hydroxide to cleave TFA-esters. Linear taeanamide was the major product, and it was purified by reverse phase HPLC. Cyclization was achieved by resuspending the linear taeanamide in DMF with 8 eq. PyAOP and 30 eq. DIPEA, mixing overnight at room temperature. Cyclized taeanamide was purified by HPLC.

**Table S1. Bacterial strains used in this study**

| Strain | Genotype | Reference |
|---|---|---|
| ***E. coli* strains** | | |
| *E. coli* XL1blue | *rec*A1 *end*A1 *gyrA*96 *thi*-1 *hsdR*17 *sup*E44 *rel*A1 *lac* F´ *pro*AB I*ac*IqZΔM15 Tn10 Tet$^r$ | Stratagene |
| *E. coli* ET12567/pUZ8002 | *dam, dcm, hsdM, hsdS, hsdR, cat, tet, kan* | Kieser *et al.*, 2000 |
| *E. coli* BL21-AI | *F⁻ompT hsdS$_B$ (r$_B$⁻, m$_B$⁻) gal dcm araB::T7RNAP-tetA* | Bhawsinghka *et al.*, 2020 |
| **Actinomycete strains** | **Genotype** | **Reference** |
| *Actinomadura madurae* DSM 43067 | | Leibniz Institute DSMZ-German Collection of Microorganisms and Cell Cultures |
| *Microbispora* ATCC 55410 | | American Type Culture Collection |
| *Nocardia asteroids* DSM 43132 | | Leibniz Institute DSMZ-German Collection of Microorganisms and Cell Cultures |
| *Nocardia brasiliensis* Tü 69 | | Tübingen strain collection |
| *Nocardia farcinica* DSM 43665 | | Leibniz Institute DSMZ-German Collection of Microorganisms and Cell Cultures |
| *Nocardia nova* DSM 44481 | | Leibniz Institute DSMZ-German Collection of Microorganisms and Cell Cultures |
| *Streptomyces bottropenensis* DSM 40262 | | Leibniz Institute DSMZ-German Collection of Microorganisms and Cell Cultures |
| *Streptomyces coelicolor* M1154 | Δ*act* Δ*red* Δ*cpk* Δ*cda rpoB* [S433L] | Gomez-Escribano and Bibb (2011)[22] |
| *Streptomyces coelicolor* M1154 pIJ6902-mCh | | This study |
| *Streptomyces coelicolor* M1154 pIJ6902-mChMurJ | | This study |
| *Streptomyces coelicolor* M1154 pIJ10257 | | This study |
| *Streptomyces coelicolor* M1154 pIJ10257-MurJ$^{orf6}$ | | This study |
| *Streptomyces coelicolor* M1154 pIJ10257-MurJ$^{imcL}$ | | This study |

| | | |
|---|---|---|
| *Streptomyces coelicolor* M1154 pIJ10257-MurJ$^{G310D}$ | | This study |
| *Streptomyces coelicolor* M1154 pIJ10257-MurJ$^{WT}$ | | This study |
| *Streptomyces diastatochomogenes* (Tü 6028) | | Tübingen strain collection |
| *Streptomyces edwardsiae* DSM 41636 | | Leibniz Institute DSMZ-German Collection of Microorganisms and Cell Cultures |
| *Streptomyces europaeiscabiei* AK02-03A | | Dr. Christopher Clarke |
| *Streptomyces europaeiscabiei* AK05-03A | | Dr. Christopher Clarke |
| *Streptomyces europaeiscabiei* AK08-03 | | Dr. Christopher Clarke |
| *Streptomyces europaeiscabiei* ND06-05F | | Dr. Christopher Clarke |
| *Streptomyces europaeiscabiei* WA08-01A | | Dr. Christopher Clarke |
| *Streptomyces olivaceus* (Tü 1379) | | Tübingen strain collection |
| *Streptomyces rochei* ATCC 10739 | | American Type Culture Collection |
| *Streptomyces ruber* DSM 40304 | | Leibniz Institute DSMZ-German Collection of Microorganisms and Cell Cultures |
| *Streptomyces violaceoruber* (Tü 22) | | Tübingen strain collection |
| *Streptomyces viridochromogenes* (Tü 57) | | Tübingen strain collection |
| *Streptomyces sp.* Tü 6071 | | Tübingen strain collection |
| **Other gram-positive strains** | **Genotype** | **Reference** |
| *Corynebacterium striatum* KPL 1959 | | Dr. Katherine Lemon |
| *Curtobacterium* KPL 2560 | | Dr. Katherine Lemon |
| *Dermacoccus* KPL 2534 | | Dr. Katherine Lemon |
| *Dermacoccus* KPL 2528 | | Dr. Katherine Lemon |
| *Staphylococcus aureus* ATCC 19095 | | American Type Culture Collection |

**Table S2. Primers used in this study**

| Primer name | Sequence 5’-3’ |
|---|---|
| GAsco3894fw | TGGACGAGCTGTACAAGGGCGGCGGGTCCGGCGGGTCCAACGC GCCGTACGACGGT |
| Sco3894HindIIIrev | TCCAAGCTTTCAGCGTCCCAGGCGTCCG |
| 27Fbac | AGAGTTTGATCMTGGCTCAG |
| 1492Runi | TACGGTTACCTTGTTACGACTT |
| CJ_366 | AGTTAATTAATCAGGGUCACCGACCAAGACGTGAAC |
| CJ_367 | ACCCTGATTAATTAACUAGAGTCGACCTGC |
| CJ_386 | AGTTACGGATGATUGCTGCACGCG |
| CJ_387 | AATTGGCCATGATUGCCTCTTTTGTTAATCTCTGGG |
| CJ_388 | AATCATGGCCAATUACGGCGACGAC |
| CJ_389 | AATCATCCGTAACUCCGGTCCGACATCGAC |
| CJ_584 | ATGAACGCCCCAUACGATGGTG |
| CJ_585 | ACCGCGAACCAUACCCACGAG |
| CJ_611 | ATGGTTCGCGGUCGCTTGGGCCGTCCCTGATTAATTAACTAGAGT CGACC |
| CJ_612 | ATGGGGCGTTCAUGATTGCCTCTTTTGTTAATCTCTGGG |
| CJ_613 | ACGACACTTTCCAAGTCGCTTACC |
| CJ_614 | CGAGGAGCCCCACGCCAA |
| CJ_746 | AGCGAGGATTAUAAAGATGATGATGATAAAC |
| CJ_749 | ATAATCCTCGCUGCCTTTGTAAAGCTCATCC |
| CJ_750 | ATATCTCCTTCTTAAAGTUAAACAAAATTATTTC |
| CJ_751 | AACTTTAAGAAGGAGATAUACCATG |
| CJ_818 | AGAGGGTGAAGGUGATGCTAC |
| CJ_821 | ACCTTCACCCTCUCCACGG |
| CJ_871 | ACCATACCCACCAUTGAGTTAAGTTCAGCAATGCGC |
| CJ_872 | ATGGTGGGTATGGUCCGTGG |
| CJ_946 | ATGATACGTTTACTGTTGCCTATACGC |
| CJ_947 | CCAGAAGTGCCGCTCCTAATG |
| CJ_1033 | ATGGACAGGCCGUACGAACAAAATAGC |
| CJ_1034 | AATTAATCAGGGUCAGCGCCCCAGACGGCCAC |
| CJ_1035 | ACCCTGATTAATUAACTAGAGTCGACC |
| CJ_1036 | ACGGCCTGTCCAUGATTGCCTCTTTTGTTAATCTCTGGG |

**Table S3. Plasmids used in this study**

| Name | Characteristics | Reference |
|---|---|---|
| pRM43mCh | *Int_ΦC31, attP, aac(3)IV, oriT,* $P_{ermE}$, *mCherry* | Günther Muth, unpublished |
| pRM43mChMurJ | pRM43mCherry with *murJ* (Sco3894) fused to the 3'end of *mCherry* | This study |
| pIJ6902 | *Int_ΦC31, attP, aac(3)IV, tsr, oriT,* $P_{tipA}$ | Huang *et al.*, 2005[23] |
| pIJ6902-mCh | *Int_ΦC31, attP, aac(3)IV, tsr, oriT,* $P_{tipA,}$ *mCherry* | This study |
| pIJ6902-mChMurJ | *Int_ΦC31, attP, aac(3)IV, tsr, oriT,* $P_{tipA,}$ *mCherry-murJ* | This study |
| pIJ10257 | *Int_ ΦBT1, ermE*P, oriT, HygR,* | Hong *et al.*, 2005[24] |
| pIJ10257-MurJ$^{orf6}$ | *Int_ ΦBT1, imcP, oriT, HygR, MurJ$^{orf6}$ (S. europaeiscabiei AK08-03A Taeanamide BGC)* | This study |
| pIJ10257-MurJ$^{imcL}$ | *Int_ ΦBT1, imcP, oriT, HygR, MurJ$^{imcL}$ (S. Tü 1379 Imacidin BGC)* | This study |
| pIJ10257-MurJ$^{G310D}$ | *Int_ ΦBT1, imcP, oriT, HygR, MurJ (S. coelicolor M1154 with G310D mutation)* | This study |
| pIJ10257-MurJ$^{WT}$ | *Int_ ΦBT1, imcP, oriT, HygR, MurJ$^{WT}$ (S. coelicolor M1154)* | This study |
| pNYCOMPs | *$T_7P$, TEV, FLAG, $HIS_{10}$, KanR* | Love *et al.*, 2010[25] |
| pNYCOMPs-MurJ$^{WT}$ | *$T_7P$, TEV, FLAG, $HIS_{10}$, KanR, sfGFP, codon-optimized Tü 22 MurJ$^{WT}$* | This study |
| pNYCOMPs-MurJ$^{G310D}$ | *$T_7P$, TEV, FLAG, $HIS_{10}$, KanR, sfGFP, codon-optimized Tü 22 MurJ$^{G310D}$* | This study |

**Table S4. MICs for imacidin congeners against *Streptomyces coelicolor* M1154**

| Imacidin Congener M+H | Acyl | Position 1 | MIC (μg mL$^{-1}$) |
|---|---|---|---|
| 1231 | C12 | L-aspartate | 0.064 |
| 1245 | C12 | L-glutamate | 0.032 |
| 1267 | C12 | L-cysteate | 0.016 |
| 1281 | C13 | L-cysteate | 0.008 |

**Table S5. Components of the imacidin biosynthetic gene cluster.** Closest characterized homologs were identified through BLAST searches in the UniProtKB-SwissProtdatabase (August 2025).

| Protein | Size (aa) | Putative function | Closest characterized homolog (%AA ID) | UniProtKB |
|---|---|---|---|---|
| ImcE | 746 | Bacterial transcription activator domain | PIN domain-containing protein (84%) | A0A0F0GNB9 |
| ImcF | 186 | Cysteine dioxygenase | Cysteine dioxygenase (66.3%) | A0A919B0R8 |
| ImcG | 361 | Cysteate synthase | Cysteate synthase (58.7%) | A0A1E7N8I9 |
| ImcD | 907 | NRPS (AT-A-T) | Amino acid adenylation domain-containing protein (59.2%) | A0A429RJH2 |
| ImcH | 283 | Keto reductase | NAD(P)-dependent oxidoreductase (83.8%)%) | A0A42RL02 |
| ImcI | 222 | Thioesterase | Thioesterase (69.9%) | A0A429RKY7 |
| ImcJ | 74 | MbtH | Protein MbtH (70.8%) | A0A1M5LZ43 |
| ImcB | 4805 | NRPS (C-A-T-C-A-T-C-A-T-C-A-T-E) | Amino acid adenylation domain-containing protein (53.7%) | A0A239MVS1 |
| ImcA | 2143 | NRPS (C-A-T-C-A-T) | Carrier domain-containing protein (49.3%) | A0A2G1XLS3 |
| ImcK | 307 | Thioesterase | Polyketide synthase thioesterase domain-containing protein (49.0%) | A0A0M8UWA8 |
| ImcC | 4548 | NRPS (C-A-T-C-A-T-C-A-T-E-C-A-T-TE) | Amino acid adenylation domain-containing protein (46.4%) | A0A944KV78 |
| ImcL | 539 | MurJ | Murein biosynthesis integral membrane protein MurJ (78.8%) | A0A964UNA2 |

**Table S6. Substrate predictions for imacidin NRPS proteins**

| Protein | Adenylation domain Stachelhaus code | Best Stachelhaus code match via antiSMASH 8.0.2 | Observed substrate |
|---|---|---|---|
| ImcA | **D G T K L D E V G K** | **D G T K L G E V G K** (L-Asparagine) | L-Cysteate |
| | **D A M L V G A V A K** | **D A M L V G A V C K** (L-Leucine) | L-Leucine |
| ImcB | **I D T T I T - - W K** | - | GABA |
| | **D A A T V A I V A K** | **D A A T L A A V A K** (L-Tyrosine) | L-Histidine |
| | **D V F S V A I V Y K** | **D V F S V A I V Y K** (L-Alanine) | L-Alanine |
| | **D F W N V G M V H K** | **D F W N V G M V H K** (L-Threonine) | L-allo-Threonine |
| ImcC | **D F W N V G M V H K** | **D F W N V G M V H K** (L-Threonine) | L-Threonine |
| | **D A A T V A I V A K** | **D A A T L A A V A K** (L-Tyrosine) | L-Histidine |
| | **D V F S V A I V Y K** | **D V F S V A I V Y K** (L-Alanine | L-Alanine |
| ImcD | **D F W N I G M V H K** | **D F W N I G M V H K** (L-Threonine) | L-allo-Threonine |

**Table S7. Summary of all MurJ mutations**

| Mutant Number | Nucleotide Change | Location in the Genome | Predicted Protein |
|---|---|---|---|
| 1 | C>T | 1,571,007 | Transcriptional regulator BldD |
| 1 | $(G)_{15>14}$ | 2,434,869 | Not in a predicted ORF |
| 1 | G>T | 4,178,795 | MurJ |
| 1 | C>T | 8,376,643 | RidA family protein |
| 2 | C>A | 1,446,608 | CDP-diacylglycerol--glycerol-3-phosphate 3-phosphatidyltransferase |
| 2 | $(G)_{15>13}$ | 2,434,868 | Not in a predicted ORF |
| 2 | C>A | 3,107,980 | Roadblock/LC7 domain-containing protein/ ATP-binding protein |
| 2 | C>G | 3,171,829 | Not in a predicted ORF |
| 2 | A>G | 4,179,282 | MurJ |
| 2 | $(CGGTCG)_{2>1}$ | 6,316,612 | APC family permease |
| 2 | A>G | 6,598,319 | Putative xylitol oxidase or alditol oxidase |
| 2 | G>A | 7,532,041 | Gamma-glutamylpolyamine synthetase GlnA3 |
| 5 | G>A | 1,059,314 | Right-handed parallel beta-helix repeat-containing protein |
| 5 | $(G)_{15>13}$ | 2,434,868 | Not in a predicted ORF |
| 5 | C>T | 3,040,704 | AAA family ATPase |
| 5 | C>T | 3,562,688 | Bifunctional uroporphyrinogen-III C-methyltransferase/uroporphyrinogen-III synthase |
| 5 | $(G)_{8>9}$ | 3,855,202 | Metallophosphoesterase |
| 5 | G>A | 4,179,293 | MurJ |
| 5 | +G | 4,550,665 | Hypothetical protein |
| 5 | C>A | 5,613,289 | HEXXH motif-containing putative peptide modification protein |
| 5 | G>A | 7,520,837 | Hypothetical protein |
| 5 | T>G | 8,407,758 | Alpha/beta fold hydrolase |
| 6 | G>A | 47,788 | YfbM family protein |
| 6 | 2 bp >AA | 590,355 | Not in a predicted ORF |
| 6 | G>A | 667,594 | TetR/AcrR family transcriptional regulator |
| 6 | G>A | 1,089,221 | Alpha-xylosidase |
| 6 | A>G | 1,446,824 | CDP-diacylglycerol-glycerol-3-phosphate 3-phosphatidyltransferase |
| 6 | C>A | 1,687,394 | ATP-binding protein |
| 6 | C>G | 1,687,553 | ATP-binding protein |

**Table S8. Summary of all MurJ mutations continued**

| Mutant Number | Nucleotide Change | Location in the Genome | Predicted Protein |
|---|---|---|---|
| 6 | G>A | 1,689,445 | Hypothetical protein |
| 6 | $(G)_{15>13}$ | 2,434,868 | Not in a predicted ORF |
| 6 | T>G | 2,750,487 | Not in a predicted ORF |
| 6 | 2 bp >GC | 3,008,344 | Lysine decarboxylase DesA |
| 6 | +C | 3,146,273 | DUF2510 domain-containing protein |
| 6 | $(G)_{6>7}$ | 3,805,252 | dTMP kinase |
| 6 | A>G | 4,083,446 | D-alanyl-D-alanine carboxypeptidase family protein |
| 6 | 2 bp >AA | 4,178,780 | MurJ |
| 6 | T>C | 4,586,740 | GNAT family N-acetyltransferase |
| 6 | +CG | 5,119,083 | DUF5691 domain-containing protein |
| 6 | T>G | 7,617,891 | ABC transporter substrate-binding protein |
| 7 | C>T | 143,962 | BON domain-containing protein |
| 7 | 2 bp >TT | 145,396 | Not in a predicted ORF |
| 7 | G>A | 4,178,780 | MurJ |
| 7 | Δ246 bp | 5,718,499 | F0F1 ATP synthase |
| 8 | G>A | 640,636 | Rhodanese-like domain-containing protein |
| 8 | C>G | 640,658 | Rhodanese-like domain-containing protein |
| 8 | C>T | 640,663 | Rhodanese-like domain-containing protein |
| 8 | C>T | 640,669 | Rhodanese-like domain-containing protein |
| 8 | C>G | 3,088,266 | Not in a predicted ORF |
| 8 | C>A | 3,107,980 | Roadblock/LC7 domain-containing protein/ ATP-binding protein |
| 8 | +C | 3,146,273 | DUF2510 domain-containing protein |
| 8 | Δ136,640 bp | 4,071,393 | Many proteins |
| 8 | C>A | 4,314,891 | MurJ |
| 8 | C>T | 4,315,963 | MurJ |
| 8 | +G | 4,687,305 | Hypothetical protein |
| 8 | +GAG | 5,097,670 | Not in a predicted ORF |
| 8 | +CG | 5,255,720 | DUF5691 domain-containing protein |
| 8 | G>T | 6,453,773 | APC family permease |

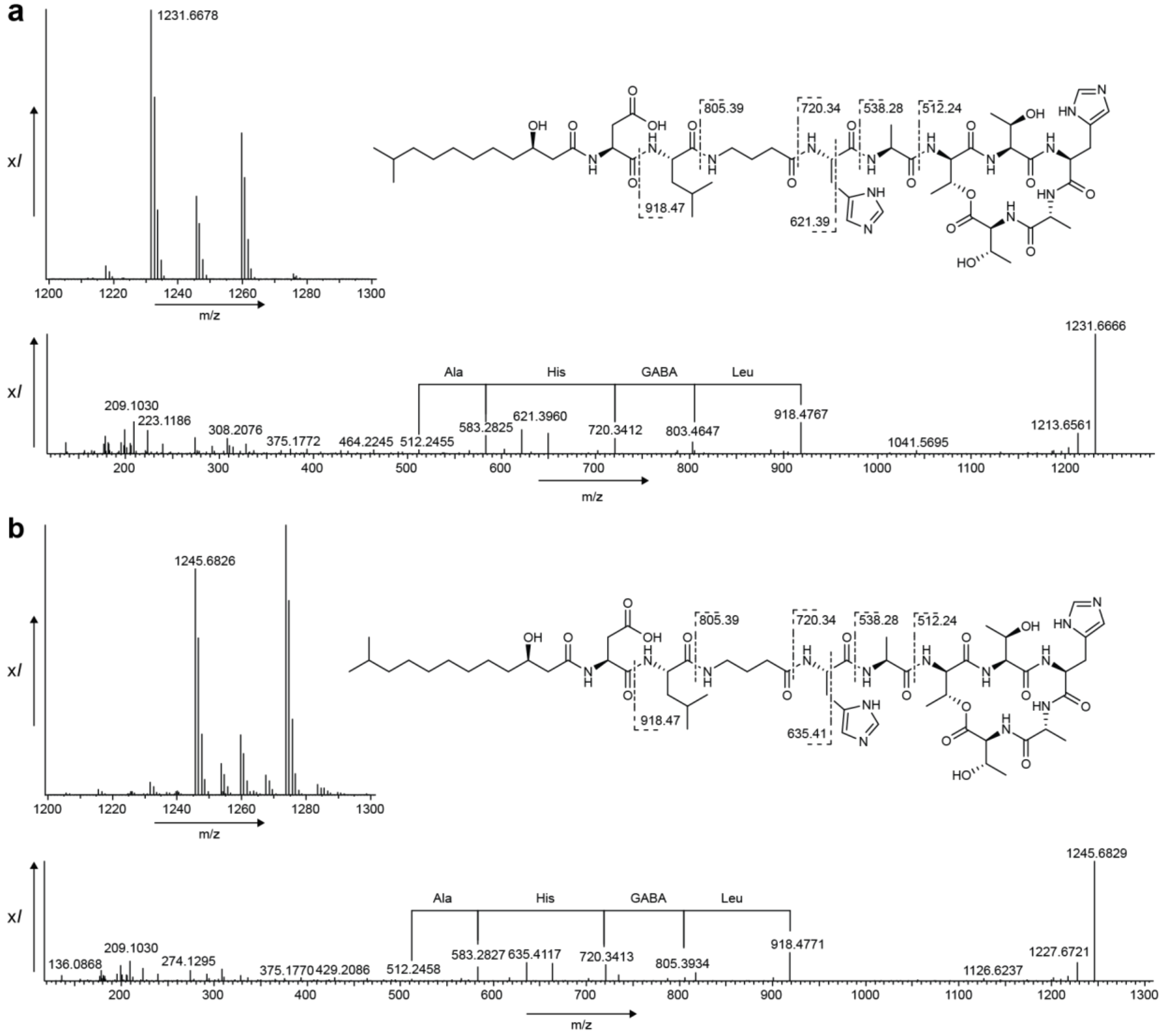


**Figure S1. High resolution mass spectrometry data for aspartate congeners of imacidin.** (**a**) Chemical structure and parent ion of *m/z* $[M+H]^+$ 1231.66 along with MS/MS spectra. (**b**) Chemical structure and parent ion of *m/z* $[M+H]^+$ 1245.68 along with MS/MS spectra.

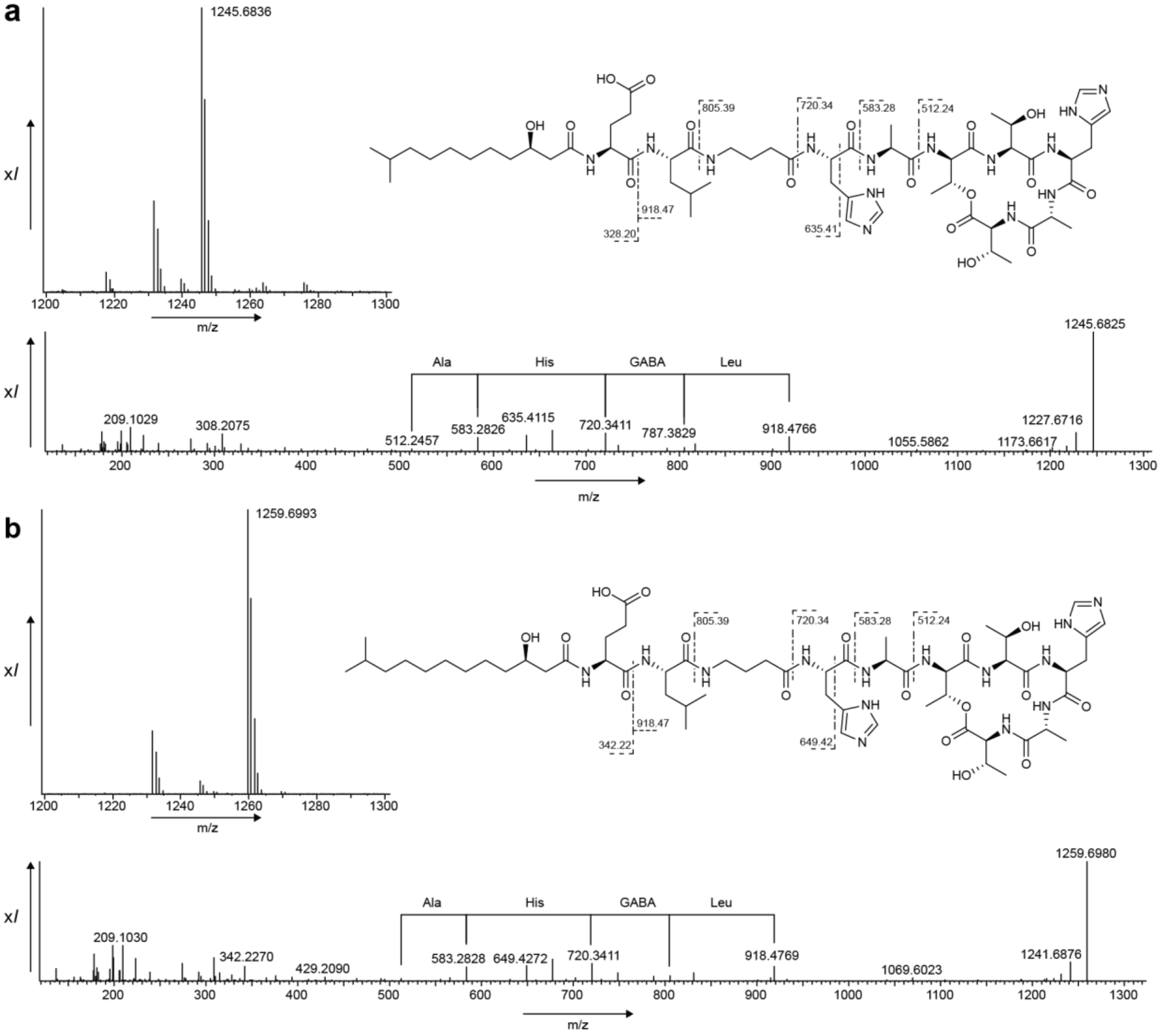


**Figure S2. High resolution mass spectrometry data for glutamate congeners of imacidin.** (**a**) Chemical structure and parent ion of *m/z* $[M+H]^+$ 1245.68 along with MS/MS spectra. (**b**) Chemical structure and parent ion of *m/z* $[M+H]^+$ 1259.69 along with MS/MS spectra.

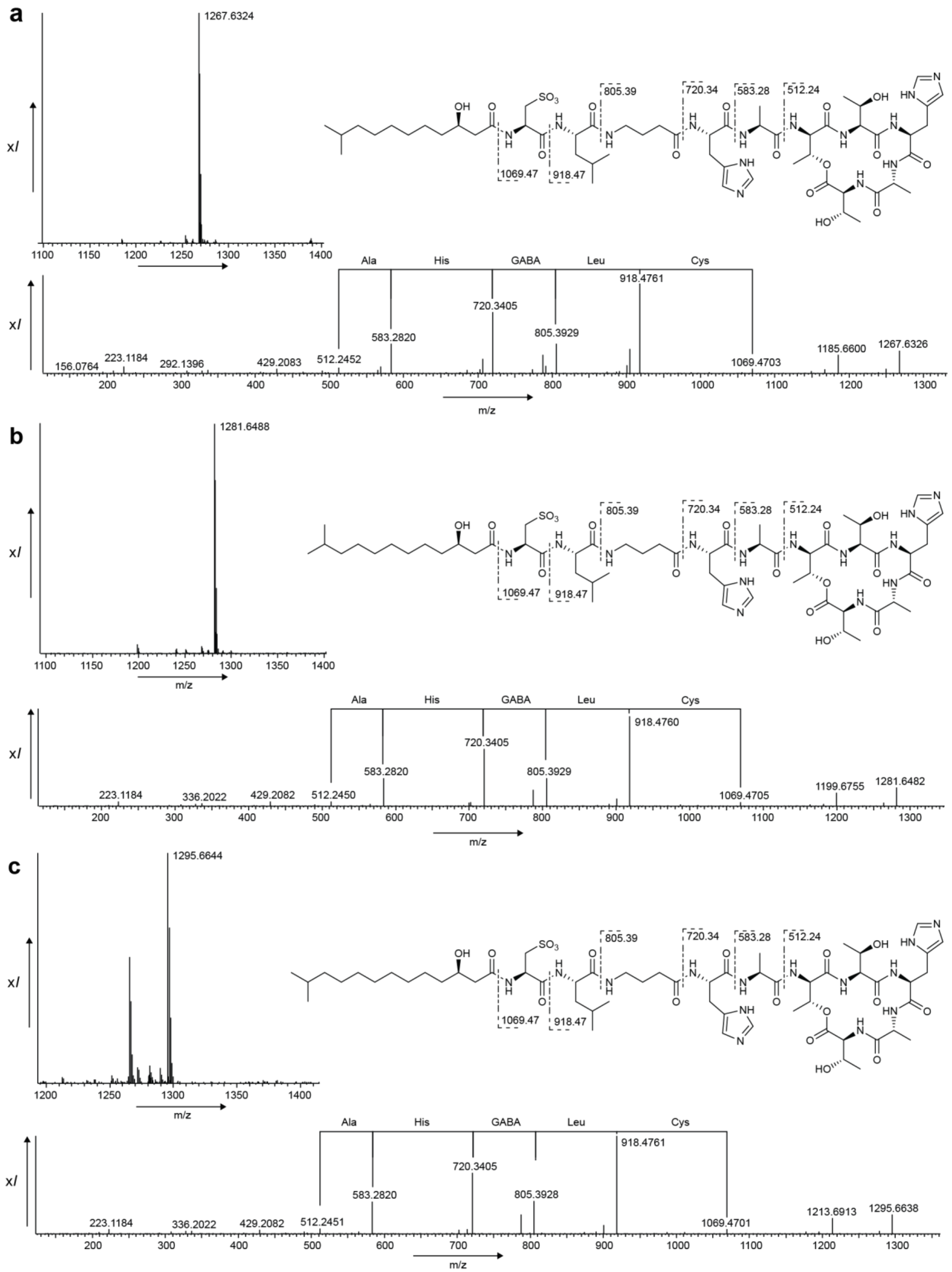


**Figure S3. High resolution mass spectrometry data for cysteate congeners of imacidin.** (**a**) Chemical structure and parent ion of *m/z* $[M+H]^+$ 1267.63 along with MS/MS spectra. (**b**) Chemical structure and parent ion of *m/z* $[M+H]^+$ 1281.64 along with MS/MS spectra. (**c**) Chemical structure and parent ion of *m/z* $[M+H]^+$ 1295.66 along with MS/MS spectra.

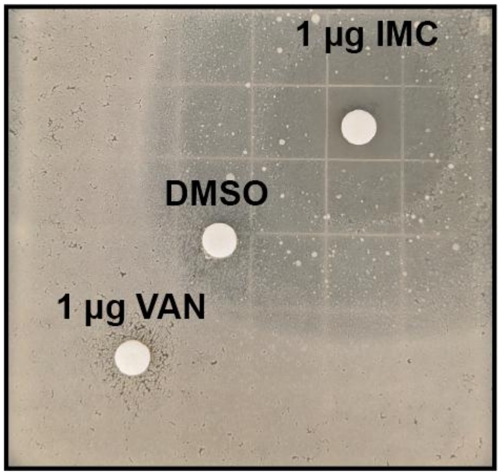


**Figure S4. *Streptomyces coelicolor* M1154 is imacidin-sensitive but vancomycin-resistant.** The activities of imacidin and vancomycin were tested against *S. coelicolor* M1154 using a disk diffusion assay with DMSO as a control. *S. coelicolor* M1154 growth was inhibited by imacidin but showed resistance to vancomycin indicating that while imacidin causes intracellular lipid accumulation like vancomycin, it is inhibiting a different target.

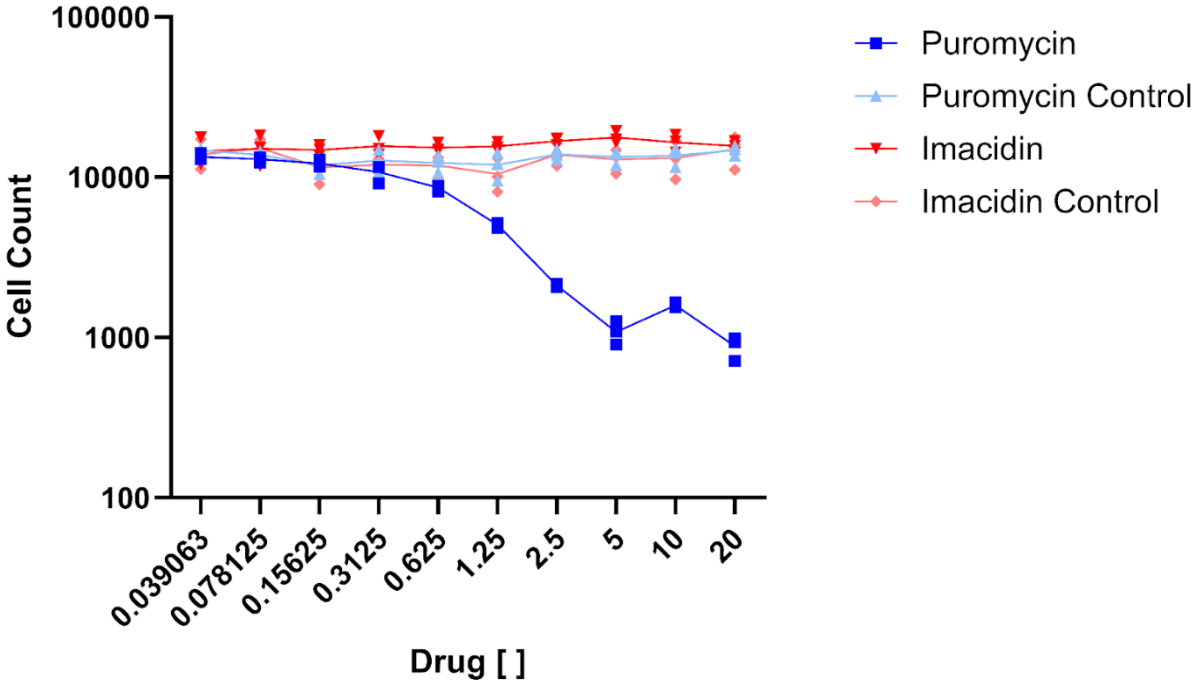


**Figure S5. Cytotoxicity assay of imacidin against hTERT-HME1 cells.** The cytotoxicity of imacidin was tested again the hTERT-HME1 human cell line. Puromycin was used as a negative control. The puromycin control and imacidin control were vehicle controls. Imacidin, the puromycin control, and the imacidin control had no effect on the growth of cells. Puromycin induced cell death at a concentration of 2.5 µg $mL^{-1}$.

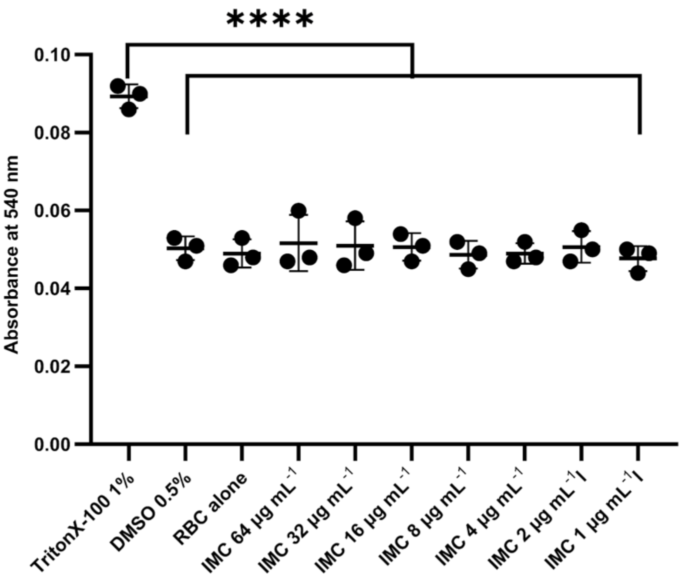


**Figure S6. Red blood cell (RBC) hemolysis assay.** Imacidin's hemolytic activity was tested against sheep red blood cells. Triton™X-100 was used as a positive control while DMSO and RBC alone were used as negative controls. There was a significant difference in absorbance at 540 nm between the positive control Triton™X-100 and the imacidin concentrations tested (1-64 μg $mL^{-1}$) indicating that imacidin had no hemolytic activity.

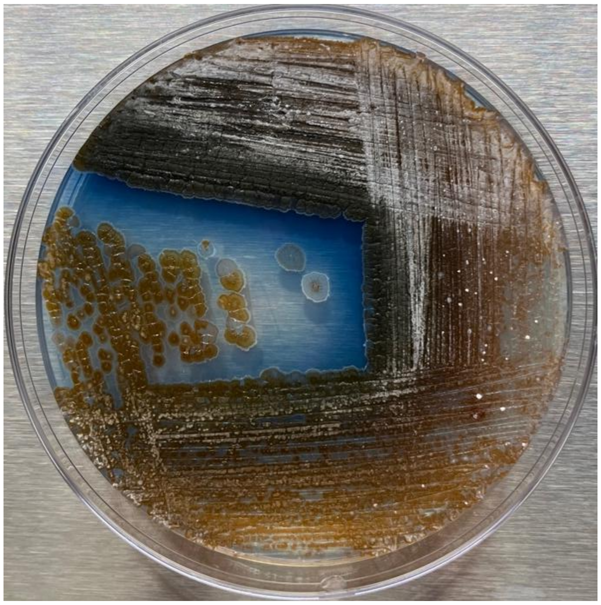

**Figure S7. Imacidin producer strain *Streptomyces* Tü 1379.** The imacidin producer strain *Streptomyces* Tü 1379 is closely related to *S. coelicolor* M1154; however, imacidin inhibits *S. coelicolor* M1154 growth but does not affect its producer strain. *S. coelicolor* M1154 makes actinorhodin, a compound that gives the species a distinctive blue color when grown on specific media (R2YE). When *Streptomyces* Tü 1379 is grown on R2YE media, it produces actinorhodin showing how closely related these species are.

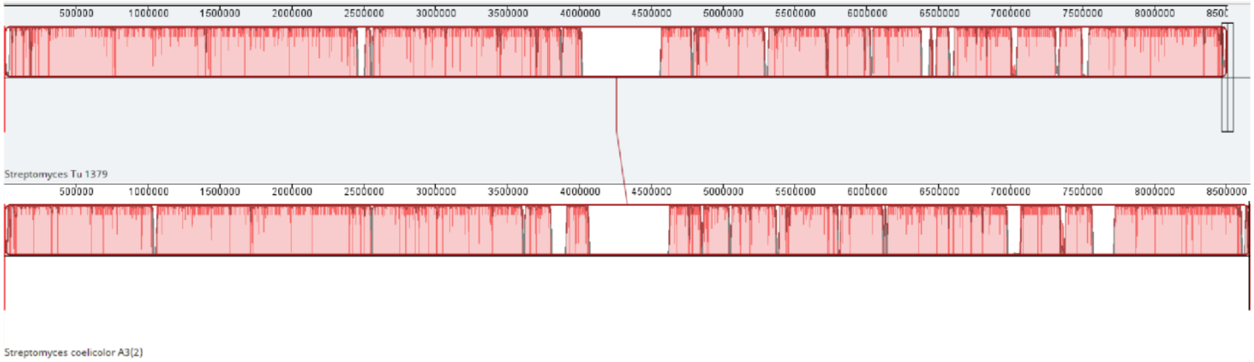


**Figure S8. Mauve alignment of *Streptomyces* sp. Tü 1379 and *Streptomyces coelicolor* A3(2).** The genomes of *Streptomyces* Tü 1379 and *Streptomyces coelicolor* A3(2) were overlayed using Mauve alignment. The overlay shows that the genomes have a majority overlapping and identical sites.

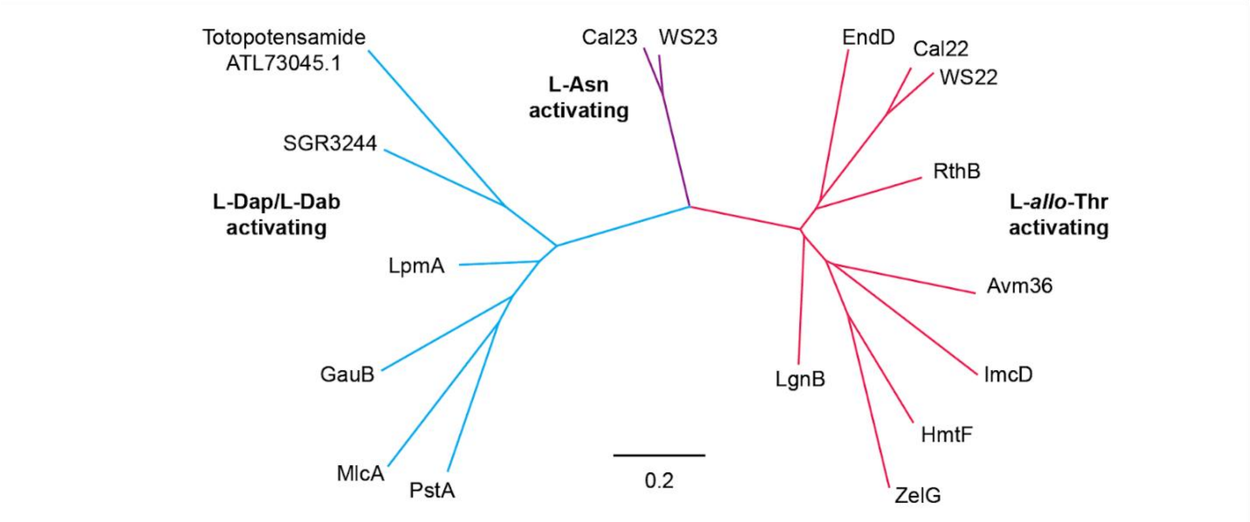


**Figure S9. Phylogenetic tree of NRPS stand-alone enzymes.** *ImcD* in the imacidin BGC encodes for a stand-alone NRPS enzyme that delivers L-*allo*-Thr in *trans*. For comparison, we searched for other stand-alone NRPS enzymes in the MiBIG database. When compared phylogenetically to other stand-alone NRPS enzymes, ImcD groups with the L-*allo*-Thr activating enzymes.

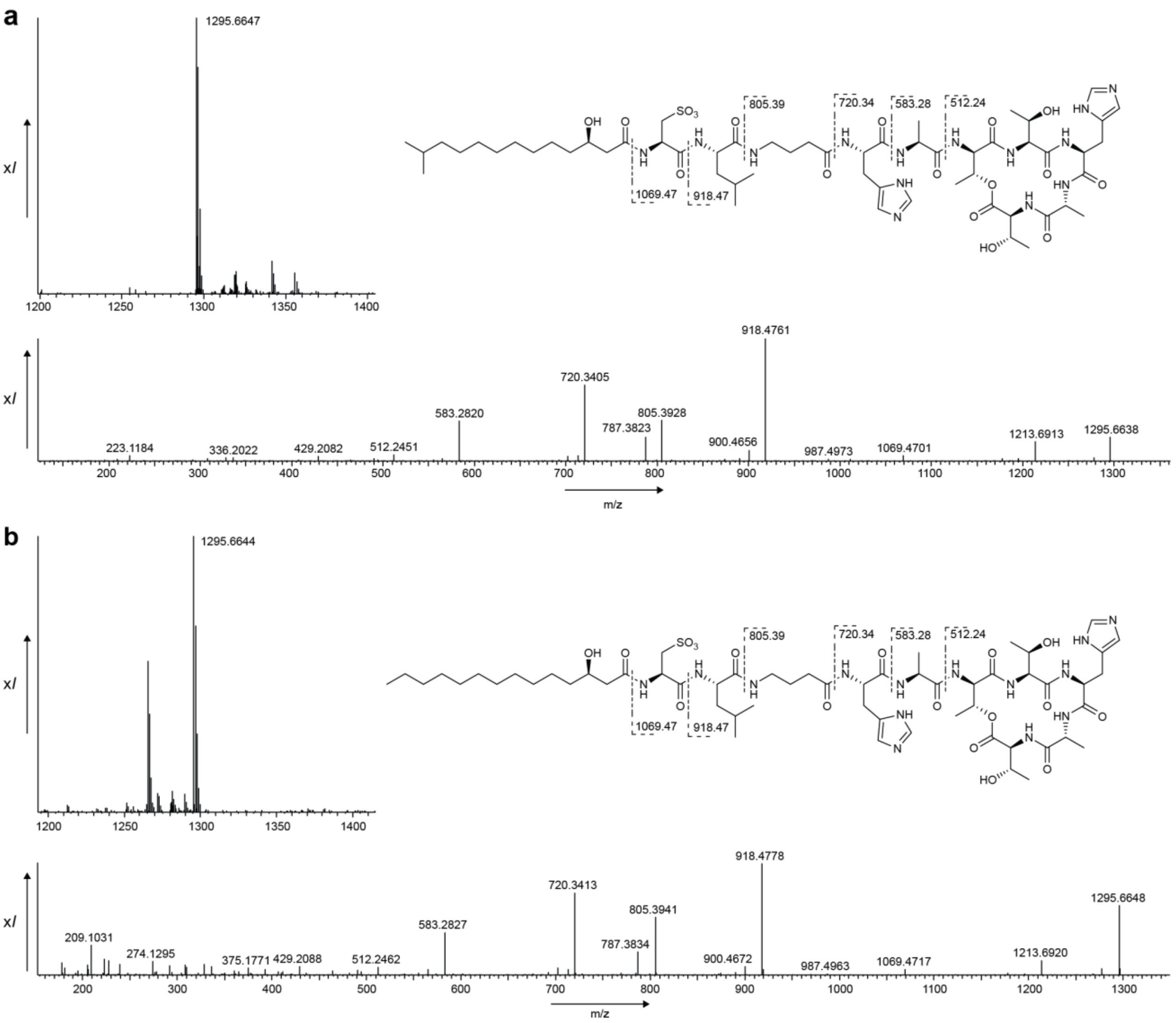


**Figure S10. High resolution mass spectrometry data comparing natural and synthetic imacidin.** (a) Synthetic Imacidin chemical structure and parent ion of *m/z* [M+H]+ 1295.66 along with MS/MS spectra. (b) Natural Imacidin chemical structure and parent ion of *m/z*[M+H]+ 1295.66 along with MS/MS spectra.

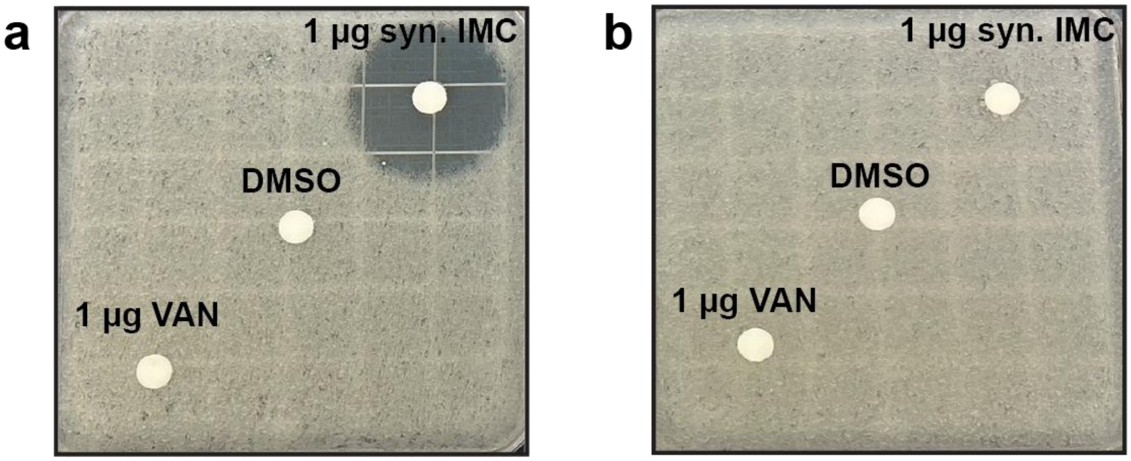


**Figure S11. Synthetic imacidin disk diffusion assay.** Using our solid-phase peptide synthesis scheme, we produced a synthetic version of imacidin and used disk diffusion assays to determine if synthetic imacidin retained activity. (a) Synthetic imacidin is tested against wildtype *S. coelicolor* M1154 and shown to inhibit growth. (b) Synthetic imacidin is tested against *S. coelicolor* M1154 expressing $MurJ^{G310D}$ and shown to have no activity.

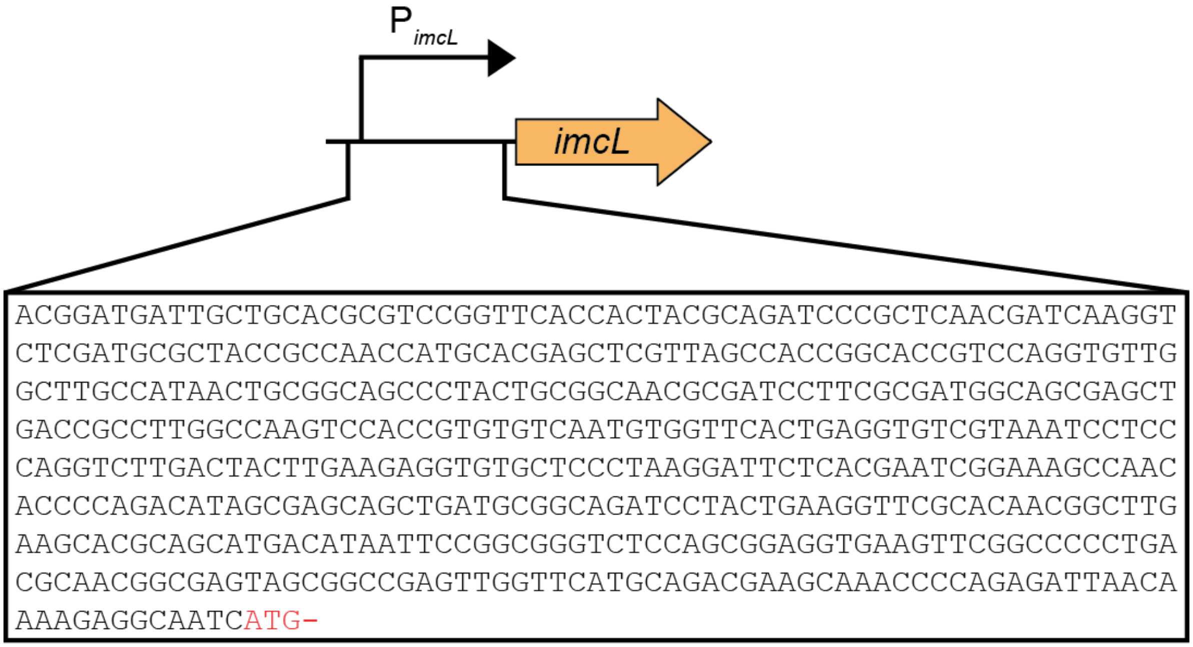


**Figure S12. Design of the modified pIJ10257 vector with the imacidin BGC *imcL* promoter.** For genetic complementation experiments a modified pIJ10257 expression vector was used. The original promoter was exchanged for the *imcL* promoter found in the imacidin BGC with the sequence shown above.

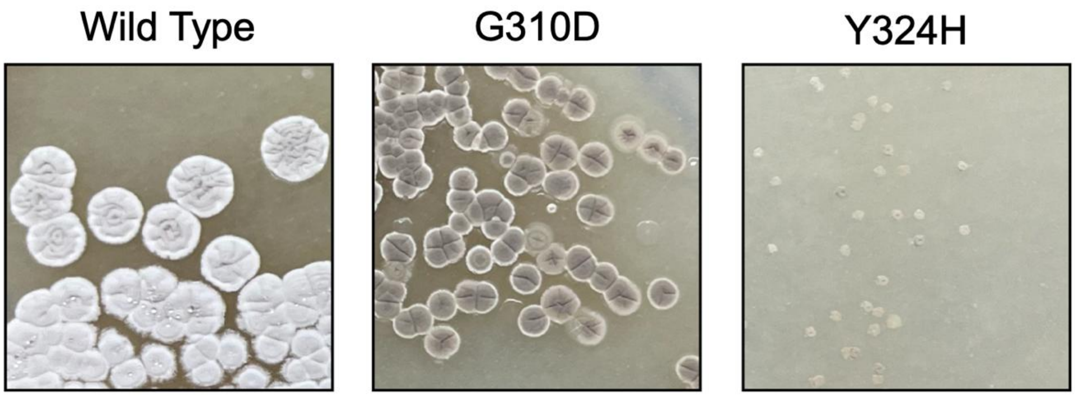


**Figure S13. Growth phenotypes of genetically complemented *Streptomyces coelicolor* M1154 mutants.** Expression of $MurJ^{WT}$ does not cause any outward changes to the morphology of *S. coelicolor* M1154. $MurJ^{G310D}$ expression leads to a decrease in colony size and reduction in spores. The expression of $MurJ^{Y324H}$ leads to a substantial reduction in overall growth.

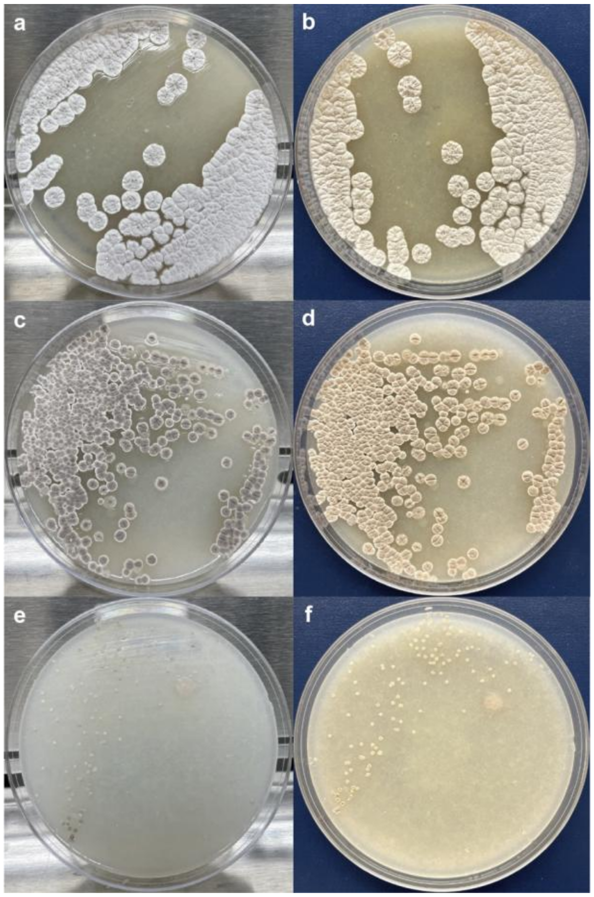


**Figure S14. Growth phenotypes of MurJ mutants on two backgrounds.** (a) *S. coelicolor* M1154 empty pIJ10257 vector against silver background. (b) *S. coelicolor* M1154 empty pIJ10257 vector against blue background. (c) *S. coelicolor* M1154 $MurJ^{G310D}$ mutant against silver background. (d) *S. coelicolor* M1154 $MurJ^{G310D}$ mutant against blue background. (e) *S. coelicolor* M1154 $MurJ^{Y324H}$ mutant against silver background. (f) *S. coelicolor* M1154 $MurJ^{Y324H}$ mutant against blue background.

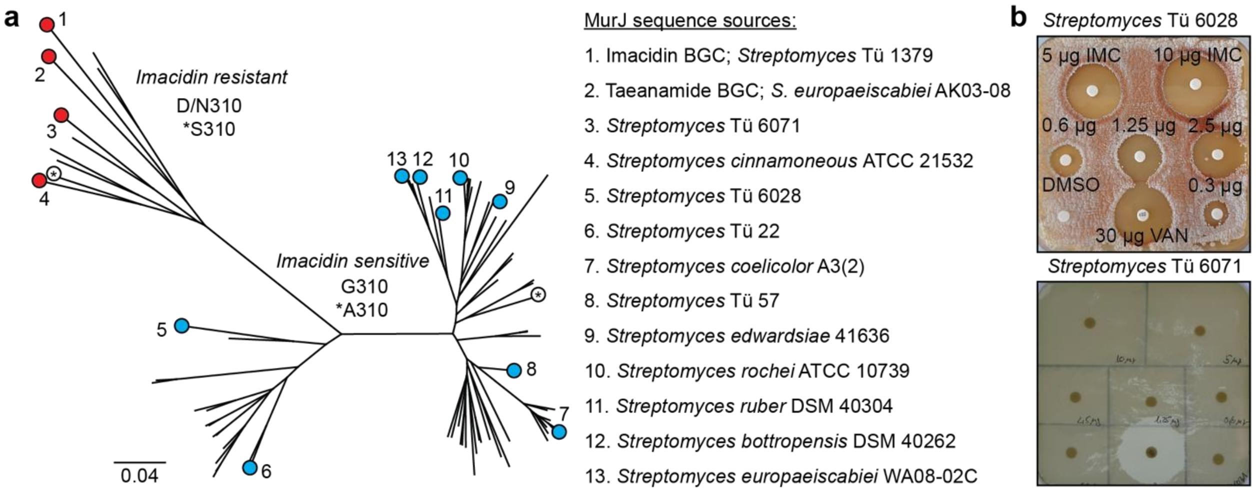


**Figure S15. *Streptomyces* MurJs separate into imacidin-resistant and imacidin-sensitive clades based on the residue at the 310 position.** (a) Imacidin-resistant species contain either an aspartate or asparagine at the 310 position. Imacidin-sensitive species contain either a glycine or alanine at the 310 position. (b) *Streptomyces* Tü 6028 expresses MurJ$^{G310}$ indicating it should be sensitive to imacidin, and agar disk diffusion tests confirm this. *Streptomyces* Tü 6071 expresses MurJ$^{N310}$ indicating it should be resistant to imacidin, and this was verified by agar disk diffusion tests.

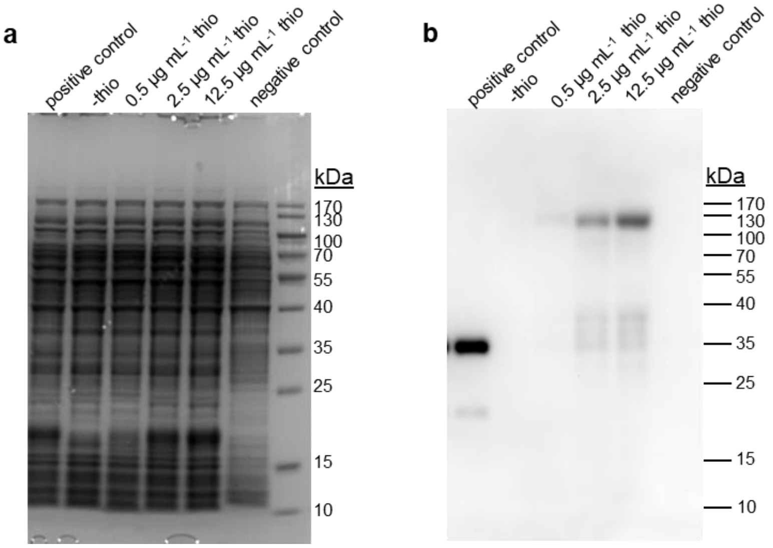


**Figure S16. Inducible expression of mCherry-labelled MurJ in *S. coelicolor* M1154.** Protein expression of mCh-MurJ in *S. coelicolor* pIJ6902-mChMurJ was induced with different concentrations of thiostrepton from the *tipA* promoter in pIJ6902. As a positive control, an extract of *S. coelicolor* pIJ6902-mCh expressing mCherry (26.7 kDa) after induction with 12.5 µg mL$^{-1}$ thiostrepton (thio) was used. As a negative control, an extract of *S. coelicolor* M1154 without any expression plasmid was used. The Coomassie-stained SDS-PAGE (a) is the loading control for the Western blot developed with anti-mCherry (b, and cropped image Fig.4). The Western blot shows increasing expression of mCh-MurJ (112.2 kDa) dependent on the inducer (thiostrepton) concentration. Minor degradation products were detected in the mCh-MurJ expressing samples.

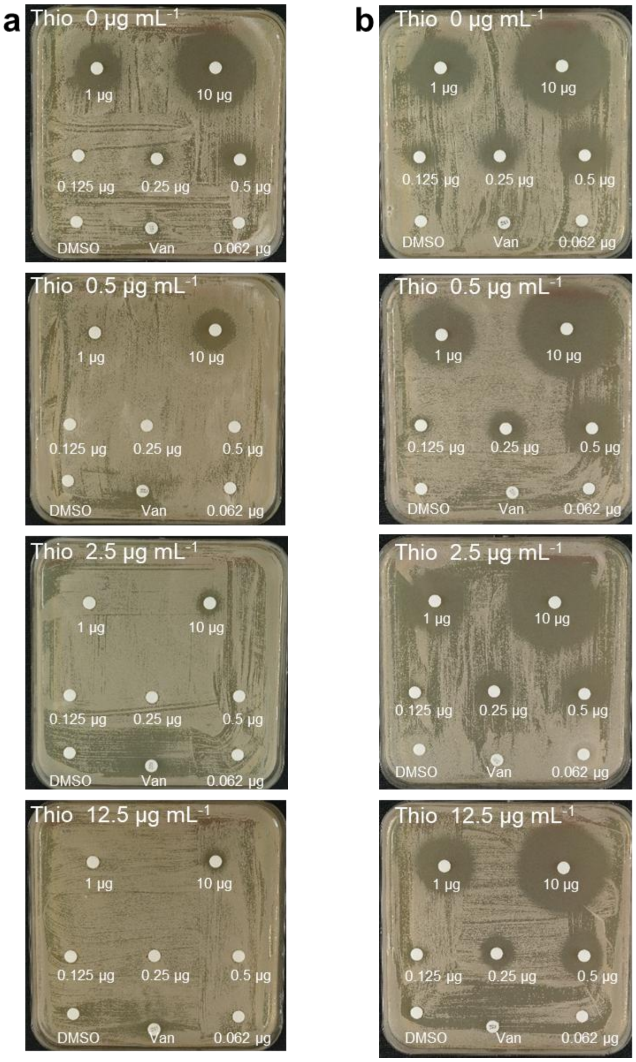


**Figure S17. Increasing induction of mCh-MurJ expression reduces the sensitivity of *S. coelicolor* M1154 pIJ6902-mChMurJ to imacidin.** Disk diffusion assay with *S. coelicolor* M1154 carrying plasmid pIJ6902-mchMurJ (a) or pIJ6902-mch (b) on GYM agar containing the indicated concentration of thiostrepton (Thio) to induce protein expression of mCh-MurJ (a) or mCherry (b). Filter disks contained the indicated amounts of imacidin. As a control, 10 µl of DMSO and 30 µg vancomycin were used. Whereas increasing amounts of thiostrepton reduced the sensitivity of *S. coelicolor* M1154 pIJ6902-mChMurJ against imacidin, thiostrepton did not change the sensitivity of *S. coelicolor* M1154 pIJ6902-mCh to imacidin. Inhibition zones around filter disks with imacidin were smaller even without induction (Thio 0 µg/ $mL^{-1}$) for the pIJ6902-mChMurJ carrying strain compared to the strain harboring plasmid pIJ6902-mCh, probably due to leaky expression of mCh-MurJ from the *tipA* promoter.

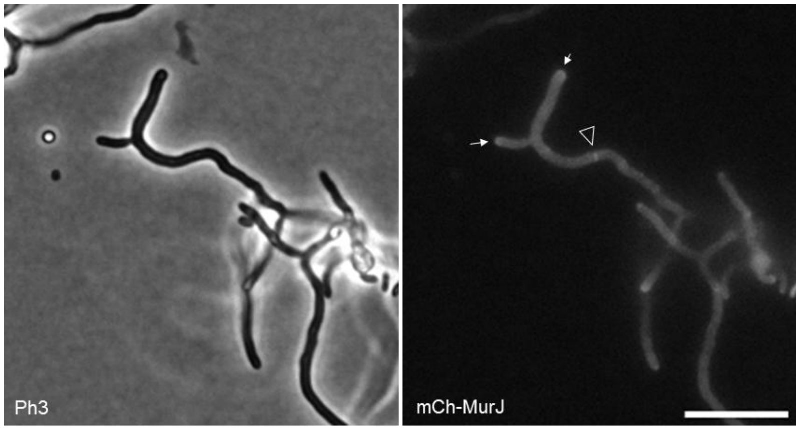


**Figure S18. Localization of mCh-MurJ in *S. coelicolor* to tips and cross-walls.** Fluorescence microscopy reveals that mCh-MurJ localizes to hyphal tips (arrows) and cross-walls (arrowhead). These localization pattern corresponds to the locations of peptidoglycan synthesis during vegetative growth. Micrographs were extracted from a time-lapse with *S. coelicolor* pIJ6902-mChMurJ shown in Movie S3. The growth medium contained a low concentration (0.05 µg $mL^{-1}$) of the inducer thiostrepton. Scale bar: 10 µm.

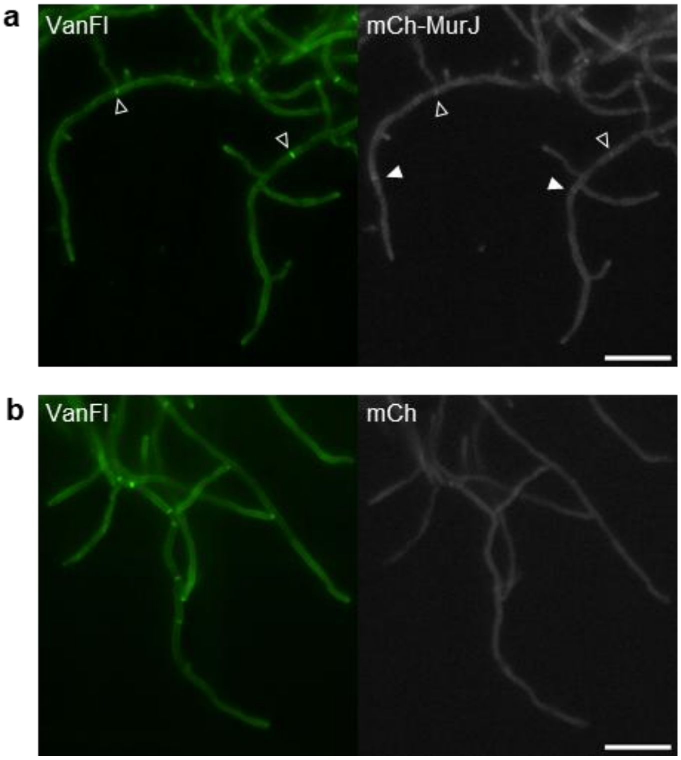


**Figure S19. Fluorescently labelled MurJ localizes to cross walls.** Microscopic analysis of *S. coelicolor* pIJ6902-mChMurJ (a) and *S. coelicolor* pIJ6902-mCh (b) stained with fluorescently labelled vancomycin (VanFl). VanFl stains sites of active peptidoglycan biosynthesis and sites of non-cross-linked peptidoglycan. Expression of mCh-MurJ or mCh was induced by the addition of 0.1 µg $mL^{-1}$ thiostrepton to the GYM medium during growth. mCh-MurJ co-localizes with cross-walls (open arrow). In some apical compartments, mCh-MurJ localizes presumably to newly forming cross-walls without co-localizing Van-Fl staining (filled arrow). This might indicate that mCh-MurJ localizes before the new cross-wall gets visible in the VanFl-staining. When mCherry was expressed without fusion with MurJ, it did not co-localize with the VanFl- stained cross walls but showed a diffusive red signal (b). Scale bar: 10 µm

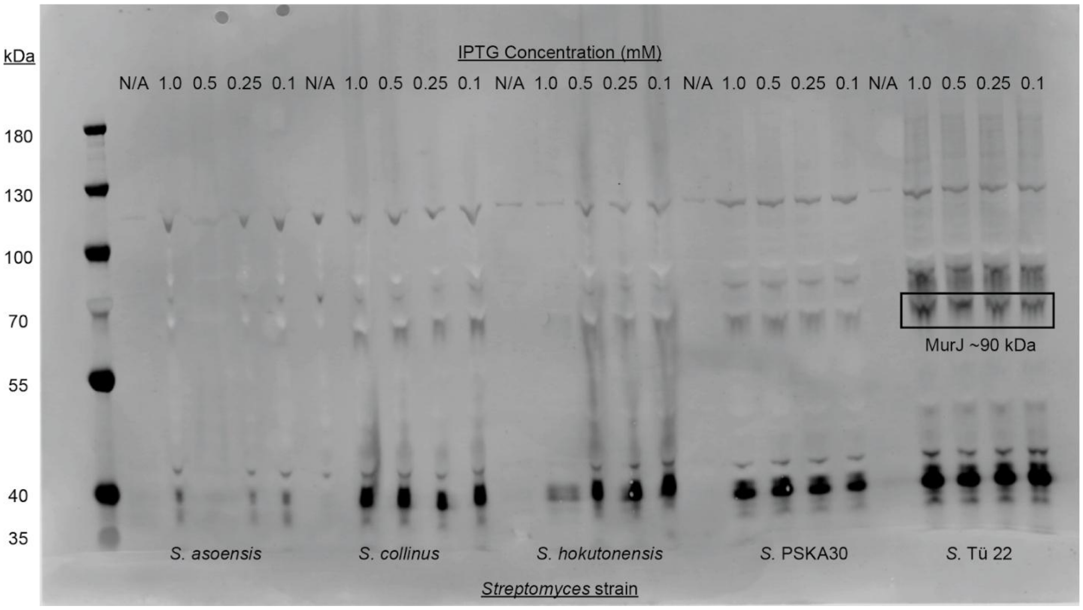


**Figure S20. Candidate MurJs for protein expression.** Expression and purification of the *S. coelicolor* MurJ led to low yield and increased protein degradation. 5 MurJ homologs from imacidin-sensitive *Streptomyces* species (*S. asoensis*, *S. collinus*, *S. hokutonensis*, *S.* PSKA30, *S.* Tü 22) were evaluated for stability in protein purification. Uninduced and induced cell lysates from *Streptomyces* candidates were run on a gel. A Western blot using anti-FLAG and anti-HIS antibodies was used to determine which *Streptomyces* candidate had the best MurJ expression. Based on the blot, *Streptomyces sp.* Tü 22 had the best protein expression and was used for purification.

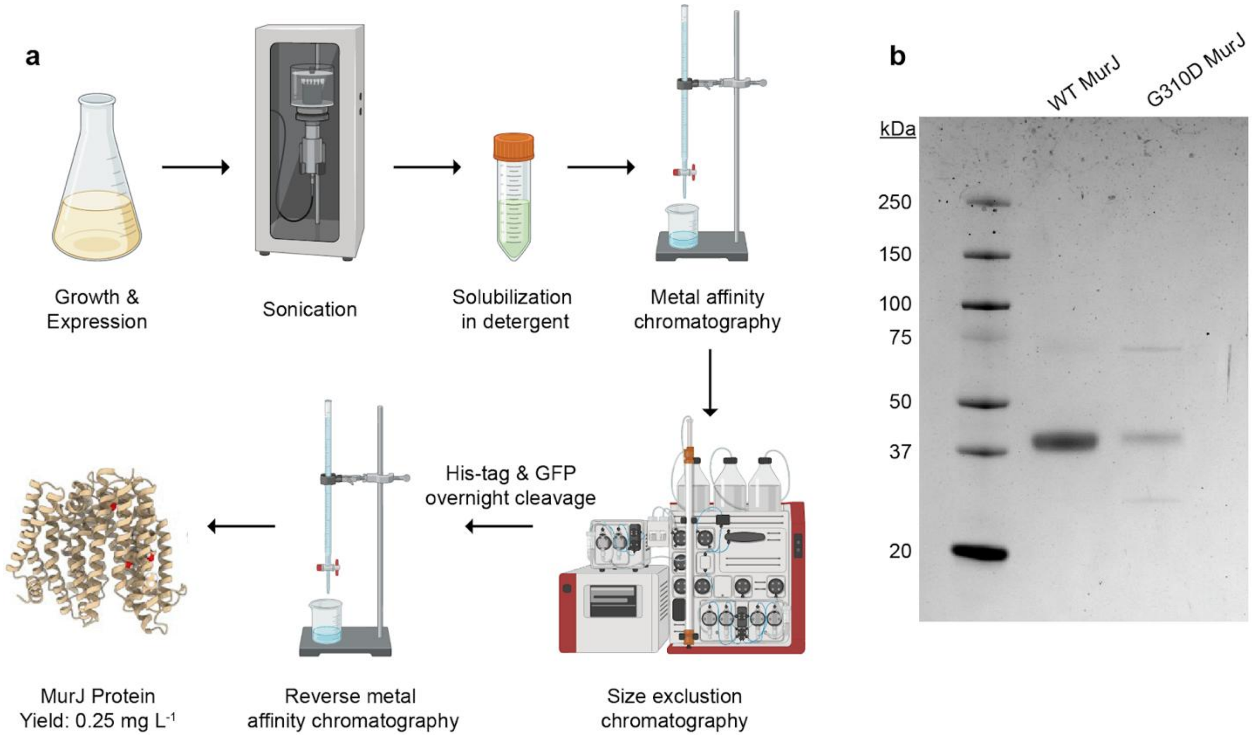


**Figure S21. MurJ protein expression workflow and proteomics.** (a) A simplified workflow of $MurJ^{WT}$ and $MurJ^{G310D}$ purification. BL21-AI *E. coli* transformed with the MurJ expression vectors are grown to an $OD_{600}$ of ~0.7 and induced using IPTG and L-arabinose. The cell lysate is sonicated then solubilized in detergent. After solubilization, the protein is isolated by metal affinity chromatography and size exclusion chromatography. Isolated protein goes through an overnight cleavage to remove the His-tag and sfGFP. To yield the final protein, reverse metal affinity chromatography is used yielding approximately 0.25 mg $L^{-1}$. (b) SDS-PAGE gel (Mini-PROTEAN TGX Stain-Free Precast Gels 4-20% BIO-RAD) of MurJ proteins. $MurJ^{WT}$ is 58.972 kDa and $MurJ^{G310D}$ is 59.030 kDa. Both proteins run at ~40 kDa along the molecular weight ladder (Precision Plus Protein Standards BIO-RAD). This molecular weight is less than expected, but not surprising as it is known that membrane proteins tend to run faster on SDS-PAGE gels. Both $MurJ^{WT}$ and $MurJ^{G310D}$ were confirmed by mass spectrometry proteomics.

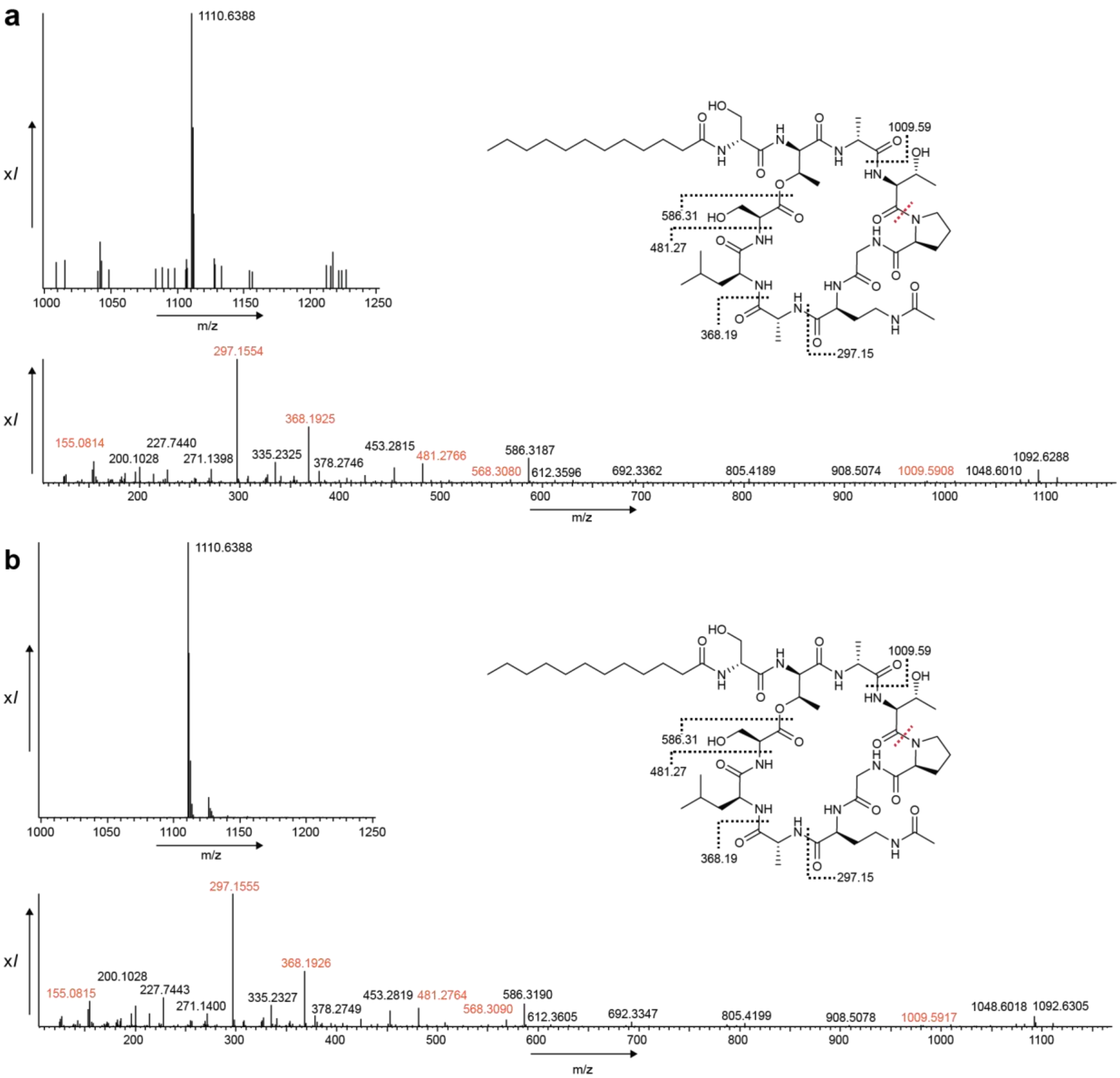


**Figure S22. High resolution mass spectrometry data comparing natural and synthetic taeanamide C.** (a) Taeanamide C detected from cultures of *S. europaeiscabiei* AK08-03 is shown, providing its chemical structure, parent ion (1110.63 [M+H]) and MS/MS spectra, which effectively matches the reported fragmentation pattern for taeanamide A (b) The chemical structure of synthetic taeanamide C is provided, along with its parent ion (1110.63 [M+H]) and MS/MS spectra.

**Movie S1.** ***S. coelicolor*** **M1154 treated with imacidin.** Time-lapse imaging was performed with mycelium of *S. coelicolor* as starting material. The agar pad (1.5 % agarose in GYM) contained 4 x MIC (0.0032 µg $mL^{-1}$ imacidin). The frame rate was 5 min. Hyphae swelled during treatment and lysed. Original hyphal tips did not elongate further, but multiple new branches were formed. Elongation (or swelling) of branches eventually resulted in pronounced lysis indicating loss of cell wall integrity. Scale bar: 10 µm

**Movie S2. Non-treated** ***S. coelicolor*** **M1154.** Time-lapse imaging was performed with mycelium of *S. coelicolor* as starting material on an agar pad (1.5 % agarose in GYM). The frame rate was 5 min. At the beginning of the time-lapse imaging, mycelia needed a lag phase to resume fast growth. In the absence of imacidin, there is only occasional lysis. Scale bar: 10 µm

**Movie S3. Localization of mCh-MurJ to tips and cross walls during growth.** Time-lapse imaging of germinating spores and growing vegetative hyphae of *S. coelicolor* pIJ6902-mChMurJ on an agarose pad in 25 % GYM agar containing a low concentration (0.05 µg $mL^{-1}$) of thiostrepton. Images were taken every 10 min. The first 640 min of the time-lapse imaging were needed for spores to germinate and are not represented in this movie. During growth, mCh-MurJ localizes to extending tips and presumably cross walls. In vegetative hyphae, these are the sites of ongoing active peptidoglycan synthesis. Scale bar: 10 µm

**Table S9. NMR spectroscopic data for imacidin (1)(600 MHz in DMSO-$d_6$)[a,b]**

| Position | $\delta_H$ mult. | $\delta_C$ | Position | $\delta_H$ mult. | $\delta_C$ |
|---|---|---|---|---|---|
| 1 | 0.81 (ov) | 18.9 | 34 | 6.88 (s) | 116.5 |
| 2 | 1.27 (ov) | 33.5 | 35 | 7.59 (ov) | 134.3 |
| 3 | 0.82 (ov) | 11.1 | 36 | - | 170.0 |
| 4 | 1.09 (ov) | 28.7 | 37 | 7.91 (d, 7) | - |
| 5 | 1.25 (ov) | - | 38 | 4.23 (ov) | 48.7 |
| 6 | 1.25 (ov) | - | 39 | 1.24 (ov) | 17.3 |
| 7 | 1.25 (ov) | - | 40 | - | 172.5 |
| 8 | 1.25 (ov) | - | 41 | 8.53 (d, 3.5) | - |
| 9 | 1.25 (ov) | - | 42 | 4.13 (ov) | 58.6 |
| 10 | 1.39, 1.40 (ov) | 36.9 | 43 | 4.97 (q, 6) | 70.5 |
| 11 | 3.71 (br) | 68.3 | 44 | 1.17 (d, 6) | 17.0 |
| 12 | 2.21, 2.25 (dd, 5.5) | 44.6 | 45 | - | 168.9 |
| 13 | - | 170.9 | 46 | 7.63 (ov) | - |
| 14 | 8.30 (d, 3.5) | - | 47 | 4.10 (ov) | 59.3 |
| 15 | 4.39 (dd, 5.6) | 50.4 | 48 | 3.81 (ov) | 66.1 |
| 16 | 2.83, 3.09 (dd, 6) | 50.1 | 49 | 0.91 (d, 6) | 19.2 |
| 17 | - | 170.6 | 50 | - | 172.0 |
| 18 | 7.85 (d, 8) | - | 51 | 7.95 (d, 7) | - |
| 19 | 4.09 (ov) | 51.1 | 52 | 4.46 (br) | 52.7 |
| 20 | 1.51, 1.57 (ov) | 39.3 | 53 | 2.83, 2.93 (dd, 6.5) | 29.0 |
| 21 | 1.53 (ov) | 23.5 | 54 | - | 133.2 |
| 22 | 0.82 (ov) | 23.1 | 55 | 6.84 (s) | 117.0 |
| 23 | 0.75 (d, 6) | 20.7 | 56 | 7.58 (ov) | 134.3 |
| 24 | - | 171.6 | 57 | - | 171.5 |
| 25 | 7.99 (t, 4.4) | - | 58 | 8.25 (br) | - |
| 26 | 2.86, 2.94 (m, 6) | 37.9 | 59 | 4.18 (q, 6.7) | 48.9 |
| 27 | 1.50, 1.53 (ov) | 25.2 | 60 | 1.30 (d, 7) | 16.7 |
| 28 | 2.01, 2.07 (m, 7) | 32.4 | 61 | - | 171.7 |
| 29 | - | 171.8 | 62 | 7.49 (d, 9.5) | - |
| 30 | 9.03 (d, 3.5) | - | 63 | 4.29 (dd, 5.6) | 58.1 |
| 31 | 3.82 (br) | 54.9 | 64 | 4.21 (ov) | 65.8 |
| 32 | 3.05, 3.21 (d, 13) | 25.6 | 65 | 1.06 (d, 6) | 19.3 |
| 33 | - | 134.7 | 66 | - | 169 .1 |

[a]Chemical shift □ and (multiplicity, *J* in Hz).
[b]Aliased carbonyl HMBC recorded on an 800 MHz spectrometer

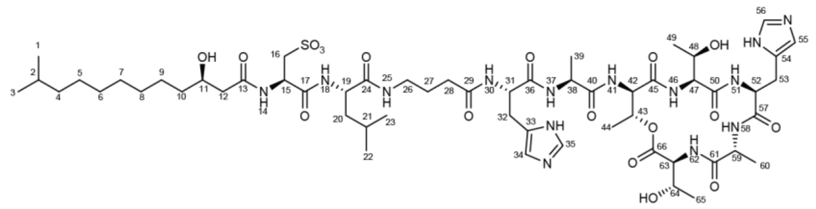

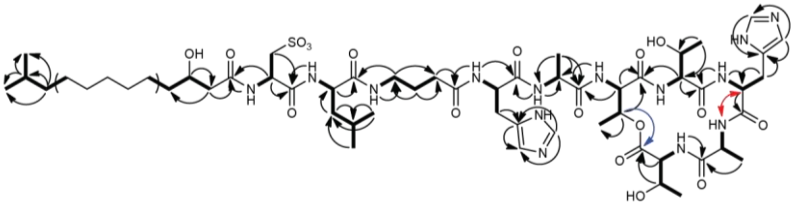

**$^1$H NMR spectrum of imacidin (1) in DMSO-$d_6$.**

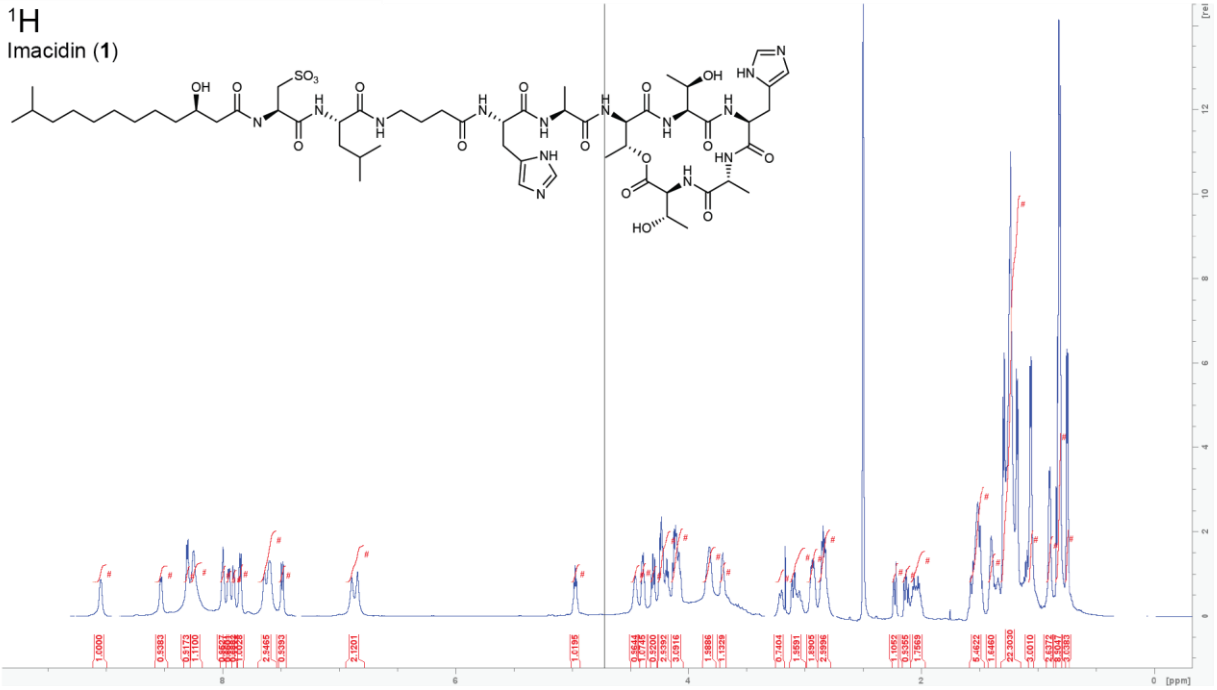


**$^1$H-$^1$H COSY NMR spectrum of imacidin (1) in DMSO-$d_6$.**

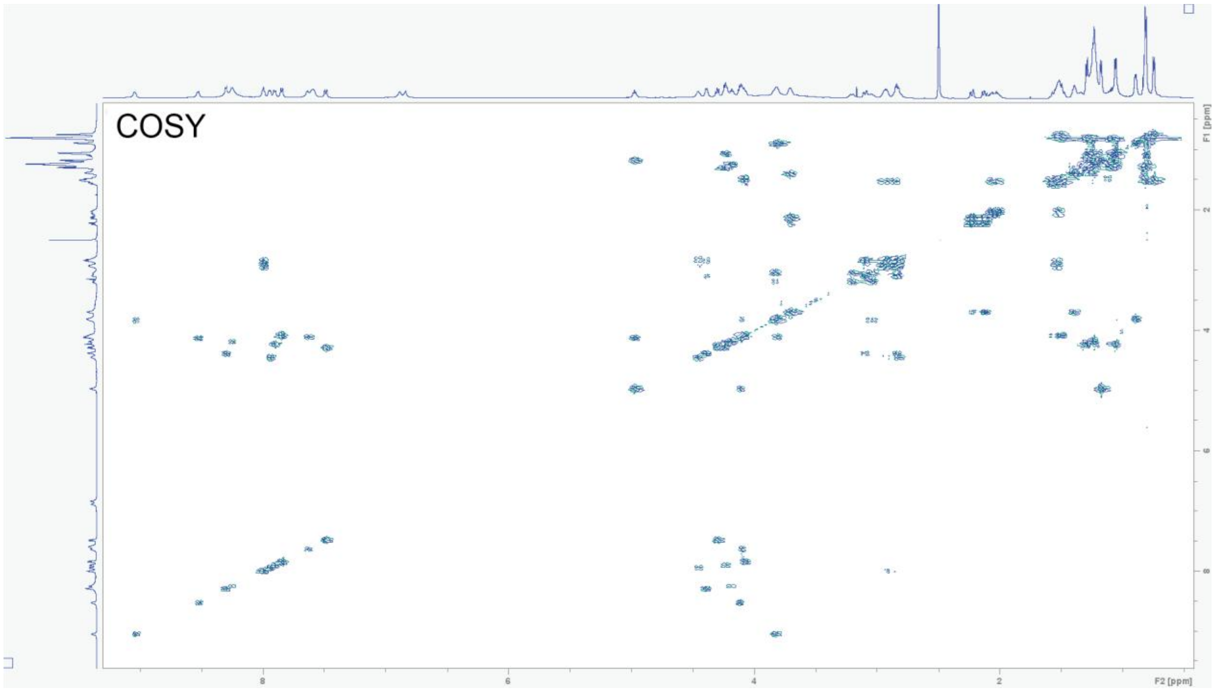

**$^{1}$H-$^{1}$H NOESY NMR spectrum of imacidin (1) in DMSO-$d_6$.**

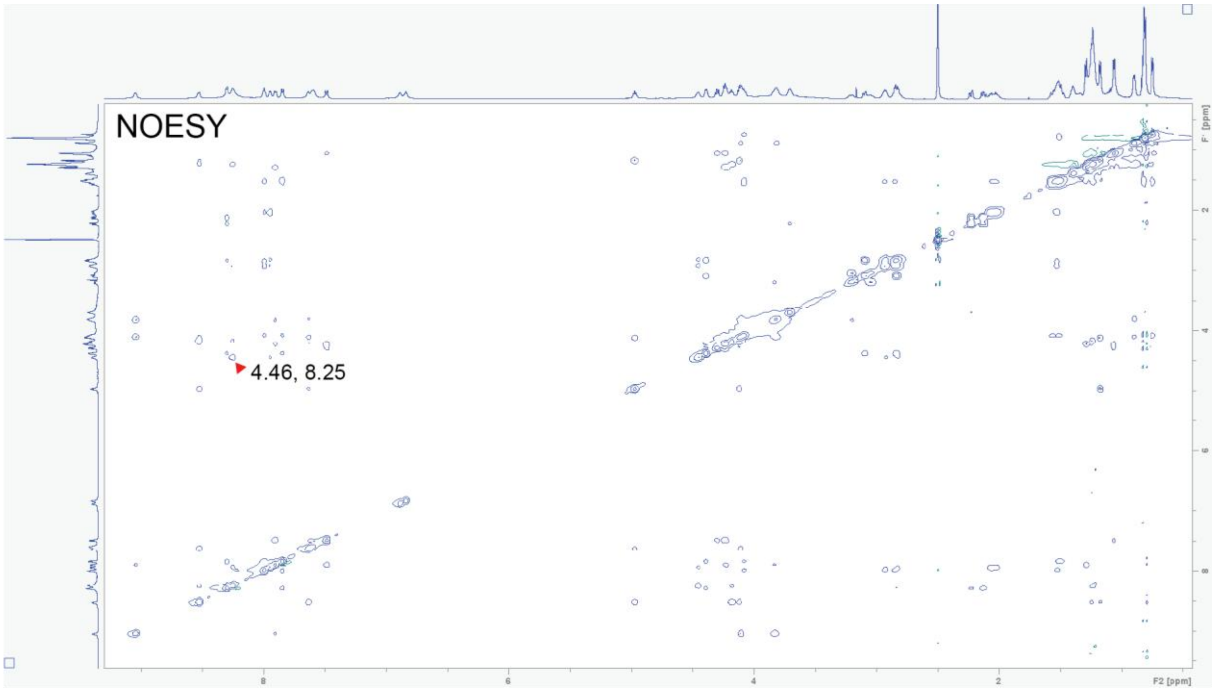


**$^{1}$H-$^{13}$C HSQC NMR spectrum of imacidin (1) in DMSO-$d_6$.**

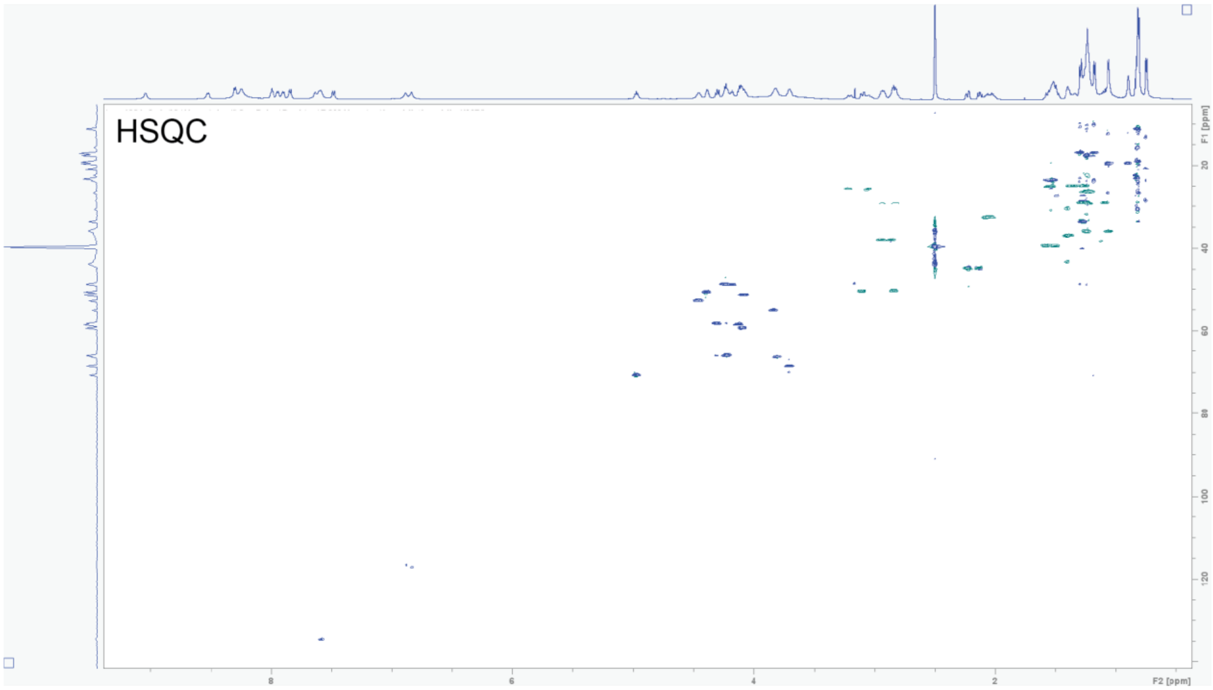

**$^1$H-$^{13}$C HMBC NMR spectrum of imacidin (1) in DMSO-$d_6$.**

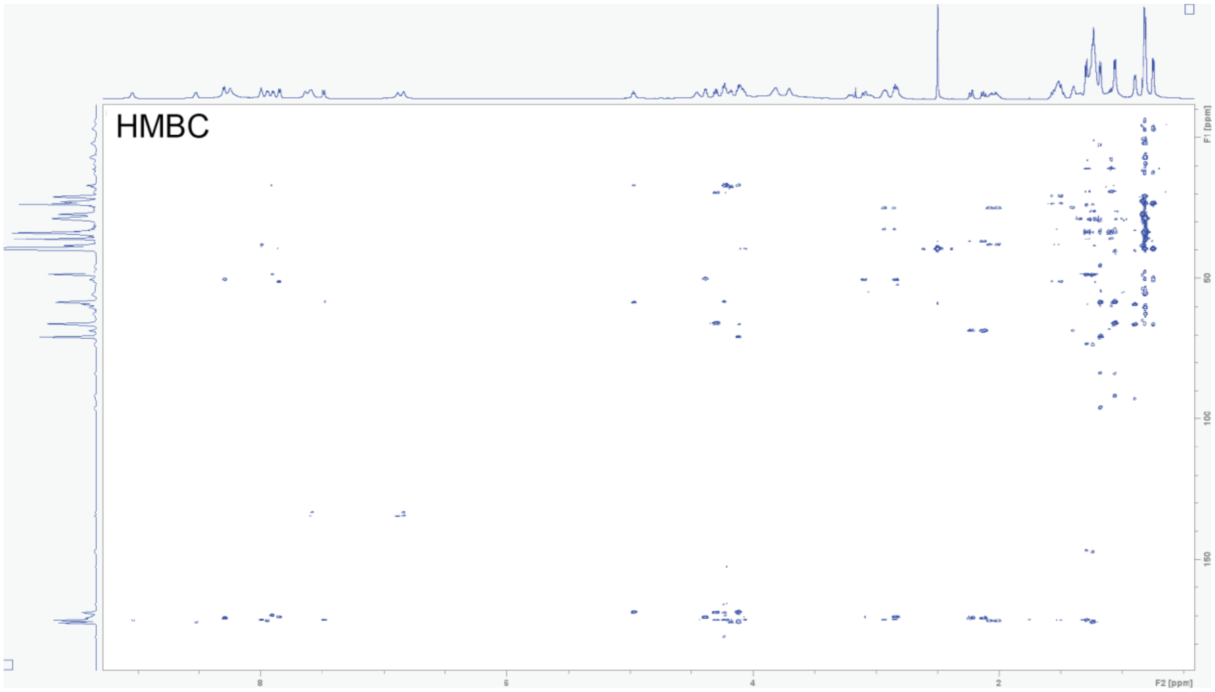


**$^1$H-$^{13}$C aliased HMBC NMR spectrum of imacidin (1) in DMSO-$d_6$**

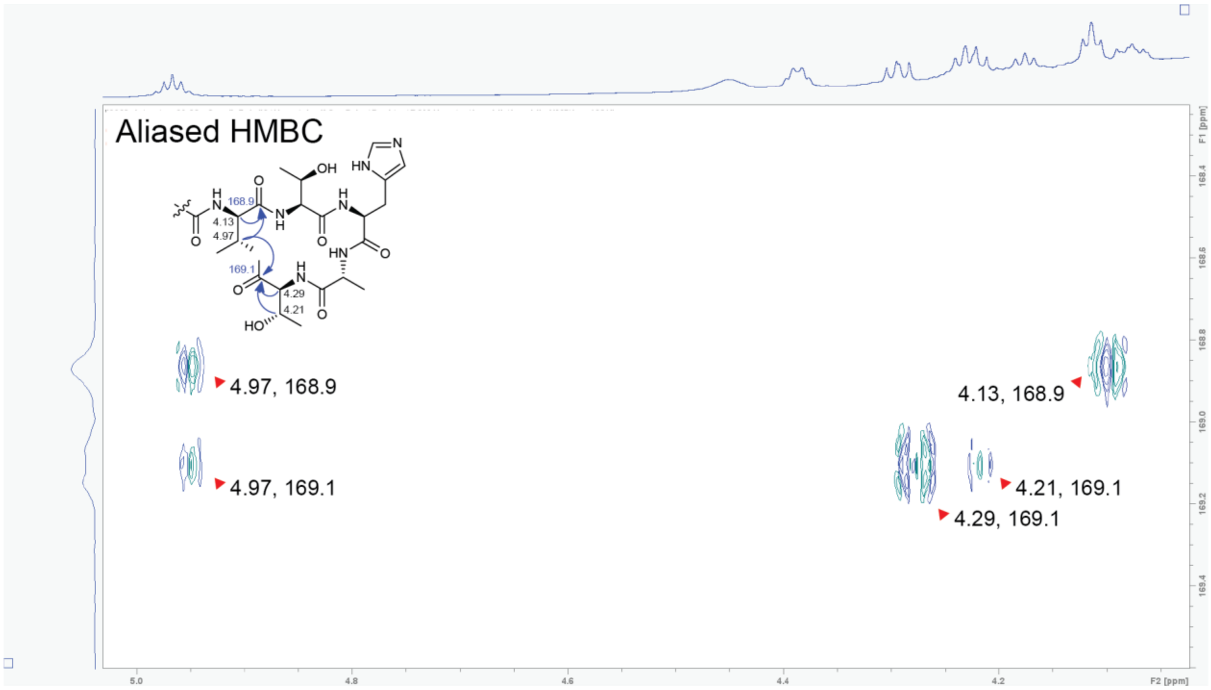

**Table S10. NMR spectroscopic data for imacidin 1231 (2)(600 MHz in DMSO-$d_6$)[a]**

| Position | $\delta_H$ mult. | $\delta_C$ | Position | $\delta_H$ mult. | $\delta_C$ |
|---|---|---|---|---|---|
| 1 | 0.83 (ov) | 22.3 | 34 | 7.42 (s) | 116.8 |
| 2 | 1.48 (ov) | 27.9 | 35 | 9.00 (s) | 133.3 |
| 3 | 0.84 (ov) | 22.3 | 36 | - | 169.2 |
| 4 | 1.12, 1.13 (ov) | 38.2 | 37 | 8.11 (ov) | - |
| 5 | 1.22 (ov) | 29.0 | 38 | 4.20 (br) | 49.0 |
| 6 | 1.22 (ov) | 29.0 | 39 | 1.27 (d, 7) | 17.4 |
| 7 | 1.22 (ov) | 29.0 | 40 | - | 172.6 |
| 8 | 1.23 (ov) | 24.9 | 41 | 8.41 (ov) | - |
| 9 | 1.35 (ov) | 36.6 | 42 | 4.11 (t, 6) | 58.3 |
| 10 | 3.76 (q, 6) | 67.7 | 43 | 4.97 (q, 6) | 70.4 |
| 11 | 2.19, 2.20 (d, 6.5) | 43.3 | 44 | 1.18 (d, 6) | 16.4 |
| 12 | - | 171.5 | 45 | - | 168.2 |
| 13 | 8.22 (br) | - | 46 | 7.41 (ov) | - |
| 14 | 4.50 (dd, 7) | 49.5 | 47 | 4.28 (br) | 58.5 |
| 15 | 2.55, 2.64 (dd, 5.5) | 35.7 | 48 | 3.81 (q, 6) | 66.8 |
| 16 | - | 171.7 | 49 | 0.89 (d, 6) | 19.4 |
| 17 | - | 170.6 | 50 | - | 172.0 |
| 18 | 7.94 (d, 8) | - | 51 | 8.13 (ov) | - |
| 19 | 4.16 (br) | 51.0 | 52 | 4.59 (dd, 7) | 51.0 |
| 20 | 1.43, 1.47 (ov) | 40.4 | 53 | 2.90, 3.04 (ov) | 26.6 |
| 21 | 1.55 (ov) | 23.8 | 54 | - | 129.4 |
| 22 | 0.79 (d, 6) | 21.0 | 55 | 7.39 (s) | 116.9 |
| 23 | 0.84 (ov) | 22.8 | 56 | 8.96 (s) | 133.4 |
| 24 | - | 171.7 | 57 | - | 170.4 |
| 25 | 7.76 (t, 5.5) | - | 58 | 8.38 (ov) | - |
| 26 | 2.92, 3.00 (m, 6.5) | 37.8 | 59 | 4.19 (br) | 48.8 |
| 27 | 1.55, 1.56 (m, 7) | 24.8 | 60 | 1.28 (d, 7) | 16.8 |
| 28 | 2.08 (t, 7) | 32.1 | 61 | - | 171.4 |
| 29 | - | 172.0 | 62 | 7.31 (br) | - |
| 30 | 9.18 (br) | - | 63 | 4.26 (br) | 58.1 |
| 31 | 4.01 (br) | 53.2 | 64 | 4.23 (br) | 66.1 |
| 32 | 3.12, 3.38 (br) | 23.9 | 65 | 1.07 (d, 6) | 19.4 |
| 33 | - | 130.6 | 66 | - | 168.9 |

[a]Chemical shift □ and (multiplicity, *J* in Hz).

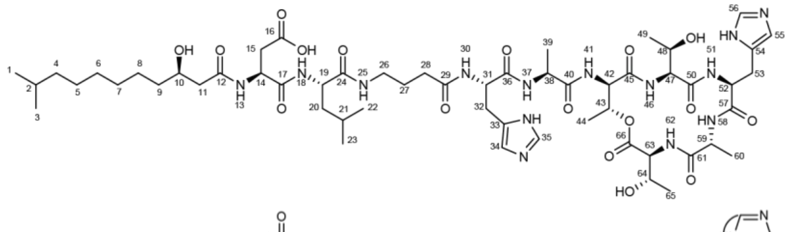

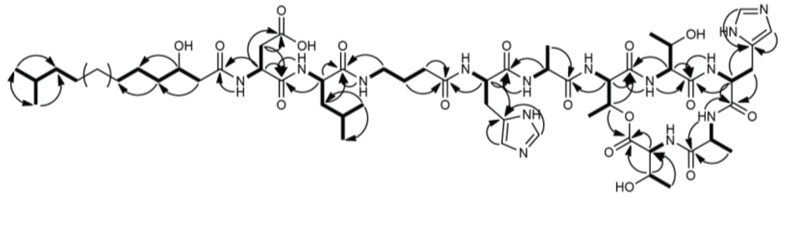

$^{1}H$ NMR spectrum of imacidin 1231 (2) in DMSO-$d_6$.

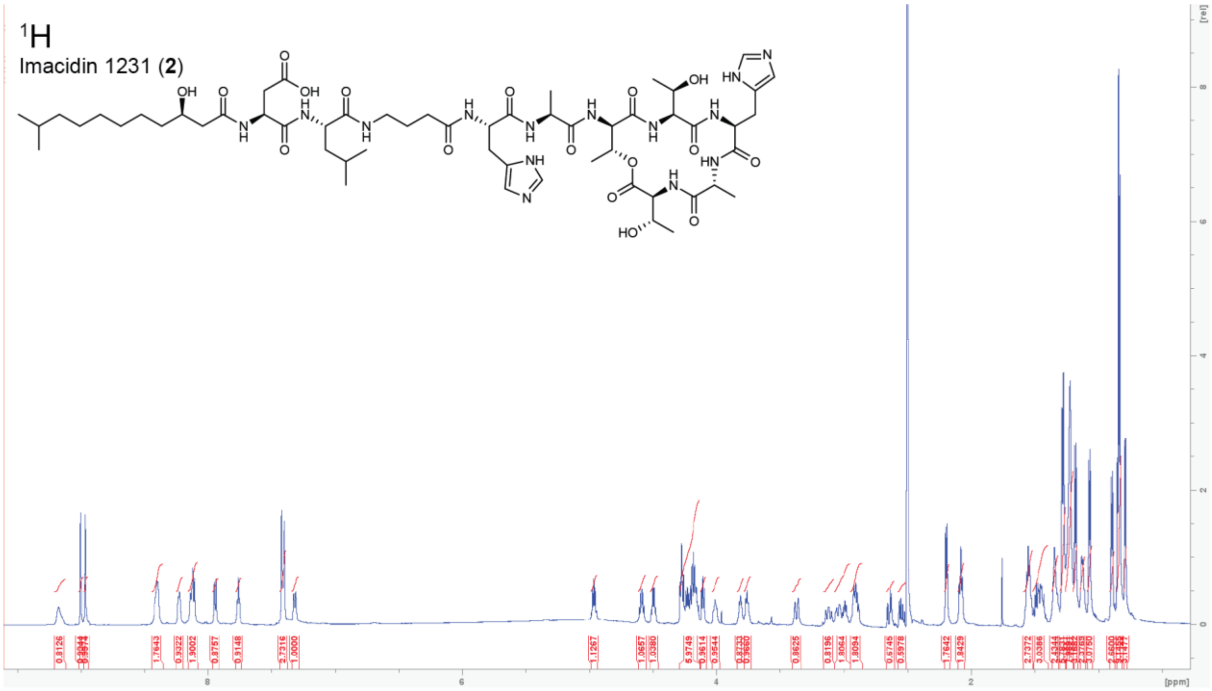


$^{1}H$-$^{1}H$ COSY NMR spectrum of imacidin 1231 (2) in DMSO-$d_6$.

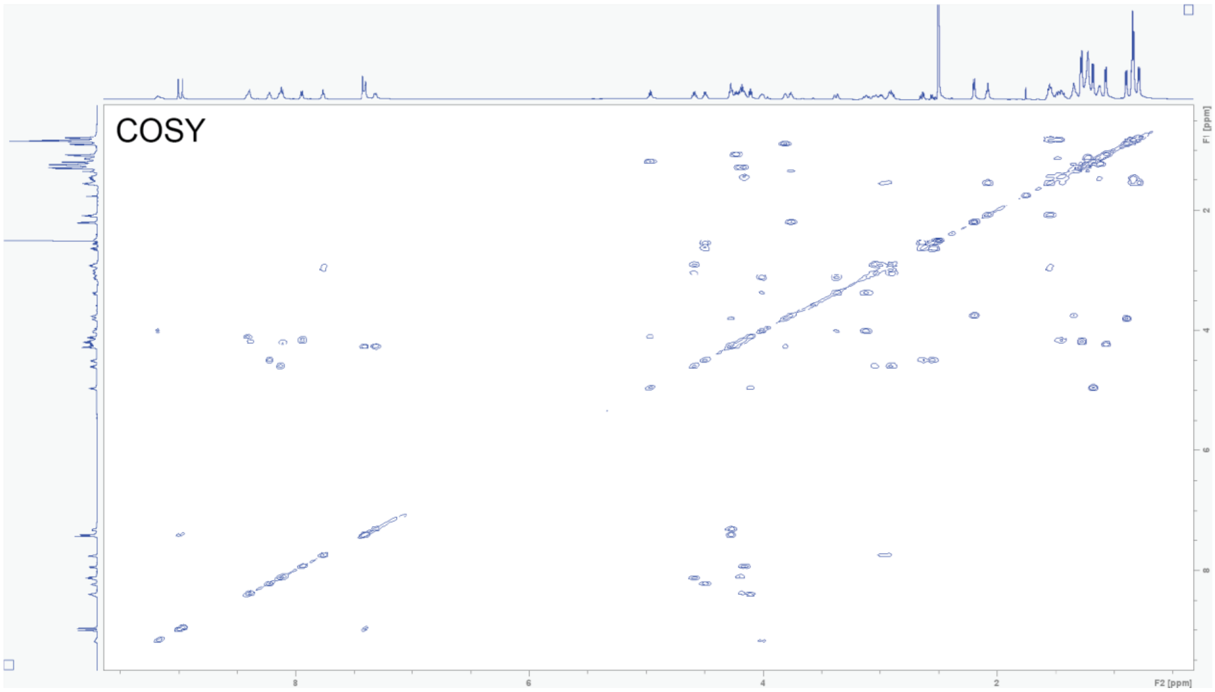

**$^{1}H$-$^{13}C$ HSQC NMR spectrum of imacidin 1231 (2) in DMSO-$d_6$.**

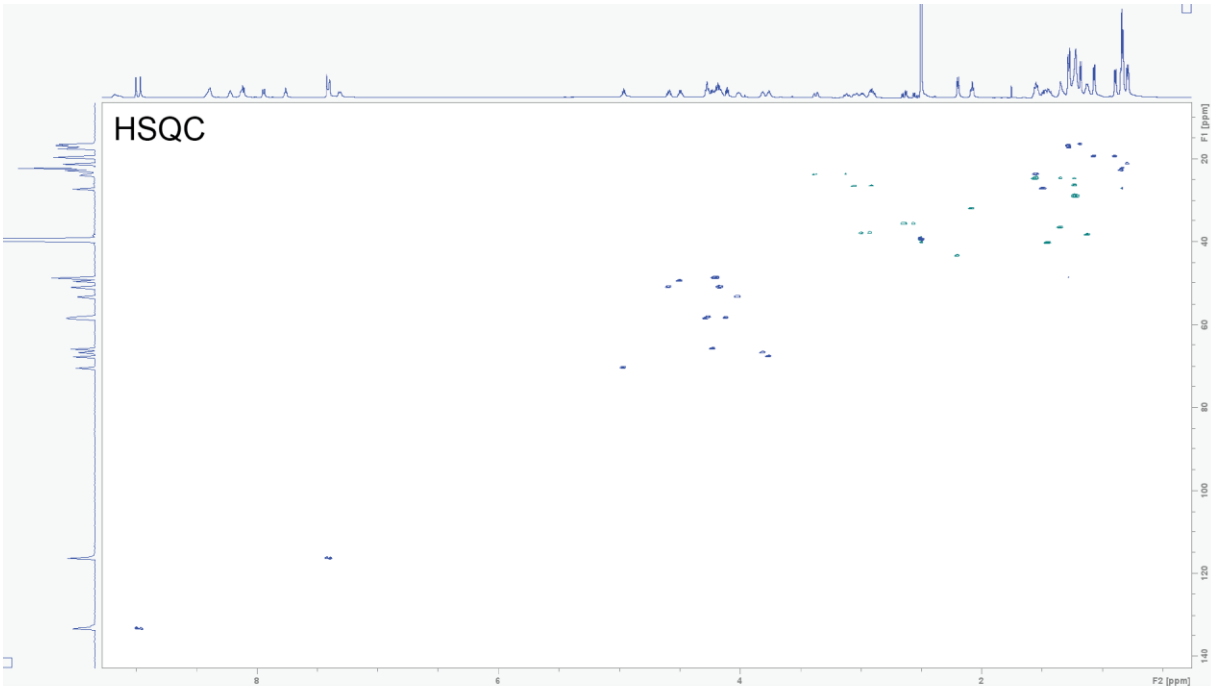


**$^{1}H$-$^{13}C$ HMBC NMR spectrum of imacidin 1231 (2) in DMSO-$d_6$.**

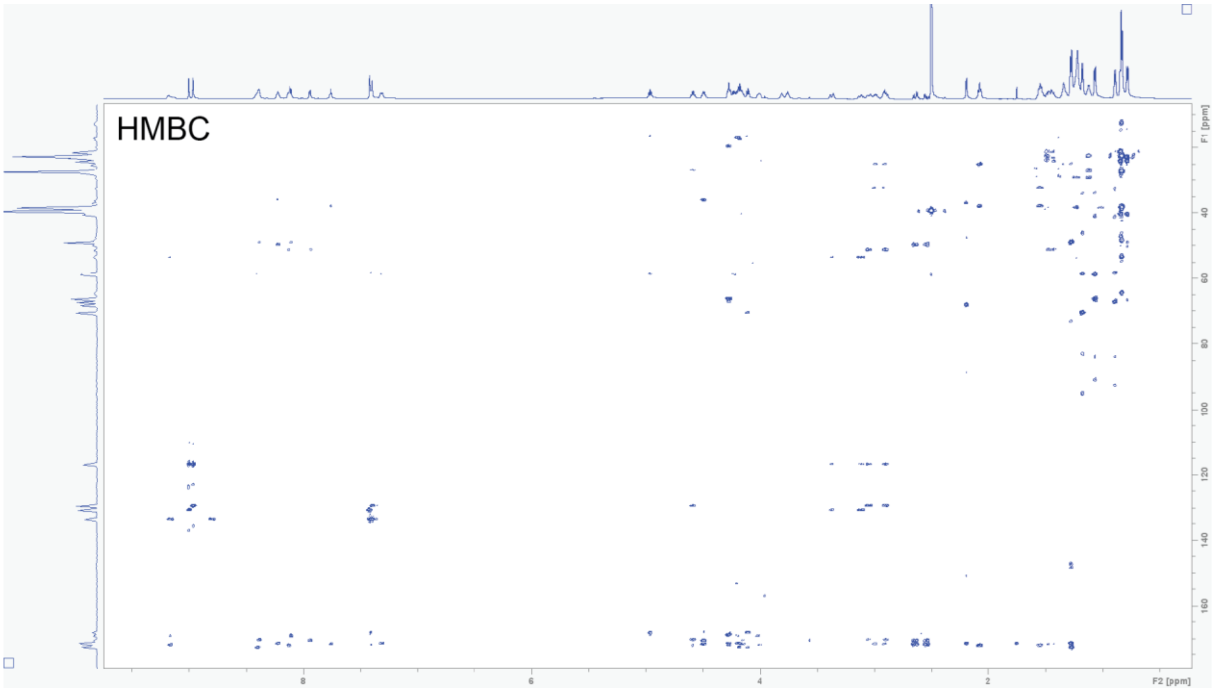

**Table S11. NMR spectroscopic data for imacidin 1245 (3)(600 MHz in DMSO-$d_6$)[a]**

| Position | $\delta_H$ mult. | $\delta_C$ | Position | $\delta_H$ mult. | $\delta_C$ |
|---|---|---|---|---|---|
| 1 | 0.85 (d, 6.5) | 23.0 | 35 | 6.86 (s) | - |
| 2 | 1.48 (ov) | 27.9 | 36 | 7.55 (s) | 135.1 |
| 3 | 0.84 (d, 6.5) | 23.0 | 37 | - | 171.9 |
| 4 | 1.13 (ov) | 38.9 | 38 | 8.37 (br) | - |
| 5 | 1.24 (ov) | 29.7 | 39 | 4.20 (ov) | 49.5 |
| 6 | 1.24 (ov) | 29.7 | 40 | 1.23 (ov) | 18.2 |
| 7 | 1.24 (ov) | 29.7 | 41 | - | 173.0 |
| 8 | 1.24 (ov) | 29.7 | 42 | 8.96 (s) | - |
| 9 | 1.35, 1.36 (ov) | 37.3 | 43 | 4.12 (br) | 59.1 |
| 10 | 3.76 (m, 6.5) | 68.3 | 44 | 4.97 (q, 6) | 71.1 |
| 11 | 2.22 (d, 6) | 44.1 | 45 | 1.16 (d, 6) | 17.5 |
| 12 | - | 171.9 | 46 | - | 169.6 |
| 13 | 8.22 (br) | - | 47 | 7.99 (ov) | - |
| 14 | 4.21 (ov) | 53.0 | 48 | 4.02 (br) | 60.3 |
| 15 | 1.77, 1.85 (ov) | 27.9 | 49 | 3.79 (m, 6) | 66.7 |
| 16 | 2.18 (m, 6.5) | 31.4 | 50 | 0.88 (ov) | 19.9 |
| 17 | - | - | 51 | - | - |
| 18 | - | - | 52 | 8.04 (br) | - |
| 19 | 8.14 (br) | - | 53 | 4.45 (br) | 53.2 |
| 20 | 4.20 (ov) | 51.0 | 54 | 2.79, 2.94 (dd, 7.7) | 29.8 |
| 21 | 1.47 (ov) | 41.0 | 55 | - | - |
| 22 | 1.55 (ov) | 24.6 | 56 | 6.79 (s) | - |
| 23 | 0.86 (ov) | 23.5 | 57 | 7.52 (s) | 135.0 |
| 24 | 0.81 (d, 6.5) | 21.8 | 58 | - | 170.4 |
| 25 | - | - | 59 | 7.90 (ov) | - |
| 26 | 7.88 (ov) | - | 60 | 4.23 (m, 7) | 49.2 |
| 27 | 2.94, 2.99 (ov) | 38.6 | 61 | 1.31 (d, 7) | 17.5 |
| 28 | 1.55 (ov) | 25.6 | 62 | - | 172.1 |
| 29 | 2.07 (m, 6) | 33.1 | 63 | 7.49 (d, 9.7) | - |
| 30 | - | 172.2 | 64 | 4.33 (dd, 5.5) | 58.5 |
| 31 | 9.11 (s) | - | 65 | 4.21 (ov) | 66.4 |
| 32 | 3.85 (br) | 55.7 | 66 | 1.05 (d, 6) | 19.8 |
| 33 | 3.06, 3.21 (br) | 26.4 | 67 | - | 169.5 |
| 34 | - | - | | | |

[a]Chemical shift  and (multiplicity, *J* in Hz).

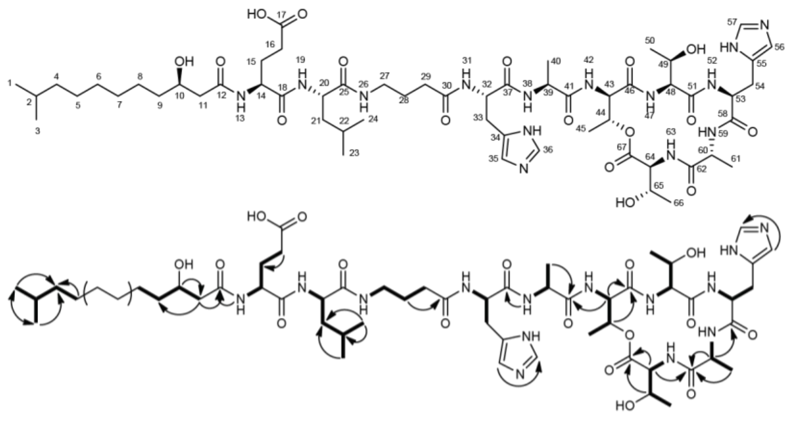

**$^{1}H$ NMR spectrum of imacidin 1245 (3) in DMSO-$d_6$.**

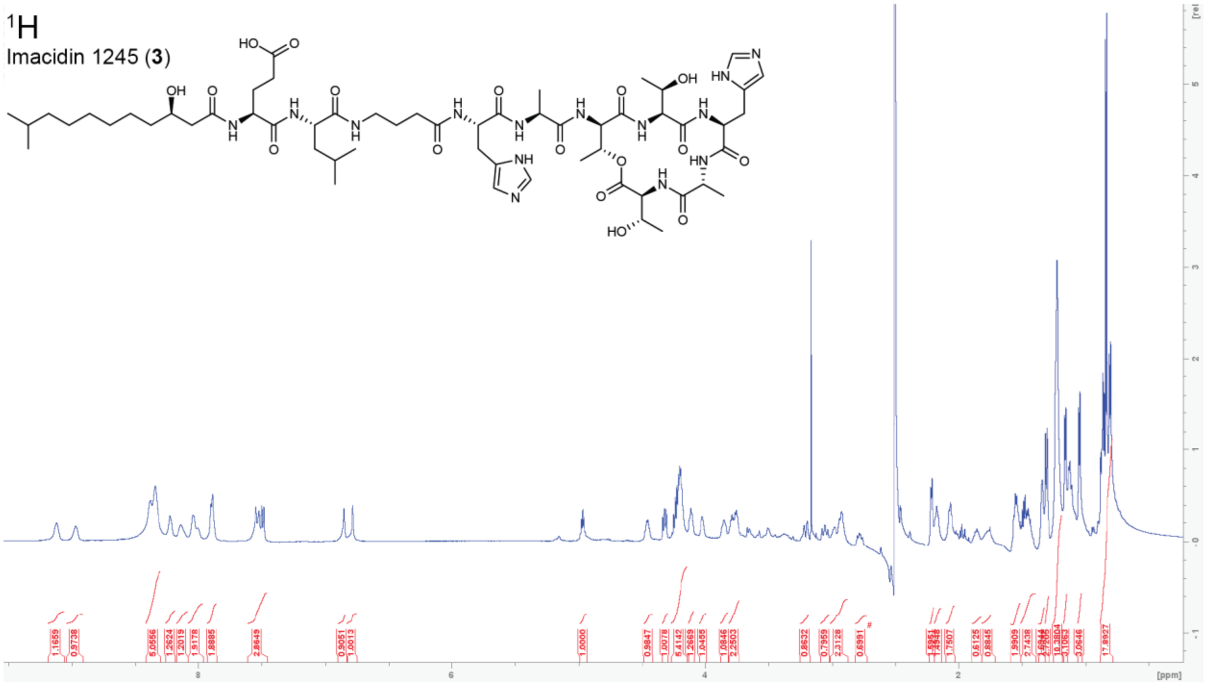


**$^{1}H$-$^{1}H$ COSY NMR spectrum of imacidin 1245 (3) in DMSO-$d_6$.**

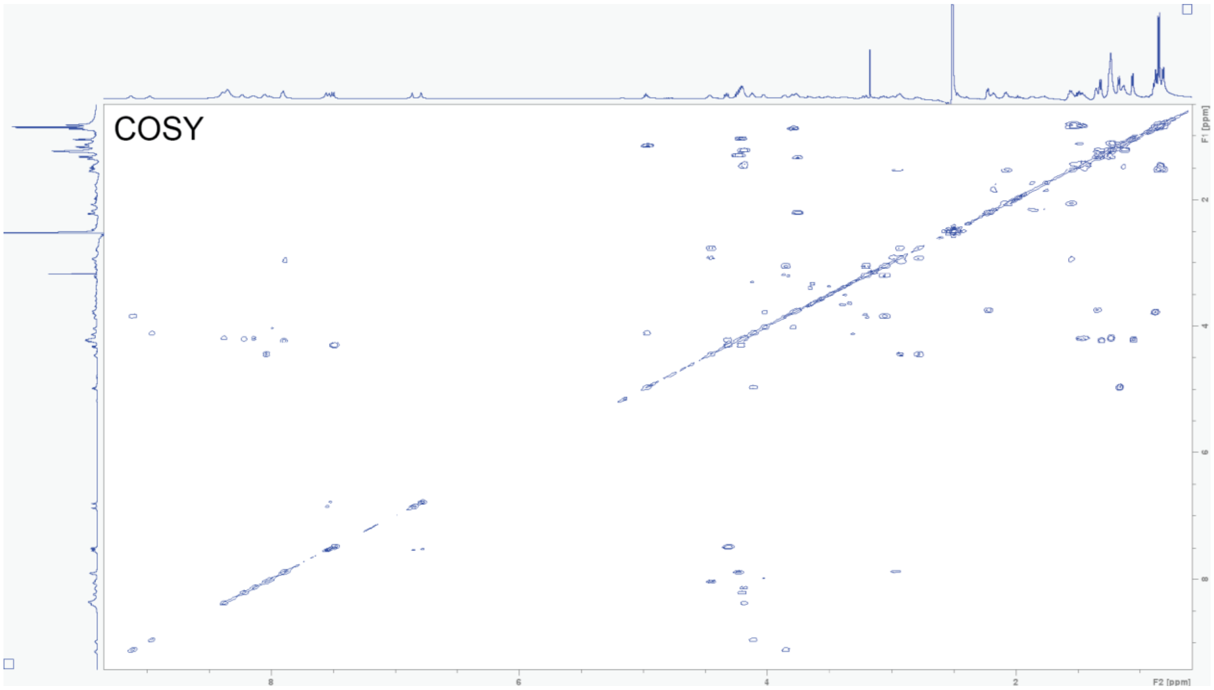

**$^{1}$H-$^{13}$C HSQC NMR spectrum of imacidin 1245 (3) in DMSO-$d_6$.**

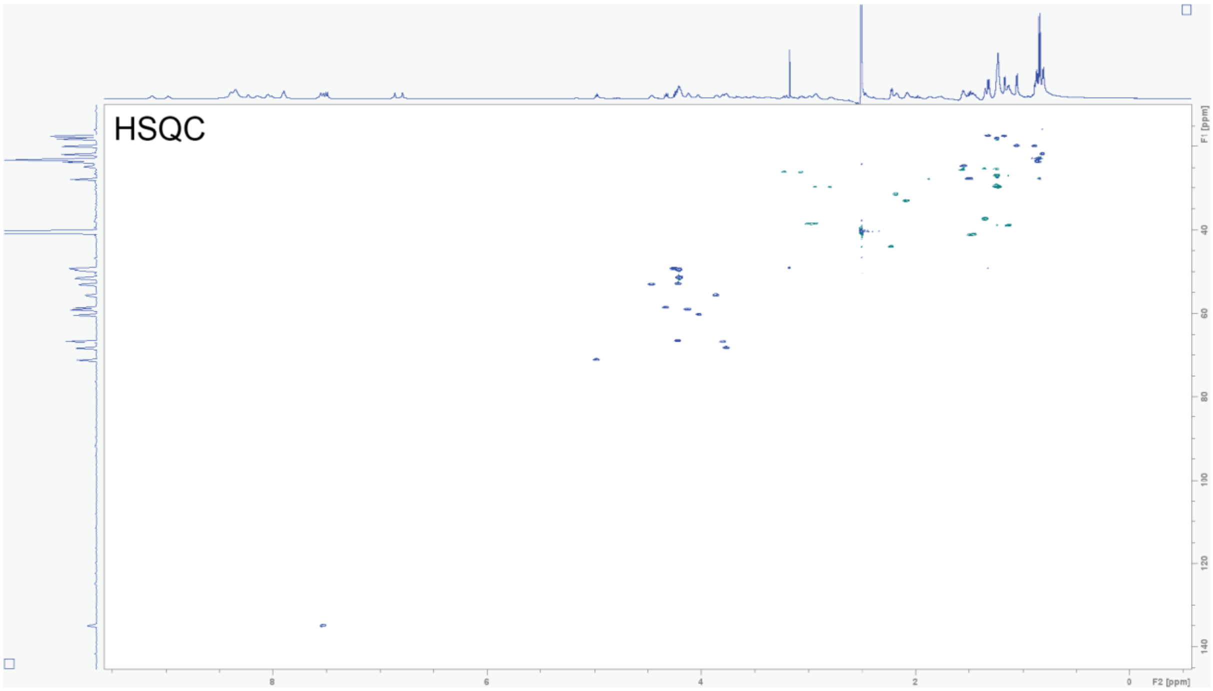


**$^{1}$H-$^{13}$C HMBC NMR spectrum of imacidin 1245 (3) in DMSO-$d_6$.**

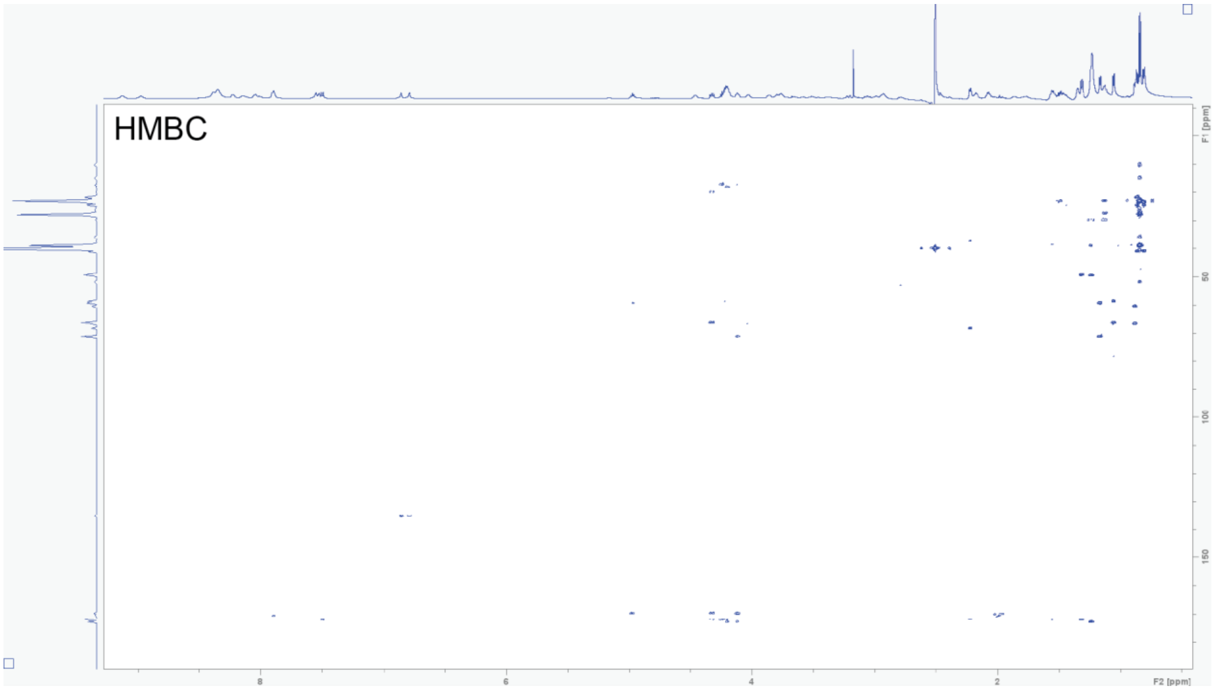

**Table S12. NMR spectroscopic data for N-Fmoc-D-allo-Thr(N-Boc-L-allo-Thr(tBu))-OH (4)(600 MHz in DMSO-$d_6$)[a, b]**

| Position | $\delta_H$ mult. | $\delta_C$ | Position | $\delta_H$ mult. | $\delta_C$ |
|---|---|---|---|---|---|
| 1 | 1.39 (s) | 28.5 | 13 | 4.40 (dd, 4.4) | 57.3 |
| 2 | - | 78.5 | 14 | - | 170.9 |
| 3 | - | 155.9 | 15 | 7.83 (ov) | - |
| 4 | 6.67 (d, 9) | - | 16 | - | 156.6 |
| 5 | 4.08 (dd, 5) | 59.6 | 17 | 4.30 (br) | 66.6 |
| 6 | - | 172.8 | 18 | 4.24 (dd, 7) | 47.1 |
| 7 | 3.96 (q, 6) | 67.3 | 19 | - | 144.2 |
| 8 | 1.06 (ov) | 19.4 | 20 | 7.74 (dd, 3.5) | 125.7 |
| 9 | - | 73.8 | 21 | 7.33 (t, 7.5) | 127.4 |
| 10 | 1.10 (s) | 28.4 | 22 | 7.41 (t, 7.4) | 128.0 |
| 11 | 5.17 (q, 5.5) | 70.8 | 23 | 7.87 (d, 7.5) | 120.4 |
| 12 | 1.26 (br) | 16.0 | 24 | - | 141.2 |

[a]Chemical shift □ and (multiplicity, *J* in Hz).
[b]DMF contamination is visible in the spectra ($^1$H 2.74, 2.88, 7.96; $^{13}$C 31.0, 36.1, 162.7)

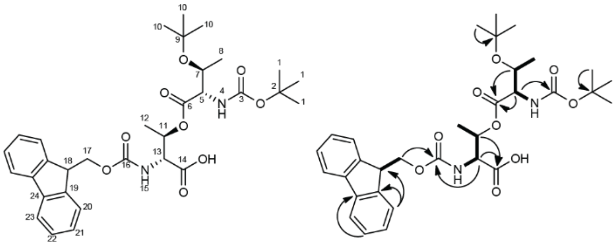

**$^{1}$H NMR spectrum of N-Fmoc-D-allo-Thr(N-Boc-L-allo-Thr(tBu))-OH (4) in DMSO-$d_6$.**

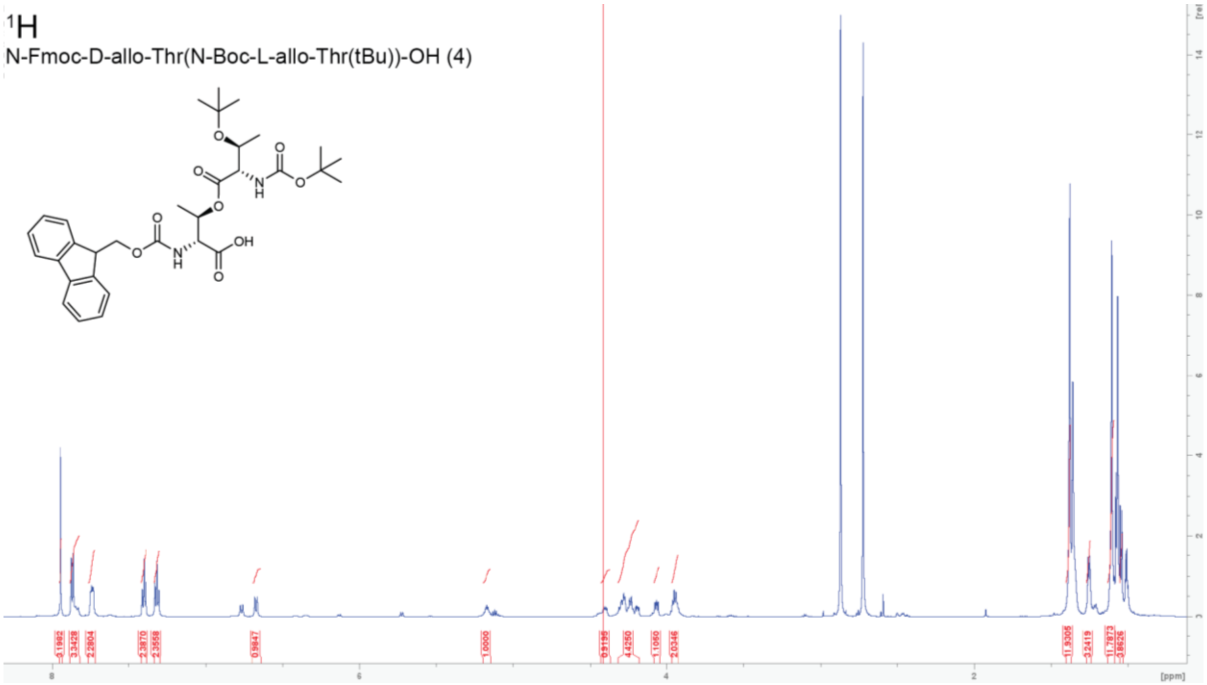


**$^{1}$H-$^{1}$H COSY NMR spectrum of N-Fmoc-D-allo-Thr(N-Boc-L-allo-Thr(tBu))-OH (4) in DMSO-$d_6$.**

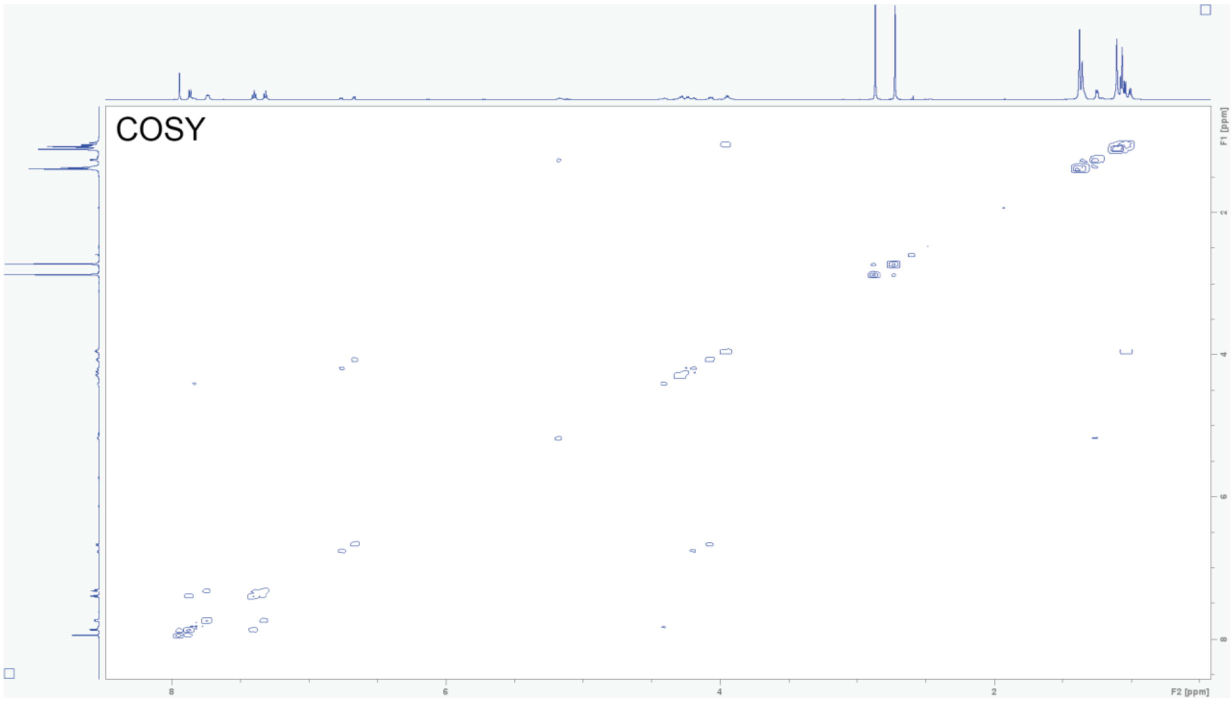

**$^1$H-$^{13}$C HSQC NMR spectrum of N-Fmoc-D-allo-Thr(N-Boc-L-allo-Thr(tBu))-OH (4) in DMSO-*d*$_6$.**

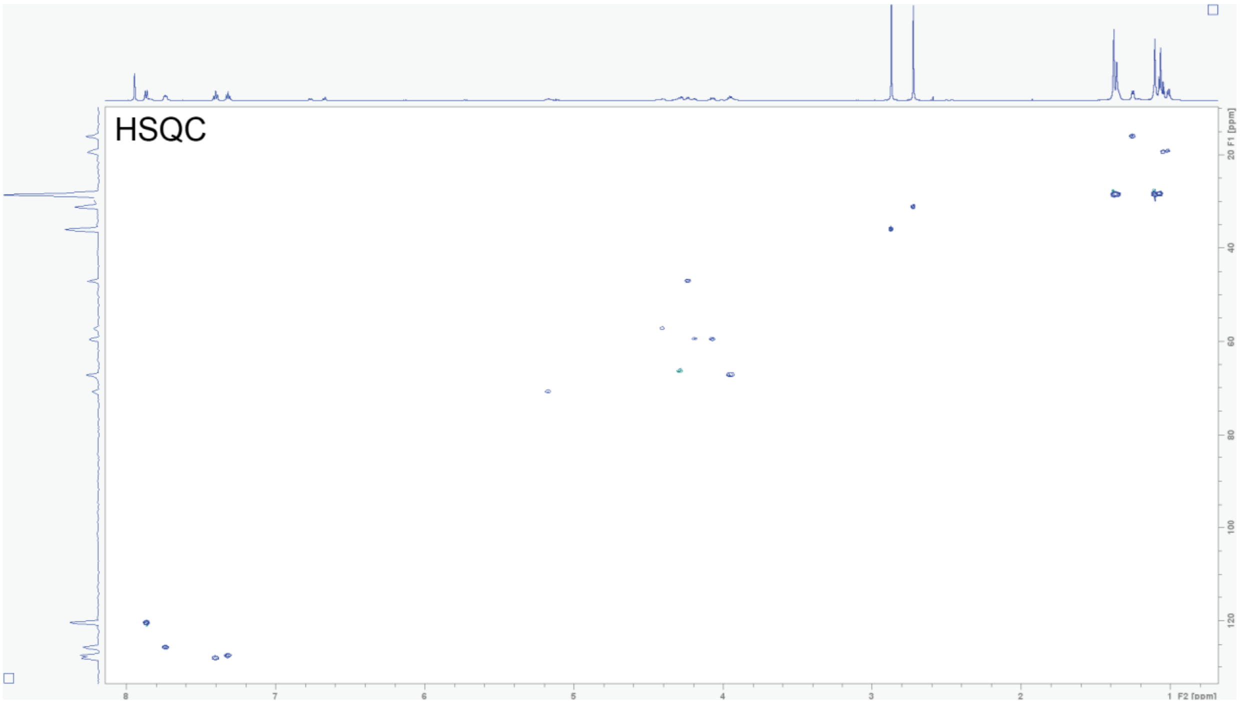


**$^1$H-$^{13}$C HMBC NMR spectrum of N-Fmoc-D-allo-Thr(N-Boc-L-allo-Thr(tBu))-OH (4) in DMSO-*d*$_6$.**

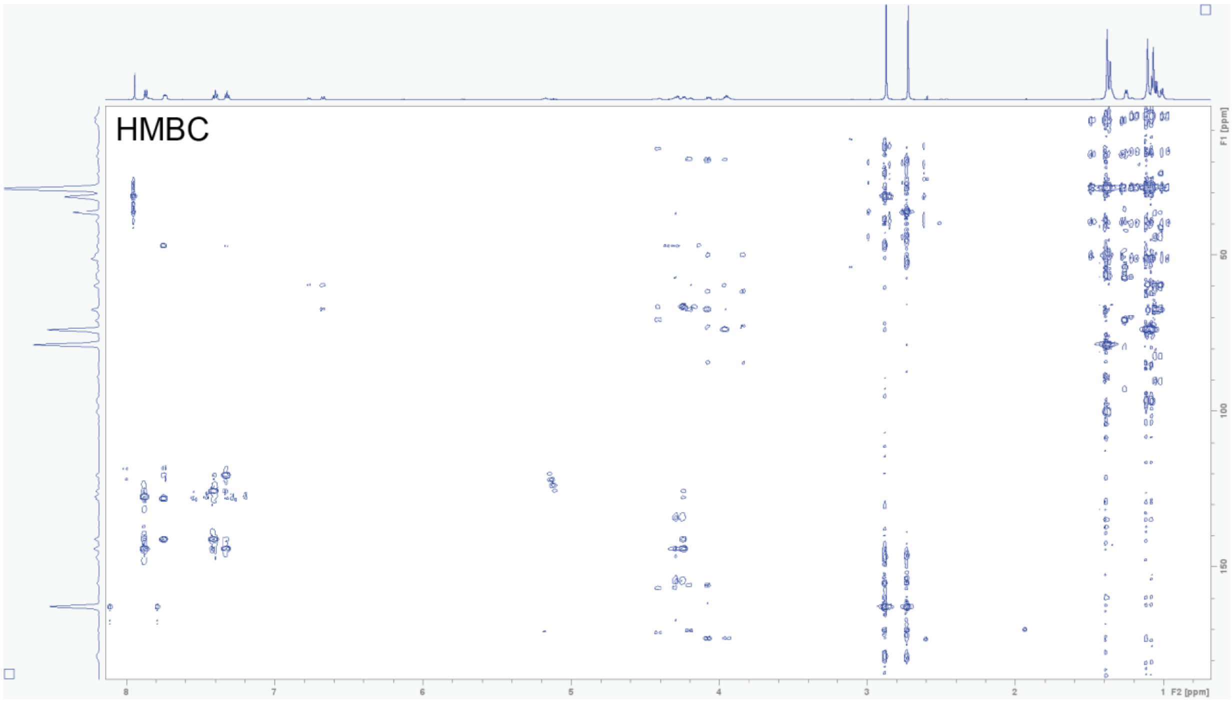

**Table S13. NMR spectroscopic data for 4-cyano-L-tryptophan imacidin (5)(600 MHz in DMSO-$d_6$)[a]**

| Position | $\delta_H$ mult. | $\delta_C$ | Position | $\delta_H$ mult. | $\delta_C$ |
|---|---|---|---|---|---|
| 1 | 0.85 (t, 7) | 14.1 | 38 | 3.05, 3.21 (br) | 26.2 |
| 2 | 1.25 (ov) | 22.2 | 39 | - | - |
| 3 | 1.21 (ov) | 29.2 | 40 | - | - |
| 4 | 1.21 (ov) | 29.2 | 41 | - | - |
| 5 | 1.21 (ov) | 29.2 | 42 | - | - |
| 6 | 1.21 (ov) | 29.2 | 43 | - | - |
| 7 | 1.21 (ov) | 29.2 | 44 | 7.88 (d, 8) | - |
| 8 | 1.21 (ov) | 29.2 | 45 | 4.23 (br) | 48.7 |
| 9 | 1.30, 1.33 (ov) | 37.0 | 46 | 1.31 (d, 7.5) | 17.1 |
| 10 | 3.73 (br) | 68.1 | 47 | - | - |
| 11 | 2.13, 2.21 (ov) | 43.9 | 48 | 8.55 (br) | - |
| 12 | - | - | 49 | 4.13 (t, 7) | 59.0 |
| 13 | 8.32 (ov) | - | 50 | 4.97 (m, 6) | 71.0 |
| 14 | 4.55 (br) | 53.1 | 51 | 1.18 (d, 5.5) | 17.3 |
| 15 | 2.76, 2.94 (br) | 52.5 | 52 | - | - |
| 16 | - | - | 53 | - | - |
| 17 | 7.97 (br) | - | 54 | 4.12 (t, 7) | 58.7 |
| 18 | 4.52 (br) | 53.0 | 55 | 3.79 (br) | 66.6 |
| 19 | 3.10, 3.63 (br) | 26.2 | 56 | 0.89 (br) | 19.7 |
| 20 | - | - | 57 | - | - |
| 21 | 7.31 (s) | 127.4 | 58 | 7.95 (br) | - |
| 22 | - | - | 59 | 4.45 (br) | 53.1 |
| 23 | - | - | 60 | 2.85, 2.90 (br) | 29.3 |
| 24 | 7.44 (d, 7.5) | 125.7 | 61 | - | - |
| 25 | 7.17 (d, 7.8) | 121.0 | 62 | - | - |
| 26 | 7.66 (d, 8) | 117.2 | 63 | - | - |
| 27 | - | - | 64 | - | - |
| 28 | - | - | 65 | - | - |
| 29 | - | - | 66 | 8.23 (ov) | - |
| 30 | - | - | 67 | 4.18 (br) | 48.9 |
| 31 | 8.30 (ov) | - | 68 | 1.24 (ov) | 17.7 |
| 32 | 2.46, 2.94 (br) | 38.5 | 69 | - | - |
| 33 | 1.55, 1.57 (br) | 25.5 | 70 | 7.48 (ov) | - |
| 34 | 2.08, 2.11 (br) | 32.9 | 71 | 4.30 (t, 7.8) | 58.4 |
| 35 | - | - | 72 | 4.22 (m, 6.5) | 66.6 |
| 36 | 9.05 (br) | - | 73 | 1.05 (d, 6.5) | 19.6 |
| 37 | 3.83 (br) | 55.5 | 74 | - | - |

[a]Chemical shift  and (multiplicity, *J* in Hz).

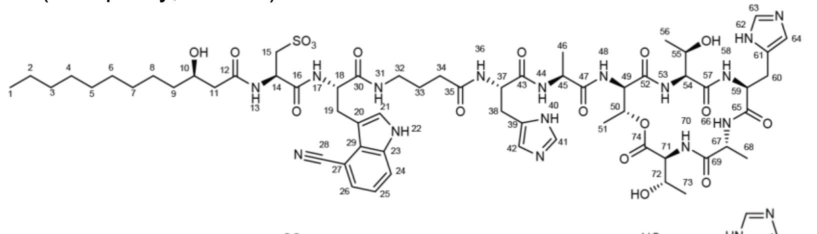

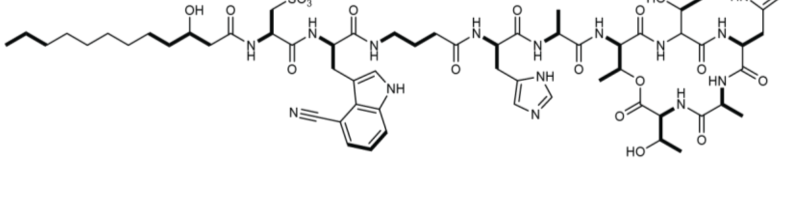

**$^1$H NMR spectrum of 4-cyano-L-tryptophan imacidin (5) in DMSO-$d_6$.**

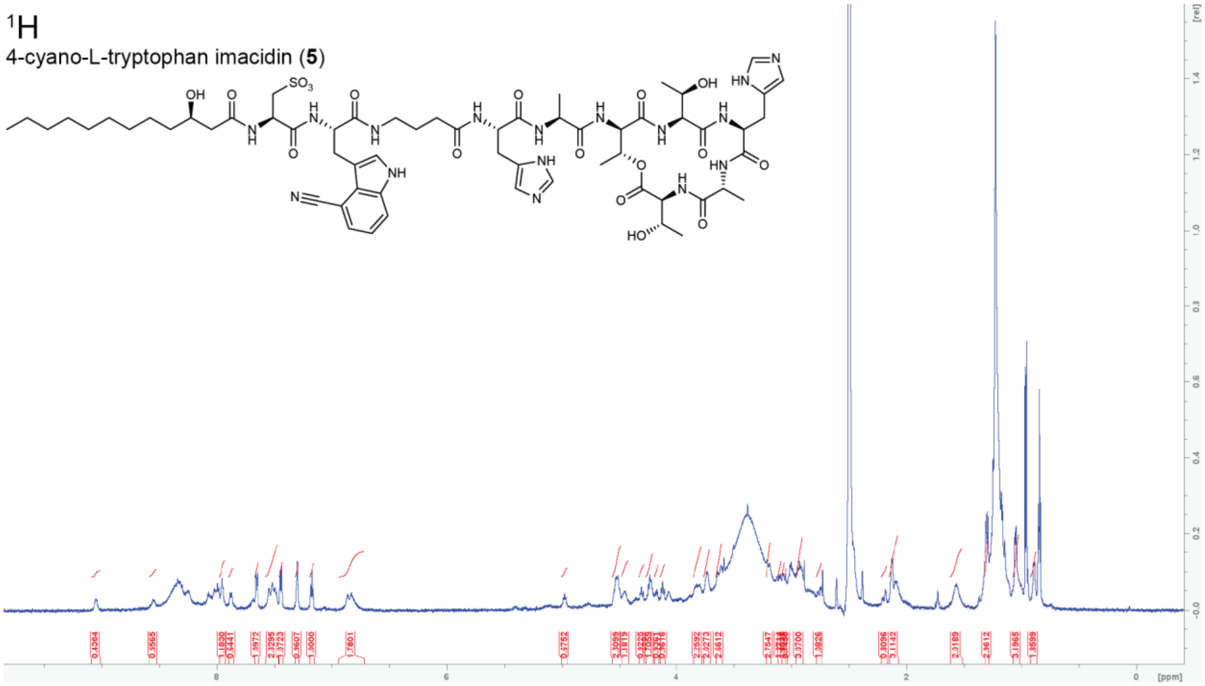


**$^1$H-$^1$H COSY NMR spectrum of 4-cyano-L-tryptophan imacidin (5) in DMSO-$d_6$.**

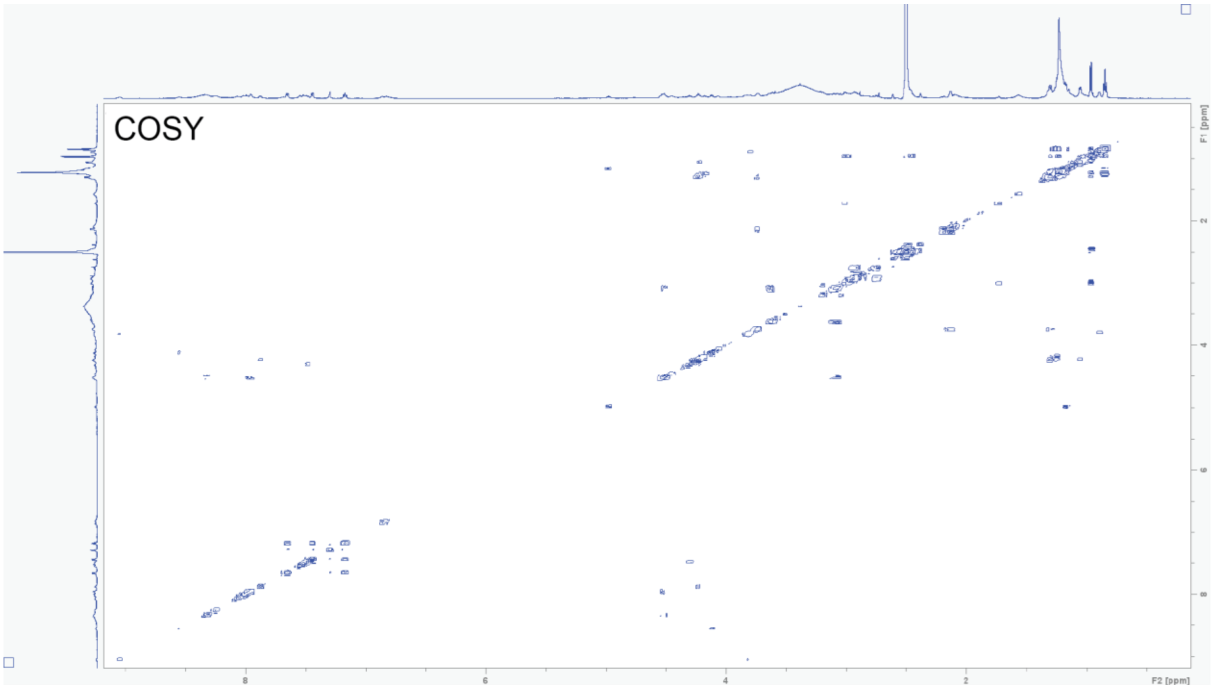

**[1]H-[13]C HSQC NMR spectrum of 4-cyano-L-tryptophan imacidin (5) in DMSO-*d*6.**

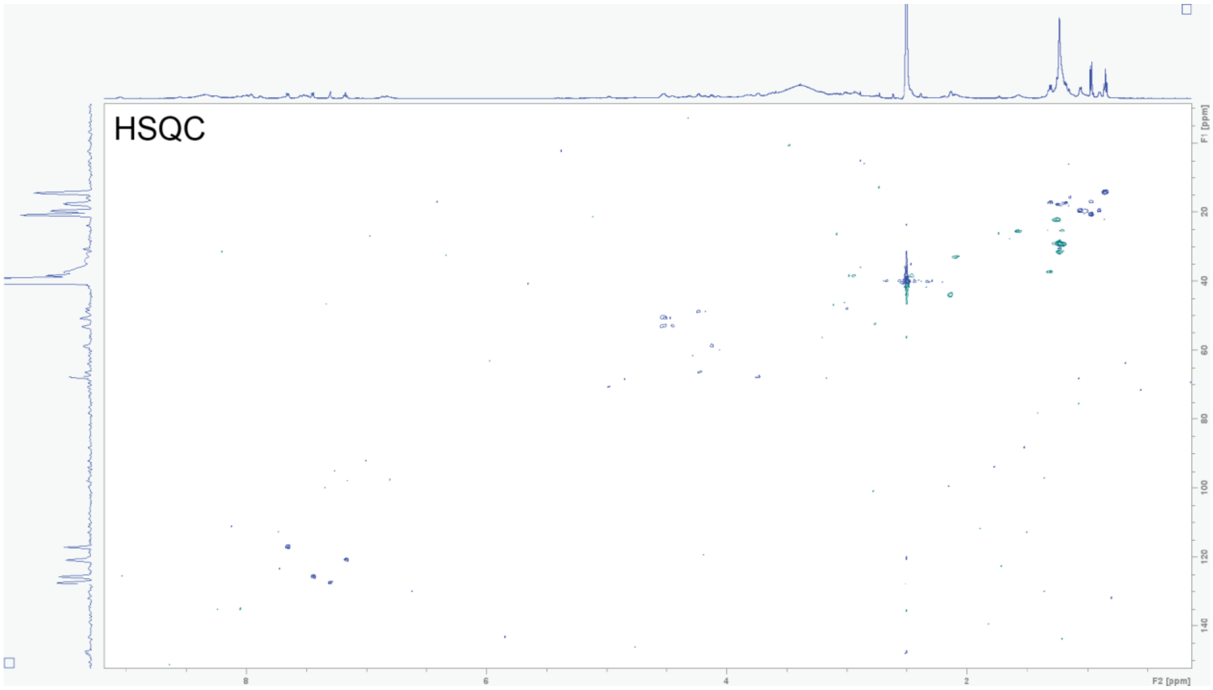


**Table S14. NMR spectroscopic data for N-Fmoc-N-acetyl-L-2,4-diaminobutyrate (6)(600 MHz in DMSO-*d*6)[a]**

| Position | $\delta_H$ mult. | $\delta_C$ | Position | $\delta_H$ mult. | $\delta_C$ |
|---|---|---|---|---|---|
| 1 | 1.75 (s) | 22.2 | 10 | 4.23, 4.24 (ov) | 65.2 |
| 2 | - | 168.8 | 11 | 4.19 (t, 7) | 46.2 |
| 3 | 7.84 (ov) | - | 12 | - | 143.4 |
| 4 | 3.03, 3.10 (m, 6.5) | 35.4 | 13 | 7.69 (d, 7.5) | 124.9 |
| 5 | 1.70, 1.85 (m, 6.5) | 30.3 | 14 | 7.29 (t, 7.5) | 126.7 |
| 6 | 3.94 (m, 4.5) | 51.5 | 15 | 7.37 (t, 7.5) | 127.2 |
| 7 | - | 173.4 | 16 | 7.85 (d, 7.5) | 119.7 |
| 8 | 7.59 (d, 8) | - | 17 | - | 140.3 |
| 9 | - | 155.7 | | | |

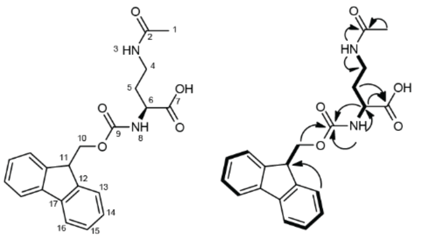

**$^1$H NMR spectrum of N-Fmoc-N-acetyl-L-2,4-diaminobutyrate (6) in DMSO-$d_6$.**

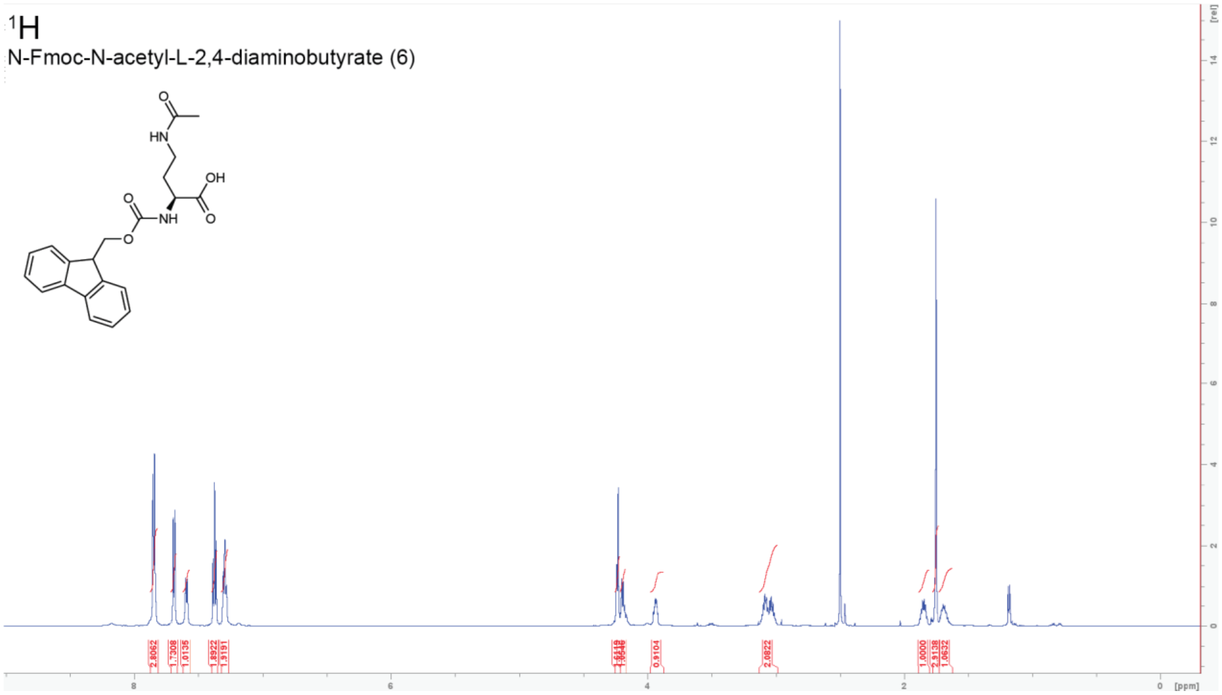


**$^1$H-$^1$H COSY NMR spectrum of N-Fmoc-N-acetyl-L-2,4-diaminobutyrate (6) in DMSO-$d_6$.**

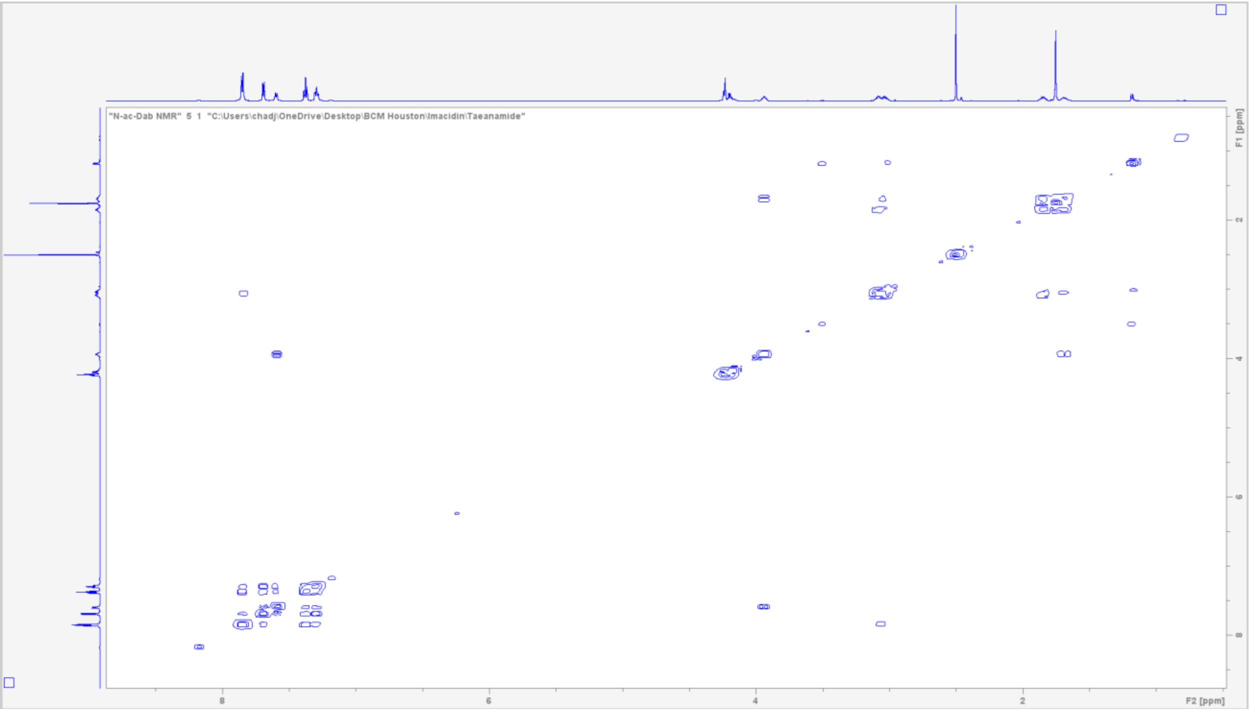

**$^{1}H$-$^{13}C$ HSQC NMR spectrum of N-Fmoc-N-acetyl-L-2,4-diaminobutyrate (6) in DMSO-$d_6$.**

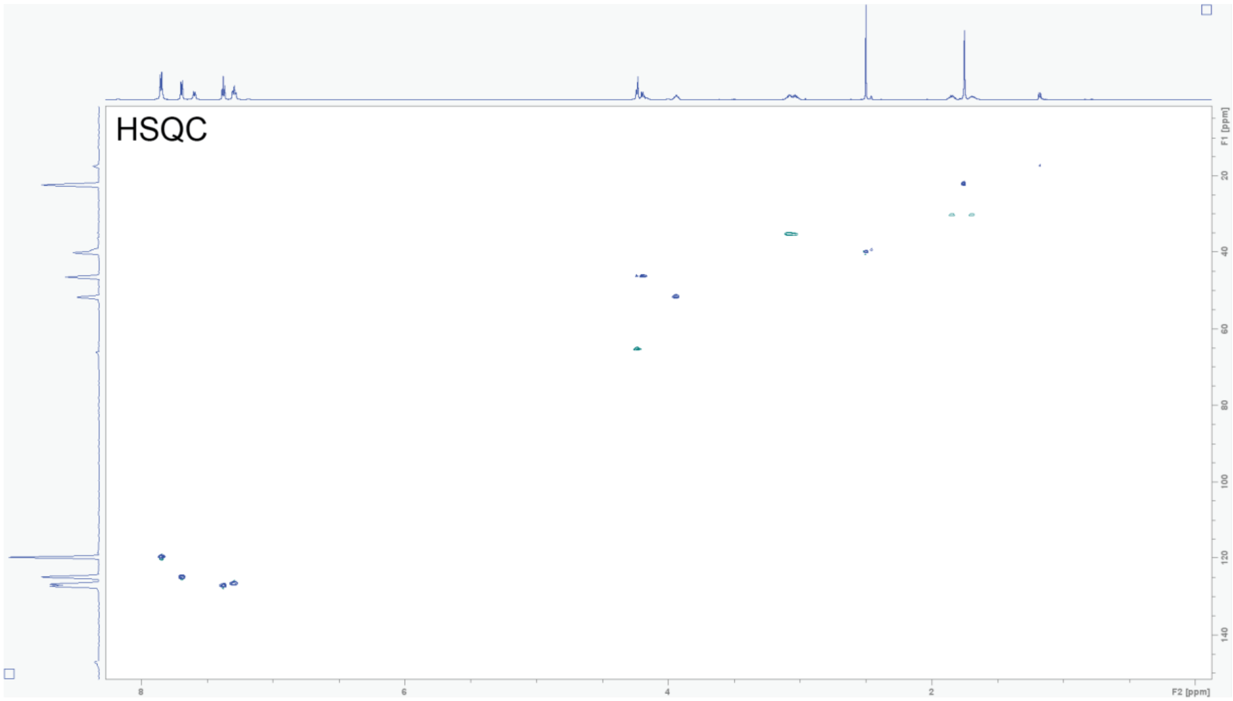


**$^{1}H$-$^{13}C$ HMBC NMR spectrum of N-Fmoc-N-acetyl-L-2,4-diaminobutyrate (6) in DMSO-$d_6$.**

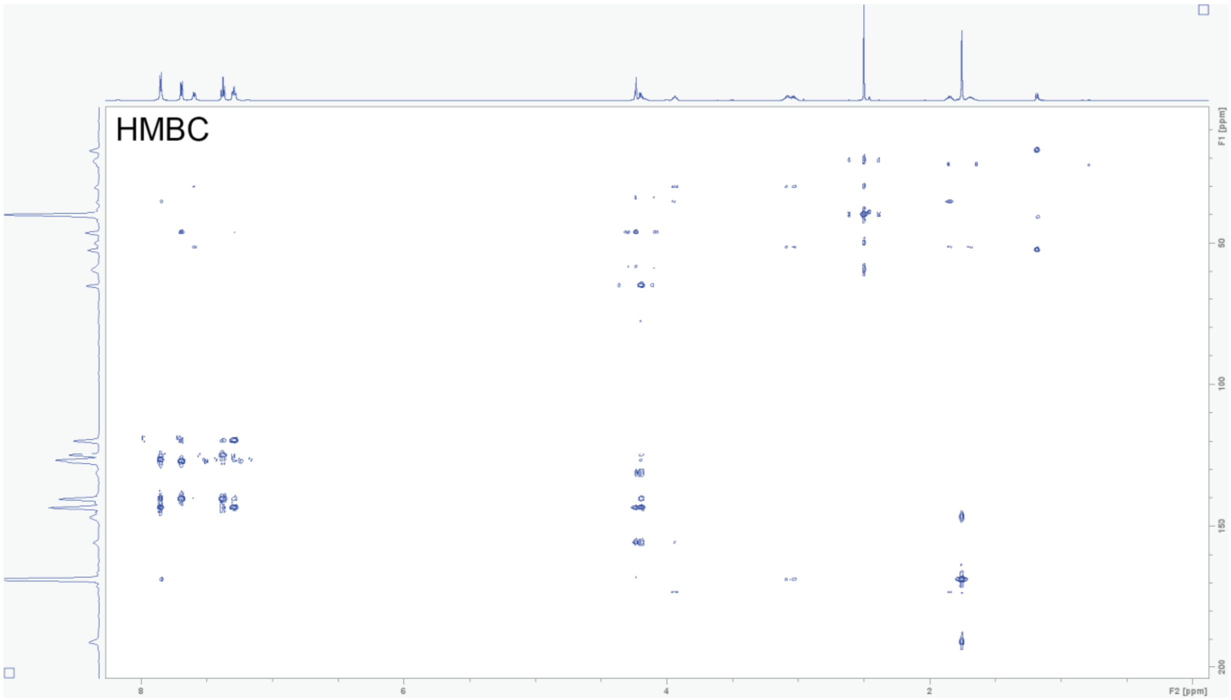

**Table S15. High Resolution mass spectrometry of compounds described in this study.**

| Compound | Molecular Formula | *m/z* Calculated | *m/z* Observed | Error (ppm) |
|---|---|---|---|---|
| Imacidin-cys 1 | $C_{55}H_{90}N_{14}O_{18}S$ | 1267.6351 | 1267.6324 | -2.09 |
| Imacidin-cys 2 | $C_{56}H_{92}N_{14}O_{18}S$ | 1281.6507 | 1281.6488 | -1.52 |
| Imacidin-cys 3 | $C_{57}H_{94}N_{14}O_{18}S$ | 1295.6664 | 1295.6644 | -1.52 |
| Imacidin-glu 1 | $C_{57}H_{92}N_{14}O_{17}$ | 1245.6837 | 1245.6836 | -0.08 |
| Imacidin-glu 2 | $C_{58}H_{94}N_{14}O_{17}$ | 1259.6994 | 1259.6993 | -0.07 |
| Imacidin-asp 1 | $C_{56}H_{90}N_{14}O_{17}$ | 1231.6681 | 1231.6678 | -0.22 |
| Imacidin-asp 2 | $C_{57}H_{92}N_{14}O_{17}$ | 1245.6837 | 1245.6826 | -1.15 |
| IMC synthetic | $C_{57}H_{94}N_{14}O_{18}S$ | 1295.6664 | 1295.6647 | -1.31 |
| Taeanamide C | $C_{51}H_{87}N_{11}O_{16}$ | 1110.6405 | 1110.6388 | -1.54 |
| TAE synthetic | $C_{51}H_{87}N_{11}O_{16}$ | 1110.6405 | 1110.6388 | -1.54 |
| Boc-L-*allo*-Thr(tBu)-OH | $C_{13}H_{25}NO_5$ | 276.1805 | 276.1801 | -1.46 |
| Fmoc-D-*allo*-Thr(Boc-L-*allo*-Thr(tBu))-OH | $C_{32}H_{42}N_2O_9$ | 599.2963 | 599.2961 | -0.27 |
| Fmoc-4-*N*-acetyl-L-2,4-diaminobutyric acid | $C_{21}H_{22}N_2O_5$ | 383.1601 | 383.1598 | -0.93 |
| 4-cyano-L-tryptophan imacidin | $C_{63}H_{92}N_{16}O_{18}S$ | 1393.6569 | 1393.6539 | -2.13 |

**Supplementary note.** Early designs for fluorescent imacidin derivatives used amide coupling of the commercial fluorophore Cy5.5 amine (Lumiprobe) to the carboxylic acid found in abundant aspartate and glutamate congeners (Supplementary Fig. 23). This yielded fluorescent derivatives that lacked antibacterial activity against *S. coelicolor* M1154. As an alternative, we incorporated 4-cyano-L-tryptophan instead of L-leucine during SPPS, resulting in a minimally-altered blue-fluorescent imacidin derivative (Supplementary Fig. 24, Supplementary Table 13). This derivative retained partial activity against M1154 (Supplementary Fig. 25; MIC: 2.5 μg ml$^{-1}$). Unfortunately, while this derivative fluoresced in buffer, it requires an aqueous micro-environment for fluorescence and displays spectral overlap with protein tryptophans, complicating analysis of binding to the interior of the membrane protein MurJ.

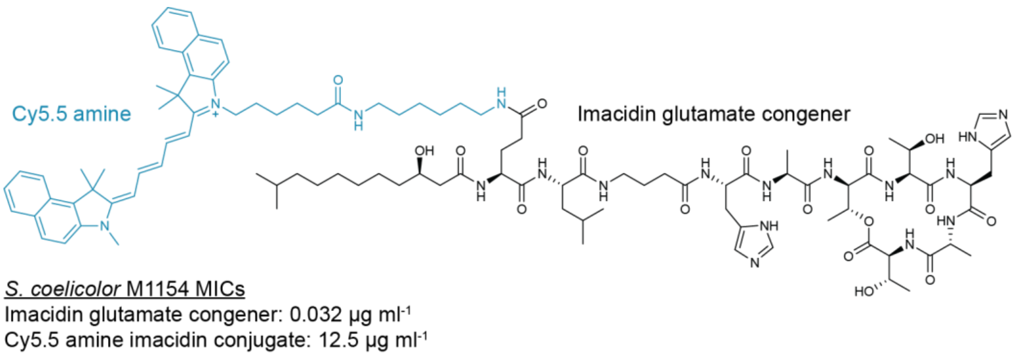


**Figure S23. Cy5.5 Conjugate Structure and MICs.** To probe interactions with the MurJ protein we attempted to create a fluorescent version of imacidin using amide coupling to attach a Cy5.5 amine to the glutamate residue on the lipid tail. The addition of Cy5.5 caused imacidin activity to significantly decrease rendering it ineffective for further studies.

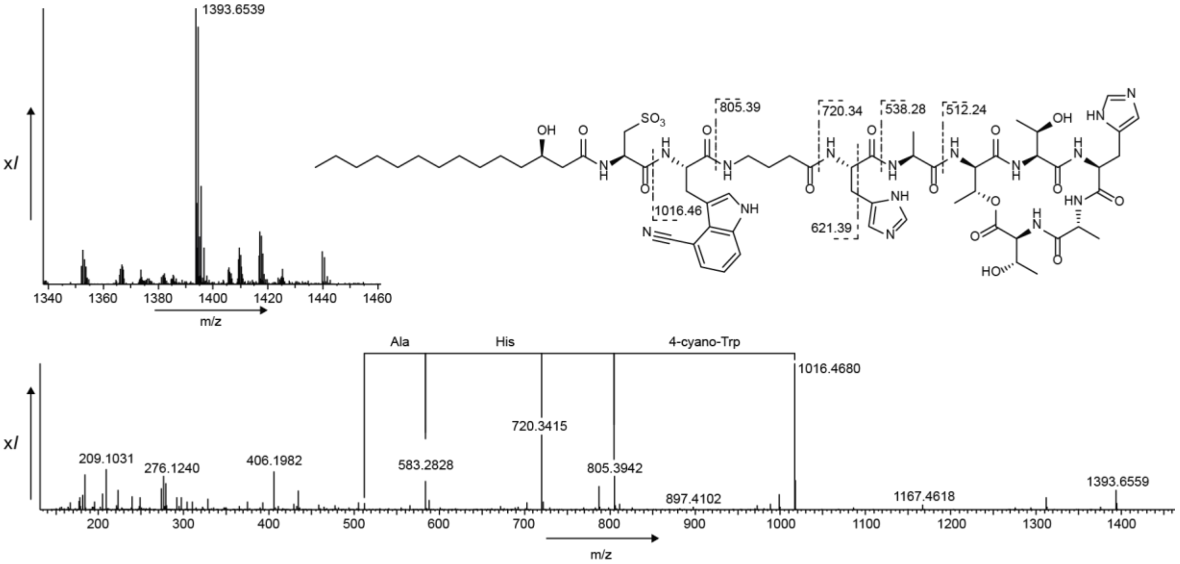


**Figure S24. High resolution mass spectrometry data for 4-cyano-tryptophan imacidin.** Chemical structure and parent ion of *m/z* [M+H]$^{+}$ 1393.65 along with MS/MS spectra.

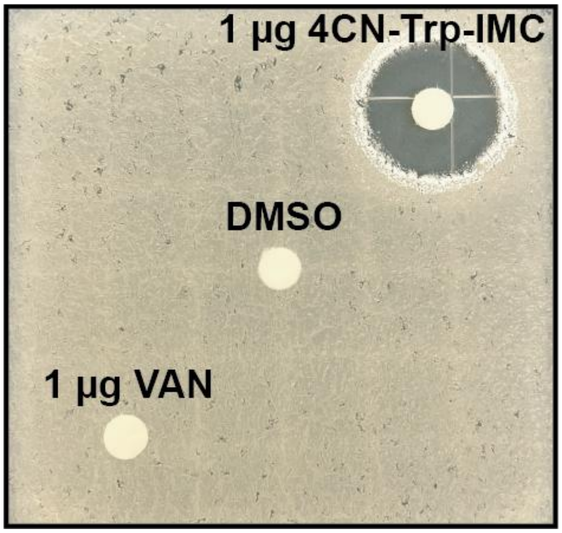


**Figure S25. 4-cyano-L-tryptophan imacidin antibacterial activity.** As an alternative to using Cy5.5 to create fluorescent imacidin derivatives, we incorporated a 4-cyano-L-tryptophan in the place of L-leucine during solid-phase peptide synthesis. The 4-cyano-L-tryptophan imacidin derivative retained partial activity against *S. coelicolor* M1154 in an agar disk diffusion assay with MICs showing inhibition at 2.5 µg $mL^{-1}$.

### References


1. Tobias Keiser, Mervyn J. Bibb, Mark J. Buttner, Keith F. Chater, & David A. Hopwood. *Practical Streptomyces Genetics.* (The John Innes Foundation, 2000).
2. Imai, Y. *et al.* A new antibiotic selectively kills Gram-negative pathogens. *Nature* **576**, 459–464 (2019).
3. Johnston, C. W. *et al.* Assembly and clustering of natural antibiotics guides target identification. *Nat. Chem. Biol.* **12**, 233–239 (2016).
4. De Coster, W. & Rademakers, R. NanoPack2: population-scale evaluation of long-read sequencing data. *Bioinformatics* **39**, btad311 (2023).
5. Chen, S., Zhou, Y., Chen, Y. & Gu, J. fastp: an ultra-fast all-in-one FASTQ preprocessor. *Bioinformatics* **34**, i884–i890 (2018).
6. Chen, S. Ultrafast one-pass FASTQ data preprocessing, quality control, and deduplication using fastp. *iMeta* **2**, e107 (2023).
7. Wick, R. R. *et al.* Trycycler: consensus long-read assemblies for bacterial genomes. *Genome Biol.* **22**, 266 (2021).
8. Bouras, G. *et al.* How low can you go? Short-read polishing of Oxford Nanopore bacterial genome assemblies. *Microbial Genomics* vol. 10 (2024).
9. Chklovski, A., Parks, D. H., Woodcroft, B. J. & Tyson, G. W. CheckM2: a rapid, scalable and accurate tool for assessing microbial genome quality using machine learning. *Nat. Methods* **20**, 1203–1212 (2023).
10. Barrick, J. E. *et al.* Genome evolution and adaptation in a long-term experiment with Escherichia coli. *Nature* **461**, 1243–1247 (2009).
11. Abramson, J. *et al.* Accurate structure prediction of biomolecular interactions with AlphaFold 3. *Nature* **630**, 493–500 (2024).
12. Mirdita, M. *et al.* ColabFold: making protein folding accessible to all. *Nat. Methods* **19**, 679–682 (2022).
13. Altschul, S. F., Gish, W., Miller, W., Myers, E. W. & Lipman, D. J. Basic local alignment search tool. *J. Mol. Biol.* **215**, 403–410 (1990).
14. Schindelin, J. *et al.* Fiji: an open-source platform for biological-image analysis. *Nat. Methods* **9**, 676–682 (2012).
15. Kuk, A. C. Y., Mashalidis, E. H. & Lee, S.-Y. Crystal structure of the MOP flippase MurJ in an inward-facing conformation. *Nat. Struct. Mol. Biol.* **24**, 171–176 (2017).
16. Kohga, H. *et al.* Crystal structure of the lipid flippase MurJ in a 'squeezed' form distinct from its inward- and outward-facing forms. *Struct. Lond. Engl. 1993* **30**, 1088-1097.e3 (2022).
17. Bolla, J. R. *et al.* Direct observation of the influence of cardiolipin and antibiotics on lipid II binding to MurJ. *Nat. Chem.* **10**, 363–371 (2018).
18. Zheng, S. *et al.* Structure and mutagenic analysis of the lipid II flippase MurJ from Escherichia coli. *Proc. Natl. Acad. Sci. U. S. A.* **115**, 6709–6714 (2018).
19. Kumar, S., Rubino, F. A., Mendoza, A. G. & Ruiz, N. The bacterial lipid II flippase MurJ functions by an alternating-access mechanism. *J. Biol. Chem.* **294**, 981–990 (2019).
20. Helbling, R. E., Aeschimann, W., Simona, F., Stocker, A. & Cascella, M. Engineering Tocopherol Selectivity in α-TTP: A Combined In Vitro/In Silico Study. *PLOS ONE* **7**, e49195 (2012).
21. Hofmann, L., Gulati, S., Sears, A., Stewart, P. L. & Palczewski, K. An effective thiol-reactive probe for differential scanning fluorimetry with a standard real-time polymerase chain reaction device. *Anal. Biochem.* **499**, 63–65 (2016).
22. Gomez-Escribano, J. P. & Bibb, M. J. Engineering Streptomyces coelicolor for heterologous expression of secondary metabolite gene clusters. *Microb. Biotechnol.* **4**, 207–215 (2011).
23. Huang, J. *et al.* Cross-regulation among disparate antibiotic biosynthetic pathways of Streptomyces coelicolor. *Mol. Microbiol.* **58**, 1276–1287 (2005).
24. Hong, H.-J., Hutchings, M. I., Hill, L. M. & Buttner, M. J. The Role of the Novel Fem Protein VanK in Vancomycin Resistance in Streptomyces coelicolor*. *J. Biol. Chem.* **280**, 13055–13061 (2005).
25. Love, J. *et al.* The New York Consortium on Membrane Protein Structure (NYCOMPS): a high-throughput platform for structural genomics of integral membrane proteins. *J. Struct. Funct. Genomics* **11**, 191–199 (2010).
